# Tracing the epistemic arc: Distinct physiological signatures for curiosity, insight, understanding and liking when viewing visual art

**DOI:** 10.1101/2025.05.15.654230

**Authors:** Dominik Welke, Edward A. Vessel

## Abstract

A series of internal states, proceeding from curiosity (drive state) to insight (uncertainty reduction) to pleasure (reward and reinforcement), might represent a fundamental pathway for motivated learning. Here we present a paradigm that combines measurement of curiosity, insight (*aha*), understanding and liking to outline the contours of this epistemic arc and trace their neuronal correlates. In two preregistered experiments we employ the “title effect” - the fact that additional semantic information can change how one understands and enjoys a piece of art. Participants viewed paintings and rated their curiosity for seeing the title. This was followed either by the original title or a dummy title lacking additional information, and a second presentation of the image, after which participants rated their strength of *aha*, aesthetic liking, and feeling of understanding the artwork. In a behavioral study (N=55 participants) we establish empirically that visual art with their titles can prompt strong feelings of *aha*. The association between ratings is complex and shaped by the type and amount of information in the stimuli; e.g. *aha* is linearly linked to liking and nonlinearly to understanding. In an EEG, ECG and eye tracking study with the same paradigm (N=49 participants) we use time-resolved decoding to trace the temporal sequence of processing. Understanding can be decoded from various (neuro)physiological correlates, over extended timespans. Robust decoding of curiosity both before and after the titles substantiates its relevance within the epistemic arc. *Aha* could be decoded from pupil size, but not from the EEG. Lastly, a fast initial EEG response was linked to aesthetic liking. Though much remains to explore, our study provides a new window into core behavioral and physiological properties of motivated information foraging.

## 1 Introduction

We are constantly confronted with novel, potentially ambiguous stimuli, and trying to make sense of this incoming information (i.e. forming coherent, meaningful representations) is a cornerstone of human cognition. Often, sense-making involves insight – a sudden jump in understanding such as when the solution to a previously unsolved problem becomes clear, or when a new piece of information allows for better understanding (Sternberg & Davidson, 1995). Salient examples can be found in the visual domain, where our perception of an ambiguous stimulus can suddenly change and allow us to perceive structure (such as with degraded two-tone “Mooney” images; Mooney & Ferguson, 1951).

Insight can be characterized as a rapid decrease in uncertainty, or gain in information. Recent work has linked insight to the preceding state of curiosity (Van de Cruys et al., 2021): the more curiosity, the greater the *aha* – a measure of the phenomenal feeling that accompanies insight. This finding can be used to place insight within a motivational framework: curiosity is a drive state to reduce uncertainty, improved understanding is the desired outcome, and insight (or *aha*) marks moments of particularly effective uncertainty reduction. Based on their findings Van de Cruys et al. (2021) hypothesize that *aha* might act as a meta-cognitive feedback signal that can help us optimize learning efficiency and allocate limited cognitive resources to solvable tasks (see also Laukkonen et al., 2023). If insight triggers a motivational cascade, it should further engage reward processes and be associated with pleasure – the “pleasure from understanding” (Vessel, n.d.). Indeed, some studies provide empirical evidence that *aha* can feel pleasurable (Skaar & Reber, 2020; Shen et al., 2016) and is linked to the reward system (Cristofori et al., 2018; Oh et al., 2020).

We hypothesize that this series of processes, from curiosity to insight to pleasure and reward, might represent a fundamental pathway for motivated learning – referred to as the “epistemic arc” (Van de Cruys et al., 2024). Here, we present a paradigm to study meaning-making from complex naturalistic stimuli. We combine the controlled delivery of novel information (via our stimuli) with measurements of curiosity, insight (*aha*), understanding and liking, to outline the contours of this epistemic arc and study their temporal sequence and neuronal mechanisms.

Van de Cruys et al. (2021) investigated insight and information processing using a relatively low-level task of perceptual understanding in the visual domain – namely Mooney Images (Mooney & Ferguson, 1951), black-and-white two tone images derived from photographs that are initially hard to interpret, but that become readily interpretable after seeing the original photograph. While this method is very effective, it is clear that similar mechanisms can act higher up the cognitive hierarchy. For example, insight might be gained by combining information from different sensory modalities (e.g. merging visual input and language cues when watching the news) or by contextualizing novel stimuli with prior knowledge or personal memories. In this study we use visual art and employ a phenomenon called the “title effect” – the fact that text based, semantic information supplementing an artwork (such as, but not only, titles) can affect how an observer understands, interprets or judges a piece of visual art. For Mooney images it has been shown that the strength of *aha* with a particular image also predicts subsequent ratings of aesthetic liking (Muth & Carbon, 2013). We expect that this effect will also extend to visual art. In fact, insight may even be a key factor contributing to the feelings of delight or being moved that many people can derive from savoring art.

### 1.1 The “title effect” – a case of aesthetic *aha*?

Several psychological studies have investigated the effects of titles on different ratings of visual art. While all studies we found reported significant effects of the applied title manipulation on one or more dependent measures, the findings are not entirely conclusive.

The clearest picture emerges regarding ratings of felt understanding or meaningfulness of the artwork: a total of 10 experiments in 6 studies found a significant effect of title manipulation (Russell & Milne, 1997; Russell, 2003; Millis, 2001; Leder et al., 2006; Mullennix et al., 2018; Szubielska et al., 2021), while only 2 experiments in 2 studies did not. Yet even in the studies reporting overall non-significant results, differences between titled and untitled condition were present (Russell, 2003, Exp. 1; Bubić et al., 2017). Taken together, these studies provide convincing evidence that additional semantic information given in text form changes observer’s subjective understanding of paintings.

When looking at aesthetic preference ratings (e.g. liking, being moved, etc.) the findings are less clear: several studies reported significantly higher ratings for art presented with a title than without a title (Millis, 2001; Leder et al., 2006; Belke et al., 2010; Hristova et al., 2011; Gerger & Leder, 2015; Mastandrea & Umiltà, 2016; Bubić et al., 2017). However, in many cases these effects only occurred for specific types of titles: for elaborative titles, but not descriptive or random titles (Millis, 2001; Leder et al., 2006), for semantically related, but not unrelated titles (Belke et al., 2010; Gerger & Leder, 2015), or for fabricated titles emphasizing a specific topic of the work, but not the original titles or titles de-emphasizing this concept (Mastandrea & Umiltà, 2016). Furthermore, the category of artwork might matter: one study explicitly observed a difference between matching and non-matching titles only for abstract art, but not for semi-abstract or figurative art (Gerger & Leder, 2015), while most other studies were restricted to very homogeneous stimulus sets and could hence not help clarify this issue. Several other studies, though, reported null results under seemingly very similar conditions: comparing titles to no titles (Russell & Milne, 1997; Millis, 2001; Russell, 2003), descriptive to more elaborate titles (Millis, 2001; Leder et al., 2006; Mullennix et al., 2018), matching to non-matching titles (Szubielska et al., 2021), or titles alone to longer text descriptions (Russell, 2003).

While none of these studies collected ratings of *aha* or insight – an effect to be established in the current study – previous work has suggested a link between insight and aesthetic liking for lower-level perceptual reveals: the strength of *aha* with Mooney images positively predicted ratings of aesthetic preference, which Muth and Carbon (2013) termed the “aesthetic aha effect”.

### 1.2 Linking the epistemic arc to physiological correlates of insight and aesthetic preference

In order to follow the sequence of the hypothesized epistemic arc we seek to track its components and their temporal profile. To do so, we record and analyze multimodal (neuro)physiological data together with the participants’ responses.

Several studies have tried to identify neurophysiological correlates of insight or aesthetic preferences, respectively. Previous research on the neuroscience of insight has often focused on hemispheric differences, a hypothesis compatible with a set of neuroanatomical and cytoarchitectonic asymmetries in potentially relevant brain regions (especially in the context of language processing; for a review see Kounios & Beeman, 2014). However, there is a small EEG literature on insight demonstrating relatively consistent findings – namely bursts of gamma-band activity preceding self-reported insight moments in various text-based tasks such as remote associate tasks (Jung-Beeman et al., 2004; Sandkühler & Bhattacharya, 2008), verbal puzzles (Sheth et al., 2009), or anagram solving tasks (Oh et al., 2020). In addition to gamma-band activity, these studies less consistently reported increased alpha-power (Jung-Beeman et al., 2004; Sandkühler & Bhattacharya, 2008, both using remote associate tasks) and increased (Oh et al., 2020) or decreased (Sheth et al., 2009) beta-activity prior to the solution.

These activity patterns coarsely resemble those reported in the small frequency-domain M/EEG literature on aesthetic preferences. Studies reported significant effects in almost all of the canonical frequency-bands, but most consistently in the gamma (Lopez-Persem et al., 2020; Cela-Conde et al., 2004; Munar, Nadal, Rosselló, et al., 2012; Munar, Nadal, Castellanos, et al., 2012; Strijbosch et al., 2021) and alpha range (Munar, Nadal, Castellanos, et al., 2012; Kang et al., 2015; Strijbosch et al., 2021). Furthermore, hemispheric differences in alpha power (frontal alpha asymmetry) have been a particular target of neuroaesthetic investigation (Babiloni et al., 2013; Moon et al., 2013; Cheung et al., 2019), and some studies even used an EEG asymmetry index as a proxy for aesthetic preference without further verification through behavioral ratings (Babiloni et al., 2015; Lee et al., 2017).

Event related potentials (ERPs) have also been investigated in the context of aesthetic preferences. Effects on a variety of well established ERP components have been reported including posterior P2 (Righi et al., 2017; Noguchi & Murota, 2013), anterior P2 (Righi et al., 2017; Ma et al., 2015; Jiang & Cai, 2013; Wang et al., 2012), N2 (Bölte et al., 2017; Righi et al., 2017; Ma et al., 2015; de Tommaso et al., 2008; Oliver-Rodríguez et al., 1999; Carbon et al., 2018; Augustin et al., 2011), P3 (Bölte et al., 2017; Righi et al., 2017; Ma et al., 2015; de Tommaso et al., 2008; Oliver-Rodríguez et al., 1999), and less consistently on P1 (Righi et al., 2017; Bölte et al., 2017) and LPP (Marzi & Viggiano, 2010; Werheid et al., 2007; Schacht et al., 2008). In a previous study, we reported differences in very late potentials (3-5 s after stimulus onset; Strijbosch et al., 2021) though we could not replicate any of the previously listed ERP findings. While the presented EEG findings are not entirely conclusive, they nevertheless suggest that preference related information can be recovered from noninvasive scalp EEG.

Further physiological data modalities, such as eye tracking and electrocardiography (ECG) are interesting candidates as well: heart rate has been linked to aesthetic valuation (Tschacher et al., 2012), pupil size has been investigated as a potential indicator of *aha* experiences (Salvi et al., 2020) and surprise during music listening (Liao et al., 2018), and gaze patterns have been investigated as a behavior of interest in empirical aesthetics (with potential title-effects on gaze patterns/exploration behavior; Bubić et al., 2017; Kapoula et al., 2009; Temme, 1992; Hristova et al., 2011). All these physiological data types are amenable to the multivariate decoding approach applied in this study.

### 1.3 The current study

We designed an extended version of the paradigm by Van de Cruys et al. (2021) and used it in two preregistered experiments: in a first purely behavioral task participants were presented with a piece of visual art and asked to rate how curious they were for the title. This was followed by either the artwork’s original title or a statement that the piece was “untitled” (real or fake title condition randomly assigned and fully balanced across participants). Participants then saw the artwork again and were asked to rate their feeling of *aha*, as well as aesthetic liking, and how much they felt they understood the piece. This allowed us to test if previous findings on *aha* and meaning making would replicate in a new stimulus category (art), and also to functionally link these different evaluation processes that have typically been studied separately. We compiled a diverse set of artwork stimuli with matching titles: the stimuli varied by grade of visual abstraction of the artwork (art style: figurative, semi-abstract, or abstract) and information content of the titles (title style: purely descriptive titles that did not add new information, explanatory titles that aided understanding of the depicted content, or evocative titles that created a link to new, remote concepts). This enabled a finer grained investigation of the effects and interactions of these factors on meaning making and preference. We also included Mooney images to compare the strength of *aha* elicited by visual art and their titles to the feeling of *aha* caused by perceptual shifts. In particular, we preregistered the following hypotheses: there would be stronger *aha* experience under the additional information condition (art with real title or Mooney image) than in the no additional information condition (“untitled” art); curiosity ratings would predict the strength of the *aha* experience; *aha* ratings would predict aesthetic preference ratings; art with their real titles would receive higher aesthetic ratings than untitled art stimuli; art with their real titles would receive higher understanding ratings than untitled art stimuli; there would be an interaction with the degree of abstraction, at least for the understanding rating.

In a second experiment we aimed to investigate the physiological correlates and temporal sequence of reward processing and therefore recorded participants’ EEG, pupillometry, and ECG in addition to the behavioral ratings. EEG (both in time and frequency-domain), pupil dilation, and instantaneous heart rate were analyzed using a time-resolved decoding approach designed to show whether and when rating-related information was represented in the respective recordings (following Gwilliams & King, 2020, see methods). This part of the experiment was more exploratory and we only preregistered a few coarse hypotheses based on previous work: that we would find insight related activity in the gamma band; that there would be a positive linear relation between aesthetic preference and gamma band power; that there would be a nonlinear relation between alpha power and aesthetic preference.

## 2 Methods

In two experiments we presented participants with visual artworks (paintings) followed either by their original titles (adding semantic information) or dummy titles (“untitled“; without new information) that may aid interpretation of the images. The paradigm closely followed the study by Van de Cruys et al. (2021) with Mooney images. While the applied experimental paradigm was largely identical the two experiments differed in important aspects (Exp. 1: purely behavioral data, online study, larger stimulus set; Exp. 2: behavioral and physiological data, lab based study, confined stimulus subset based on ratings from Exp. 1).

The experimental design and our hypotheses were preregistered (see https://doi.org/10.17605/OSF.IO/PAGVH for Experiment 1 and https://doi.org/10.17605/OSF.IO/6KW9N for Experiment 2). Any detail of the final methods that differed from the preregistered plan is clearly described and discussed as such. Details on participants and recording devices are reported following the standards proposed for M/EEG studies by the Organization for Human Brain Mapping (COBIDAS Pernet et al., 2018, 2020).

### 2.1 Experiment 1 – Online behavioral study

#### Participants

##### Sample size estimation

Target sample size was estimated via *a priori* power analysis using G*Power software (Faul et al., 2007, version 3.1). As power analysis for multilevel linear models (see Statistical Analysis) is not straightforward, we powered our study for a similar repeated measures ANOVA with parameters “ANOVA: Repeated measures within factors”, alpha = .05, power = .8, effect size *f* = 0, 1758631 (equivalent to *η*^2^ = 0.03, a small effect size), 1 group, 26 measurements (3×4×2 + 2 design, see below), and conservative expectation of 0 correlation among repeated measures). This resulted in a target sample size of 30. As the study was conducted online and we could not control the environment of the data collection we aimed for a substantially larger sample of N = 50 participants. However, we expected higher quality data in comparison to other online studies as the participant pool consisted of motivated, often regular, participants in studies at our institute.

##### Recruitment

Participants were convenience-sampled from the general public in the Rhine-Main metropolitan region in Germany, via direct mailings to members of an institute hosted participant database (> 2000 members at time of recruitment, open to everybody to subscribe). Slots were assigned on a first come, first serve basis. There were no specific inclusion criteria for participation in this study.

##### Final sample

N = 56 participants enrolled in the study and N = 54 finished data collection. The 2 incomplete datasets (13/56 trials, and 53/56 trials) were still included in the analysis. From these 56, 2 participants reported past or current neurological problems (Stroke, MS) under active medical treatment. 5 other participants reported current or past episodes of mental or psychological disorders. 2 of these 5 indicated current medical treatment. The participants were between 20–82 years old (mean = 40.0 years, std = 17.0 years; with 1 participant not indicating their age). 37 were female, 19 male, and 0 indicated ‘other’. 3 were left handed, 49 right handed, and 3 ambidextrous (based on self report, with 1 missing response). Participants had received between 7–26 years of education (mean = 17.4 years, std = 3.9 years; with 3 participants not providing these data). The sample showed a strong bias towards highly educated people: 54 out of 56 held the German Abitur or a university degree as their highest qualification, and 1 participant did not provide these data. See Supp. Tabs. A.1 and A.2 for full details on participant demographics.

Participants spent on average 50:12 minutes to finish the online data collection (std = 23:35, min: 31:23, 25%: 36:56, 50%: 42:24, 75%: 50:27, max: 2:22:35). For 5 participants, this calculation wasn’t possible due to missing timestamps. All participants received monetary compensation of 10 for finishing the experiment and gave their informed consent prior to participation. The study adhered to the ethical standards of the Declaration of Helsinki and was approved by the local ethics committee (Ethics Council of the Max Planck Society) under the Research Ethics Framework Application for Standard Laboratory Studies. Data were collected from December 2021 through January 2022.

#### Stimuli

Each participant saw a subset of 56 art stimuli and 8 Mooney images from a stimulus set of 120 art stimuli and 20 Mooney images.

##### Art stimuli

All art stimuli were taken from the WikiArt.org database containing thousands of images in the public domain. The set of 120 images used in this study was based on WikiArt metadata and the WikiArt Emotions dataset (Mohammad & Kiritchenko, 2018) by the following procedure: 1) we first selected all paintings from 8 western art styles of the last 2 centuries (i.e. romanticism, realism, impressionism, expressionism, cubism, surrealism, abstract expressionism, lyrical abstraction - according to WikiArt metadata), 2) we then excluded all that contained faces (according to WikiArt Emotions ratings), 3) the remaining artworks were categorized into “abstract”, “semi-abstract” or “figurative” 4) and by their title into “purely descriptive” title, “explanatory” title and “evocative” title (both by the experimenter) 5) finally we selected 10 stimuli for each of the 9 factor combinations aiming for a) including stimuli from many possible art styles and b) high “art: surprise” rating from the WikiArt emotions dataset. In the 3 “explanatory title” subgroups we extracted 20 stimuli instead of 10 per bin, technically leading to 3 additional groups with explanatory titles (with slightly lower pre-rated “art: surprise”). This procedure yielded a total of 120 art stimuli, each accompanied by its original title (as stored in the WikiArt database, translated to German).

##### Mooney images

Mooney images were taken from Van de Cruys et al. 2021, publicly available online. Stimuli used in our study were selected by two criteria: 1) not depicting humans, and 2) the average *aha* rating in Van de Cruys et al. 2021. We took the top 10 *aha* rated stimuli and the lowest 10 *aha* rated stimuli from this study. Each Mooney image is accompanied by a matching gray-scale picture from which it was derived (presented as a “solution” cue analogous to the real titles for the art stimuli).

#### Experimental design

We used a within participant factorial design with repeated measures and 5 different participant groups (based on stimulus subsets, see below).

The design can be formalized as a 3×4×2 + 2 design: participants saw visual art stimuli in 3 degrees of abstraction (figurative, semi-abstract, abstract), and 4 title types (descriptive title, evocative title, explanatory title group 1, explanatory title group 2), in either of 2 title reveal conditions (real title, or “untitled” dummy title). These factors were fully crossed, with 2 trials for each of the 24 factor combinations (48 art stimuli). Additionally 2 groups of Mooney trials were presented (high *aha* rating, or low *aha* rating in Van de Cruys et al., 2021), with 4 trials per participant each (8 Mooney stimuli). This led to a total 56 trials per participant (see below for more details on the stimuli).

##### Task procedure

Participants accessed the online experiment implemented in Labvanced (Scicovery GmbH, Germany) via a personalized link. After givin informed consent, they answered a set of ancillary questionnaires (see below). Instructions for the main experiment were displayed, followed by 5 practice trials (2 Mooney images and 3 art stimuli together with their real titles; these stimuli were not presented again in the main experiment). Then started the main experiment, followed by a short debriefing questionnaire. After finishing, participants were redirected to a separate GDPR compliant Limesurvey server (LimeSurvey GmbH, Germany) to provide their details for monetary compensation.

For the main experiment we presented participants with visual artworks (paintings) followed either by their original titles or dummy titles with no additional information (“untitled”).

For each trial participants went through the following sequence: 1) a visual stimulus for 5 s (art or Mooney image), 2) they rated how curious they were for the title or the Mooney solution (self paced response, using a slider on a continuous scale), 3) an additional semantic cue followed (3s presentation time); in case of artwork stimuli this was either the original title or a “untitled” dummy (condition randomly selected, fully balanced across participants), in case of Mooney images this was the grayscale photograph the image was derived from, and 4) the initial image was presented again (5 s presentation time). They then rated 5) to what extent they experienced *aha*, 6) the aesthetic appeal of the image, and 7) their perceived understanding of the image (all self paced responses, using a slider on a continuous scale). The sequence is visualized in Fig. 1a. In this Exp. 1 the last two rating sliders (aesthetic preference and felt understanding) were presented above each other on the same page.

**Figure 1:**
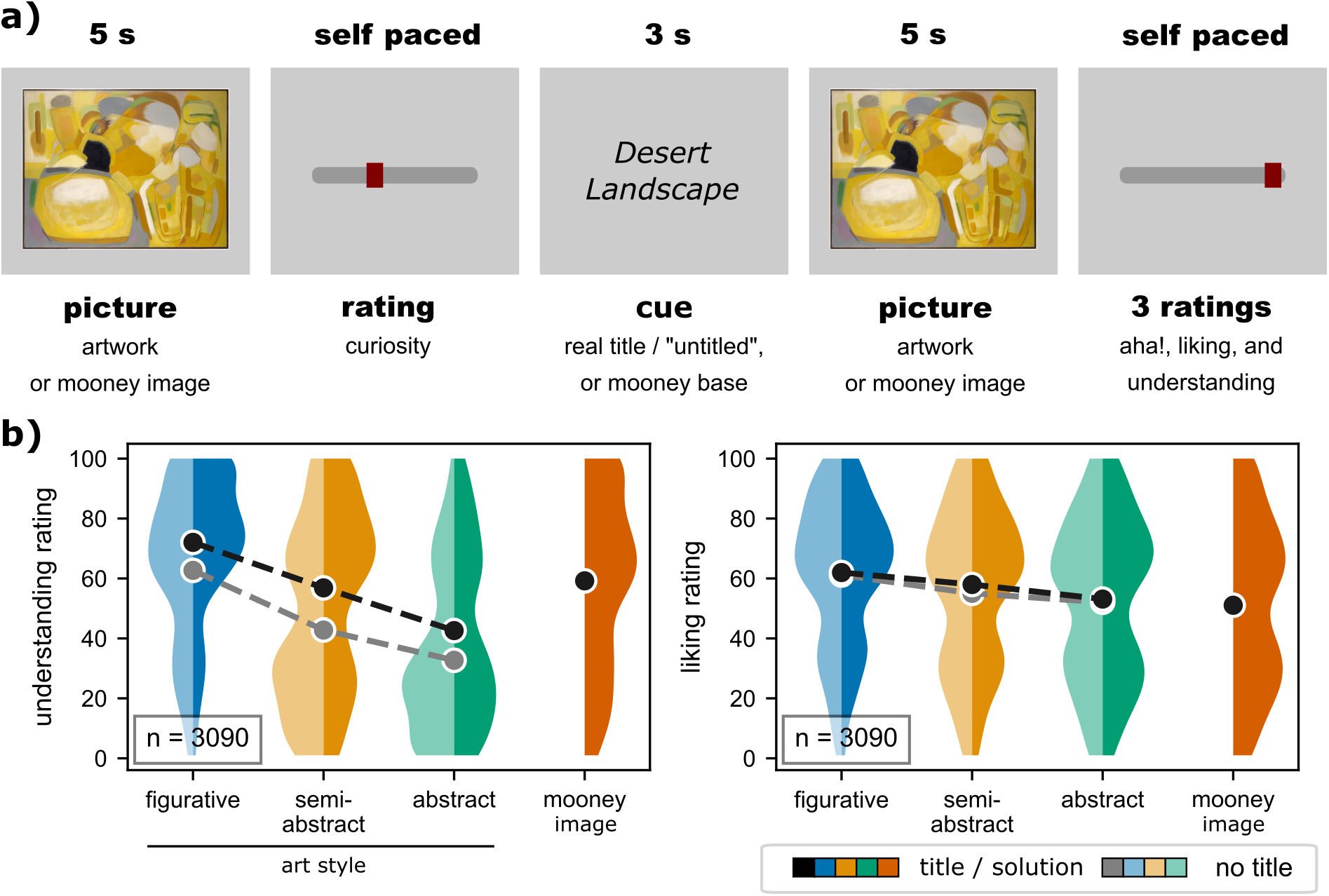
Artworks with their genuine titles evoke higher feeling of understanding than “untitled” art (replicating previous findings). **a) Experimental sequence:** Participants viewed images of visual art and were asked to rate their curiosity for its title. Then followed an artwork title – either the original title or an uninformative dummy (“untitled”). Then they saw the original image again, and were asked to rate their feeling of *aha*, aesthetic liking, and if they felt they understood the piece. Besides visual artworks, a series of Mooney images (Mooney & Ferguson, 1951) were presented (randomly intermixed). Instead of a real or fake title, the base photograph used to create the Mooney image was shown. **b) Replicating a title-effect on understanding, but not on aesthetic liking:** violin plots show the distribution of all ratings (n=3090 across 56 participants) separately for each stimulus category (visual art in 3 levels of figurativeness plus Mooney images), split by title condition (real or untitled; all Mooney images were accompanied by their real base photography, here categorized as titled). We replicated a strong effect of the title manipulation on felt understanding, present in all 3 categories of artwork, but no significant differences for aesthetic liking.

##### Randomization

The assignment of each artwork stimulus to the titled or untitled condition was randomized and fully balanced across participants. The stimulus order for each participant was randomized with fully intermixed trials (not more than 1 stimulus of the same factor combination in a row). Participants were randomly assigned to either of 5 groups - the stimulus subsets in these 5 groups differed but fully overlapped with each other: each individual stimulus was presented in 2 of the 5 groups.

#### Behavioural measures and questionnaires

First, participants answered a set of questionnaires: the big-five personality inventory (BFI-2-XS: Rammstedt et al., 2020; Soto & John, 2017), the short boredom proneness scale (sBPS: Struk et al., 2017), the short form positive negative affect schedule (PANAS-SF: Thompson, 2007), and the Aesthetic Responsiveness Assessment (AReA: Schlotz et al., 2020) in their German versions, as well as a background questionnaire on basic demographic information (including age, gender, education, and information on mood or neurological disorders).

#### Data analysis

Statistical analysis and data visualization was done in Python (version 3.7.11) and R (version 4.1.3) in virtual environments on a Macintosh PC (Apple Inc). Jupyter notebooks reproducing figures and results are publicly available online (see data availability).

##### Linear mixed models

The main hypotheses were tested using linear mixed effects regressions with the *lmer* function from R’s *lme4* package (Bates et al., 2015, version 1.1-30). Significance values for individual effect coefficients were estimated using Satterthwaite’s degrees-of-freedom method implemented in R’s *lmertest* package (Kuznetsova et al., 2017, version 3.1-3). Full LMM tables for Supp. A.2.2 were generated using the *tab_model* function from R’s *sjPlot* package. To assess the overall significance of the reported linear models, Likelihood Ratio Tests were performed: LMM-coefficients were compared to reduced intercept-only models (no fixed effect, but random effects of participant and stimuli) using R’s *anova* function.

To assess the title-effect on different ratings the following preregistered LMM formulas were applied:

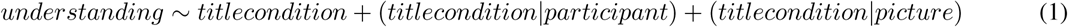

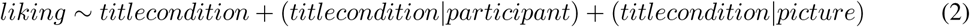

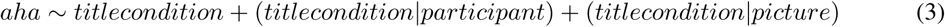

Here only data from artwork trials were used (omitting Mooney trials). We applied *dummy coding* to compare the average rating in the “titled” to the “untitled” category (baseline).

The random slope effect for participant x title condition was preregistered as we expected participants to differ in their sensitivity to a title effect. We had also preregistered a random-slope-effect for image x title-condition as we expected the title-effect to differ by image. However, the fact that each participant saw each image only once in one title-condition might weaken this comparison – in Supp. A.2.2 we thus report alternative models with random intercepts for each image; this did not drastically alter the results.

To test our hypothesis that visual art would receive higher liking ratings than Mooney images we used the following formula:

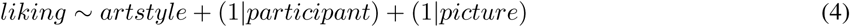

Here all trials were used. We applied *dummy coding* to compare the average liking rating in all cells to the Mooney images (baseline).

To benchmark the strength of *aha* elicited by visual art and their titles to the feeling of *aha* caused by the Mooney images we fit the following formula:

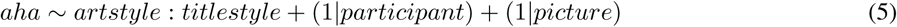

Here only data from Mooney images, and artwork shown with their real titles were used (omitting trials labelled as “untitled”). We applied *dummy coding* to compare the average *aha* rating in all interaction cells to the Mooney images (baseline).

To test the linear relation between curiosity and *aha* within the different title conditions we applied the following LMM formula:

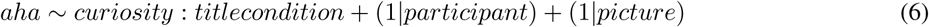

Here data from all trials were used (title condition coded as either “untitled art”, “titled art”, or “mooney”).

Likelihood ratio tests for all the above models were performed against matching intercept-only models:

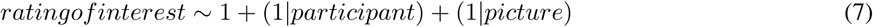

##### Linear mixed models with quadratic terms

Data visualization using local regression (see below) prompted us to further model the data using nonlinear quadratic effects, as an exploratory follow-up. Likelihood ratio tests for the model with quadratic polynomial terms against a matching model with linear terms were used to assess if this procedure significantly improved model fit. We tested whether the effect of curiosity on aha was quadratically modulated by title style (eqs. 8,9), whether the effect of aha on liking was quadratically modulated by art style (eqs. 10,11) and whether the effect of aha on understanding was quadratically modulated by art style (eqs. 12,13).

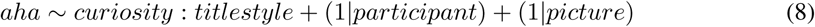

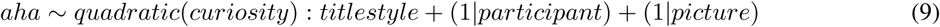

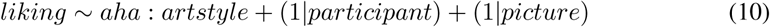

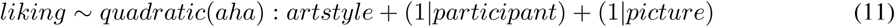

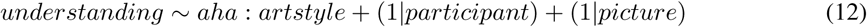

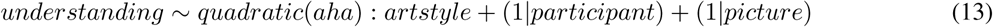

##### Local regression

Local regression plots were generated using locally weighted scatterplot smoothing (LOWESS) as implemented in the *seaborn* module for Python (Waskom, 2021, version 0.11.2).

#### Data visualization

Data visualizations were generated with custom code using the *seaborn* Python module (Waskom, 2021, version 0.11.2).

### 2.2 Experiment 2 – EEG study

#### Participants

##### Sample size estimation

Given the lack of previous EEG research on the title effect or insight with art, we decided to power the study based on the behavioral effect we aimed to observe: a main effect of the stimulus category on the *aha* ratings.

We estimated the target sample size *a priori* using G*Power (Faul et al., 2007, version 3.1). Again, we powered our experiment for a repeated measures ANOVA design similar to the LMMs used in Exp 1. We used the parameters “ANOVA: Repeated measures within factors” with *alpha* = .05, *power* = .8, Effect size *f* = 0, 1758631 (equivalent to *η*^2^ = 0.03, a small effect size), 1 group, 7 measurements (3×2 + 1 design, see below), and correlation among repeated measures *r* = .42. The correlation between repeated measures was estimated based on the online study data, as the average correlation coefficient of the participant-average insight-ratings across the 7 stimulus factor combinations used in the present study.

This power analysis resulted in a target sample size of N = 38. To account for dropout, we decided to collect data from at least N = 45 participants.

##### Recruitment

Participants were convenience-sampled from the general public in the Rhine-Main metropolitan region in Germany. Recruitment procedure was the same as for Exp. 1. Inclusion criteria for recruitment were age between 18–55 years, good command of the German language, good eyesight and no need to wear glasses during the study (as this decreases the quality of the eye tracking), no known neurological disorders, and no participation in the previous online study.

##### Final sample

50 participants enrolled and finished the data collection. 1 participant with ongoing neurological problems (essential tremor) enrolled despite the inclusion criteria, and was thus excluded from analysis post-hoc, leaving 49 datasets for analysis. Some participants had periods of missing data (EEG, ECG, eye tracking, and/or ratings) due to minor bugs or poor quality recordings, but were not excluded from analysis.

The participants were between 18–52 years old (mean = 28.0 years, std = 7.2 years). 28 were female, 19 male, and 2 indicated ‘other’. 8 were left handed, 41 right handed, and 0 ambidextrous (Flinders Handedness survey; Nicholls et al., 2013). 17 participants showed left eye, 28 right eye, and 4 no eye dominance (assessed by the experimenter; see below). Participants indicated having received between 8–27 years of education (mean = 17.3 years, std = 3.9 years; with 1 participant not providing these data). The sample showed an extreme bias towards highly educated people: all 49 participants indicated the German Abitur or a university degree as their highest qualification. 11 in 49 participants reported current or past episodes of mental or psychological disorders with 7 of these 11 indicating current medication. Although it was not a criterion for exclusion, we also assessed the participants’ caffeine intake on the day of the experiment: it ranged between 0–3.61 mg/kg body weight (mean = 1.06 mg/kg, std = 0.923 mg/kg). 8 out of 49 participants indicated zero caffeine intake before participation. See Supp. Tabs. A.1 and A.2 for full details on participant demographics.

All participants received monetary compensation of 7 per 30 minutes and gave their informed written consent prior to participation. The study adhered to the ethical standards of the Declaration of Helsinki and was approved by the local ethics committee (Ethics Council of the Max Planck Society) under the Research Ethics Framework Application for Standard Laboratory Studies. Data were collected from March 2022 through July 2022.

#### Stimuli

Each participant saw 48 art stimuli and 16 Mooney images on a calibrated LCD monitor. These stimuli were preselected from the larger stimulus set of 120 art stimuli and 20 Mooney images from Exp. 1 using the rating data from Exp. 1.

##### Image selection

From the 140 stimuli described above for Exp. 1 we selected 64 based on a linear mixed model with the formula:

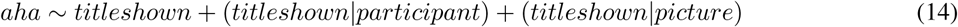

i.e. a model that predicts the insight rating from the title condition (real title or untitled), with random slopes and intercepts for each participant and each stimulus (meaning that the strength of the title effect can vary for each participant and each stimulus). Random slope coefficients for all stimuli were ranked and we chose the 16 stimuli with the highest slope within each picture category (abstract art, semi-abstract art, figurative art, Mooney images) – these were the images with the strongest title-effect on *aha* ratings in Exp. 1. The third fully crossed condition from Exp. 1 (type of title: descriptive, explanatory, evocative) was omitted here, as this would have drastically decreased freedom in the selection process. The ranked slope coefficients and the selection are depicted in Supp. Fig. A.1. All stimuli (images and titles) are depicted in Supp. A.1.4.

#### Experimental design

We used a within participant factorial design with repeated measures. It can be formalized as a 3×2 + 1 design: participants saw visual art stimuli in 3 degrees of abstraction (figurative, semi-abstract, abstract), in either of 2 title reveal conditions (real title, or “untitled” dummy title). These factors were fully crossed, with 8 trials for each of the 6 factor combinations (48 art stimuli). Additionally, 16 trials with Mooney images were presented per participant. This led to a total 64 trials per participant. As opposed to Exp. 1, all participants saw the same stimuli though with randomly drawn title assignments.

##### Task procedure

First the participant answered a set of ancillary questionnaires (see below) implemented in Labvanced (Scicovery GmbH, Germany). The participant entered the cabin, and the EEG cap and peripheral electrodes (ECG) were attached. EEG electrodes were treated until impedance were below or close to 10 kOhm using electrolyte gel. After finishing preparations, the effects of chewing, blinking, and contracting neck muscles on the EEG signal were demonstrated to the participant to raise awareness for these sources of artifacts, along with a demonstration of individual alpha oscillations during a short eyes closed period. Instructions for the main experiment were displayed, followed by 3 practice trials (1 Mooney image and 2 art stimuli together with their real titles; these stimuli were not presented again in the main experiment). After the training block, the experimenter adjusted the participants’ sitting position, head rest, and set up the eye tracker. From then on, the participant was asked to keep her position in the head rest and restrain from chewing, swallowing or moving the head during trial periods. Trials were presented in blocks of 10 stimuli, and after each block participants could choose to take a short self paced break to relax. The paradigm was largely identical to Exp. 1 (see also Fig. 1a). Yet unlike the online study, all pictures and titles were preceded by a fixation cross, presented for 2 *−* 2.5*s* (jittered), the inter stimulus interval was 1 *−* 1.5*s* (jittered), and all rating sliders (including aesthetic preference and felt understanding) were presented on separate pages. The routine was implemented in custom code using PsychoPy (Peirce, 2007). After finishing the experiment, participants filled a short debriefing questionnaire (implemented in Labvanced, Scicovery GmbH, Germany).

##### Randomization

The assignment of art stimulus to the titled or untitled condition was fully randomized across participants. However, the randomization was controlled such that within each art style (abstract, semi-abstract, figurative) the final number of title and no-title trials was balanced. To keep the assignment distribution balanced for the whole data collection, pseudorandom assignment was done only for every other participant (1,3,5, and so on), while the subsequent participants (2,4,6, and so on) saw reversed assignments of the preceeding participant. The stimulus order for each participant was randomized, with fully intermixed trials (not more than 2 stimuli of the same type in a row).

#### Behavioural measures and questionnaires

In addition to the screening tools used in Exp. 1, caffeine intake before the visit (Bühler et al., 2014), and handedness (FLANDERS; Nicholls et al., 2013) were assessed via systematic questionnaires. Ocular dominance was determined by the experimenter using a variation of the Porta test (as described in Roth, 2002).

All trial-wise ratings (curiosity, *aha*, aesthetic liking, understanding) were collected using a continuous scale with a slider controlled by moving the mouse. Both ratings and response times were logged. In order to discourage lazy non-responding, participants could not log a response without having first moved the mouse. The ends of the scales were marked with “+” and “–” and each scale was accompanied by the corresponding questions: “Wie neugierig sind Sie auf den Titel / die Auflösung?” (“How curious are you for the title / the solution?”) for the curiosity rating, “Hatten Sie ein Aha! Erlebnis?” (“Did you have an Aha! experience?”) for the *aha* rating, “Wie ästhetisch ansprechend fanden Sie das Bild?” (“How aesthetically appealing did you find the picture?”) for the aesthetic rating, or “Denken Sie, dass Sie das Bild verstanden haben?” (“Do you think you understood the picture?”) for the understanding rating respectively. The order of these ratings was fixed. In the detailed task description preceding the study, all ratings were explained – the full descriptions can be found in the Supp. A.1.2.

#### Data acquisition and devices

##### Study environment

EEG recording took place in an acoustically shielded cabin (model: IAC 120a, IAC GmbH, Germany; internal dimension: 2.74 *∗* 2.54 *∗* 2.3*m*). Participants placed their chin on a chin rest with forehead support (SR Research Head Support, SR Research Ltd., Canada); the distance between the chin rest and the screen was 72 cm. The height of the desk was adjusted such that participants sat in a comfortable upright position that they were able to sustain for the time of the study. All data were collected by the same experimenter, with assistance during EEG gel preparation. The participant could contact the experimenter any time via a room microphone installed in the cabin. The experiment was run on a PC running 64-bit Microsoft Windows 7.1.7601 service pack 1 (Microsoft Corporation, USA), using PsychoPy3 standalone software (Peirce, 2007, version 3.0.7). Visual stimuli were presented on a 24 inch BenQ XL2420Z screen (BenQ Corporation, Taiwan) with nominal frame rate of 144 Hz, resolution 1920 × 1080 px (mirrored for the experimenter). Luminance and color calibration was performed using the screen’s settings and validated using a chromameter (Konica Minolta CS-150) with a staircase routine from the PsychoPy monitor center; the measured gamma curve can be found online in the OSF repository.

##### EEG and peripheral physiology data acquisition

Bio-signals were amplified using a BrainVision actiCHamp 128 Amplifier with BIP2Aux Adapters (Brain Products GmbH, Germany) with 24-bit digitization and an analogue band-pass filter from DC - 280 Hz. The amplifier was DC battery powered by 2 ActiPower PowerPacks (Brain Products GmbH, Germany). Data was recorded with 1 kHz sampling frequency using BrainVision Recorder (version 1.21.0303; Brain Products GmbH, Germany) from an independent recording PC running Microsoft Windows 7. TTL triggers from the experimenter PC were sent to the amplifier via parallel port. EEG data were collected using a 64 channel actiCAP system with active Ag/AgCl electrodes without active shielding (Brain Products GmbH, Germany), placed according to extended international 10-20 localization system (Commitee, 1958; Oostenveld & Praamstra, 2001) with FCz online recording reference and AFz ground. Forehead and skin behind the ears were wiped with alcohol to enhance the impedance. The cap was positioned by centering the Cz electrode on the axes nasion to inion and left ear to right ear. After the end of the data collection, electrode positions were digitized using a Polhemus FastTrak system (Polhemus, USA) and Curry software (version 8; Compumedics Neuroscan). ECG data was recorded by the EEG amplifier as a bipolar auxiliary channel: electrodes were placed on the right mid clavicle and lower left rib cage, in correspondence with the II Einthoven’s derivation, using disposable electrodes (Covidien, USA). To enhance EEG data quality the participants were asked in the invitation mail to not apply hair styling products.

##### Eye tracking data acquisition

Binocular gaze position (coordinates on the screen) and pupil size were recorded using a desktop mount EyeLink 1000 Plus eye tracking system (SR Research Ltd.) and EyeLink 1000 Plus Host software running on an independent recording PC. The experimenter PC and the EyeLink recording system were linked via ethernet, and the tracker system was controlled using the PyLink Python module (SR Research Ltd., version 1.11.0.0). Data was recorded with a sampling frequency of 500 Hz, with recording breaks between each block. The eye tracker was (re-)calibrated before every trial block using a 13 point horizontal and vertical calibration routine. To enhance data quality the participants were asked in the invitation mail to not apply make-up, especially eyeliner and mascara.

#### Data analysis

Data preprocessing, processing, and statistical analysis was done on the institute’s high performance computer (HPC) using Python (version 3.10.4), R (version 3.6.1) in conda virtual environments, and Matlab (The MathWorks Inc., version R2019b). Jupyter notebooks reproducing figures and results are publicly available online at OSF.

##### EEG data early preprocessing

EEG Data were transferred into BIDS format (Pernet et al., 2019, version 1.6.0) using the MNE-BIDS Python module (Appelhoff et al., 2019, version 0.10). Diverging from the preregistered analysis plan, we did not apply a preprocessing and cleaning pipeline including ICA. Instead we used the PREP pipeline (version 0.56.0, EEGLAB plugin in MATLAB, using the default parameter set; Bigdely-Shamlo et al., 2015) a less aggressive fully automatic early-stage preprocessing pipeline consisting of only line noise removal, rereference to (robust) average, and detection and interpolation of noisy channels. This decision was based on 1) recent work suggesting that heavy preprocessing and/or data rejection can sometimes be less helpful than intended (Delorme, 2023) as any gain in SNR has to outweigh the decrease in statistical power due to data rejection, and 2) work that showed that ICA based data cleaning can leave residuals in the data (Dimigen, 2020; Robbins et al., 2020) and that those residuals can remain problematic with sensitive machine learning algorithms (Quax et al., 2019; Thielen et al., 2019), as planned in this study.

##### Time resolved decoding from EEG, pupil size, and instantaneous heart rate

We sought to decode different targets (stimulus categories and ratings) from the multivariate physiological data (EEG, pupil size, heart rate). Time-resolved decoding was implemented in a sliding window approach: independent decoding models were fit at each time sample across the 64 EEG sensors (or 2 pupil traces, or single heart rate channel). This analysis yielded a decoding time course indicating if and when a rating or condition could be linearly decoded from the physiological signals. The decoding time courses were then subjected to second-level cluster-correction across participants (see below).

##### Decoding pipelines

We built 3 different categorical classification pipelines for repeated presentation (1st/2nd presentation of the image), the stimulus condition (Mooney/art) and the title condition (real or dummy title) and 4 different float-value prediction pipelines for the different ratings (curiosity, *aha*, liking, understanding; all transformed to participant wise z-scores). The decoder design closely follows Gwilliams and King (2020), implemented in MNE Python: We applied logistic regressions for categorical classification (scikit-learn; Pedregosa et al., 2011, default parameters: C = 1) and ridge regression for continuous prediction (default parameters: alpha = 1). Decoding performance was assessed in a stratified 10-fold cross-validation. Decoding accuracy was scored using an AUC metric for logistic regression, and spearman R for ridge regression. All decoders used data normalized by the mean and the standard deviation of the training set.

##### Data features

These decoding pipelines were applied to different data: first, to time domain EEG (single-trial ERPs) on all sixty-four channels – continuous PREP preprocessed EEG data were band-pass filtered (0.1-40 Hz, FIR, MNE default parameter) and cut into epochs (−0.5-5 s around stim onset); baseline correction was applied by subtracting the average of the baseline period (−0.5-0 s) from the entire epoch (for each trial and channel individually). Data were re-referenced to infinite reference (REST; Yao, 2001, implemented in MNE). Lastly, a fully automatic trial-rejection procedure was applied (based on the min-max amplitude of the trial data within each participant): a gamma distribution was fit to the empirical distribution of min-max amplitudes of all trials used in the respective decoding subset, and epochs with a value outside the 90th percentile of the fitted distribution were rejected as outliers. While this step reduced our trial number (numbers below), we deemed it necessary, as the scaling step makes the decoding pipeline vulnerable to outliers.

The second dataset for decoding was frequency band power in the canonical EEG frequency bands on all sixty-four channels – PREP preprocessed continuous EEG data were band-pass filtered to the respective frequency band (delta: .1-4 Hz; theta: 4-8 Hz; alpha: 8-23 Hz; beta: 13-30 Hz; gamma: 30-80 Hz – FIR, MNE default parameter) and cut into initially longer than needed epochs (−2–6.5 s for delta and −1–5.5 s for the other bands, to avoid edge artifacts in the analyzed data); no baseline correction was applied at this stage. The data were re-referenced to infinite reference. To reveal oscillatory activity the evoked response was subtracted from every single trial and the envelope of the Hilbert transformed EEG was calculated as the time-frequency response (TFR). Lastly, epochs were cropped to the time window intended for decoding (−0.5–5 s) and baseline correction was applied by subtracting the average of the baseline period (−0.5–0 s) from the entire epoch (for each trial and channel individually). The same trial-rejection as for ERP data was performed.

The third dataset was the binocular pupil trace. Raw continuous data were low-pass filtered (cutoff 10 Hz, FIR, MNE default parameters). Missing data due to eye blinks were linearly interpolated (using MNE’s *interpolate_blinks* function), and trials with a min-max amplitude two standard deviations larger than the average were rejected as outliers.

The fourth dataset was instantaneous heart rate. ECG recordings were preprocessed using the neurokit2 python module (version 0.2.7; Makowski et al., 2021): recordings were checked for inverted signal using neurokit2’s *ecg_invert* function and inverted if necessary, cut into initially longer than needed epochs (−3.5–8 s; no baseline correction) followed by data cleaning (neurokit’s *ecg_clean* function), R-peak detection (neurokit’s *ecg_peak* function), and conversion to instantaneous heart rate (neurokit’s *ecg_rate* function, all with default parameters). Lastly, epochs were cropped to the time window intended for decoding (−0.5–5 s). Recording quality in each trial was assessed using neurokit’s *ecg_quality* function (using the scoring method following Zhao & Zhang, 2018) and all trials not scored as “excellent” were excluded from analysis.

ERP, TFR, pupil, and heart rate data were then down-sampled to 50 Hz prior to decoding (using MNE’s *resample* function with default parameters).

##### Trial rejection

Below we report trial rejection across all the different decoding pipelines within a given data modality. For decoding of repeated exposure (1st or 2nd presentation) and stimulus category (Mooney or visual art) all 64 trials were used (art and Mooney images), all other analyses used only art trials (48 trials per participant). The criteria for rejecting outlier trials is described above per data modality. In addition, within a given decoding pipeline a participant was skipped entirely if 1) there were fewer trials left than required for cross validation, or 2) if the variance in a participant’s ratings was too low for the decoding model to converge.

The cleaning and trial rejection procedures described above led to a moderate trial rejection in the low percentage range. Note that the maximum trial number varied by pipeline. ERP – data from 46–49 participants (mean=48.2 out of 49, or 98.45%) and within those 91.3–95.6% (mean=93.51%) of the maximum possible trials were used. EEG delta – data from 47–49 participants (mean=48.3 out of 49, or 98.64%) and within those 91.0–95.3% (mean=92.85%) of the maximum possible trials were used. EEG theta – data from 41–49 participants (mean=47.9 out of 49, or 97.71%) and within those 87.8–93.3% (mean=91.06%) of the maximum possible trials were used. EEG alpha – data from 45–49 participants (mean=48.3 out of 49, or 98.64%) and within those 88.7–94.5% (mean=91.46%) of the maximum possible trials were used. EEG beta – data from 47–49 participants (mean=48.5 out of 49, or 98.89%) and within those 93.3–96.6% (mean=95.01%) of the maximum possible trials were used. EEG gamma – data from 47–49 participants (mean=48.4 out of 49, or 98.70%) and within those 91.8–96.6% (mean=94.29%) of the maximum possible trials were used. Pupil dilation – data from 47–49 participants (mean=48.3 out of 49, or 98.64%) and within those 98.9–99.4% (mean=99.07%) of the maximum possible trials were used. Heart rate – data from 45–49 participants (mean=47.0 out of 49, or 95.98%) and within those 95.8–98.3% (mean=97.12%) of the maximum possible trials were used.

##### Second-level statistical analysis of decoding accuracies

Significance of the univariate and multivariate decoding results was tested using a one-sample permutation cluster test (Maris & Oostenveld, 2007) implemented in MNE’s *permutation_cluster_1samp_test* function.

To account for noise in the individual decoding traces, decoding timecourses for each participant were mildly smoothed prior to the permutation test (using scipy’s *gaussian_filter* function, with sigma=1.0).

The data were centered around the theoretical chance level for the respective metric (0.5 for AUC, 0 for Spearman R). Then the permutation cluster test was performed: first, one-sided one-sample t-tests were done at each point in time of the original data. Spatio-temporally adjacent data-points were clustered based on a cluster-forming threshold equivalent to *p <* 0.05 and the test statistic for each cluster was found as the sum of *t*-values across time. Then a permutation null distribution was generated (10000 permutation steps): randomized data were created from the original data with random sign flips, and a new set of clusters and test statistics were formed. The original test statistics were then compared to the permutation null distribution to assess statistical significance.

#### Data visualization

Data visualizations for the time-resolved decoding analysis were generated with custom code using matplotlib Python module (Hunter, 2007, version 3.8.0) and exported in .svg format.

## 3 Results

### 3.1 Experiment 1 – Behavioral ratings of curiosity, insight, aesthetic preference, and understanding

The stimulus set was designed to contain equal numbers of trials from different art styles (abstract, semi-abstract or figurative art) and title styles (descriptive titles, explanatory titles, evocative titles). Together with varying the title condition (real or uninformative dummy title) and the contrasting category of Mooney images, this allows for a large number of different analyses comparing different ratings across varying type and amount of information provided to the participant. The majority of hypothesis tested below were preregistered (https://doi.org/10.17605/OSF.IO/PAGVH), and additional exploratory follow-up analyses are clearly marked as such. Full details of all reported linear mixed models (LMMs) can be found in Supp. A.2.2.

#### Informative titles increase aha and felt understanding, but not aesthetic appeal

We first tested for the presence of a “title effect” on the various behavioral ratings (i.e. differences in ratings between trials with meaningful titles compared to “untitled” trials). To do so we applied three preregistered LMMs (eqs. 1–3). In this analysis only trials with art stimuli were used, ignoring the Mooney images. However, for comparison figures include data for the Mooney images as well.

We were able to confirm our hypothesis and replicate previous findings of a positive title effect on perceived understanding, but not on aesthetic ratings (Fig. 1b; splitting the stimuli by art style, i.e. grade of abstraction). The title effect on felt understanding was substantial (*slope* = 11.06, *CI* = [8.11; 14.02], *p <* 0.001; this slope indicates that the average understanding rating with titles is 11.06 points higher than without titles on the used 0-100 points rating scale). The likelihood ratio test was significant (*χ*^2^[5] = 173.88, *p <* 0.001). The title effect on aesthetic rating, however, was only marginal (*slope* = 1.69, *CI* = [*−*0.16; 3.53], *p* = .073) and the likelihood ratio test was not significant (*χ*^2^[5] = 6.53, *p* = .258), thus rejecting our preregistered hypothesis. We did not see hints that this was different in any of the art or title styles (see e.g. Fig. 1b).

Besides revisiting these previously investigated title effects, we confirmed our preregistered novel hypothesis that meaningful titles would also increase *aha* ratings (see Fig. 2a). The effect of titles on *aha* was even stronger than on understanding ratings (*slope* = 17.89, *CI* = [13.80; 21.97], *p <* 0.001) and the likelihood ratio test was significant (*χ*^2^[5] = 564.95, *p <* 0.001). Importantly, the effect of the title manipulation on *aha* ratings varied drastically across stimuli, with a few of them even receiving lower *aha* ratings with the actual title than with the dummy titles (see Supp. A.1.3 and Fig. Supp. A.1).

**Figure 2:**
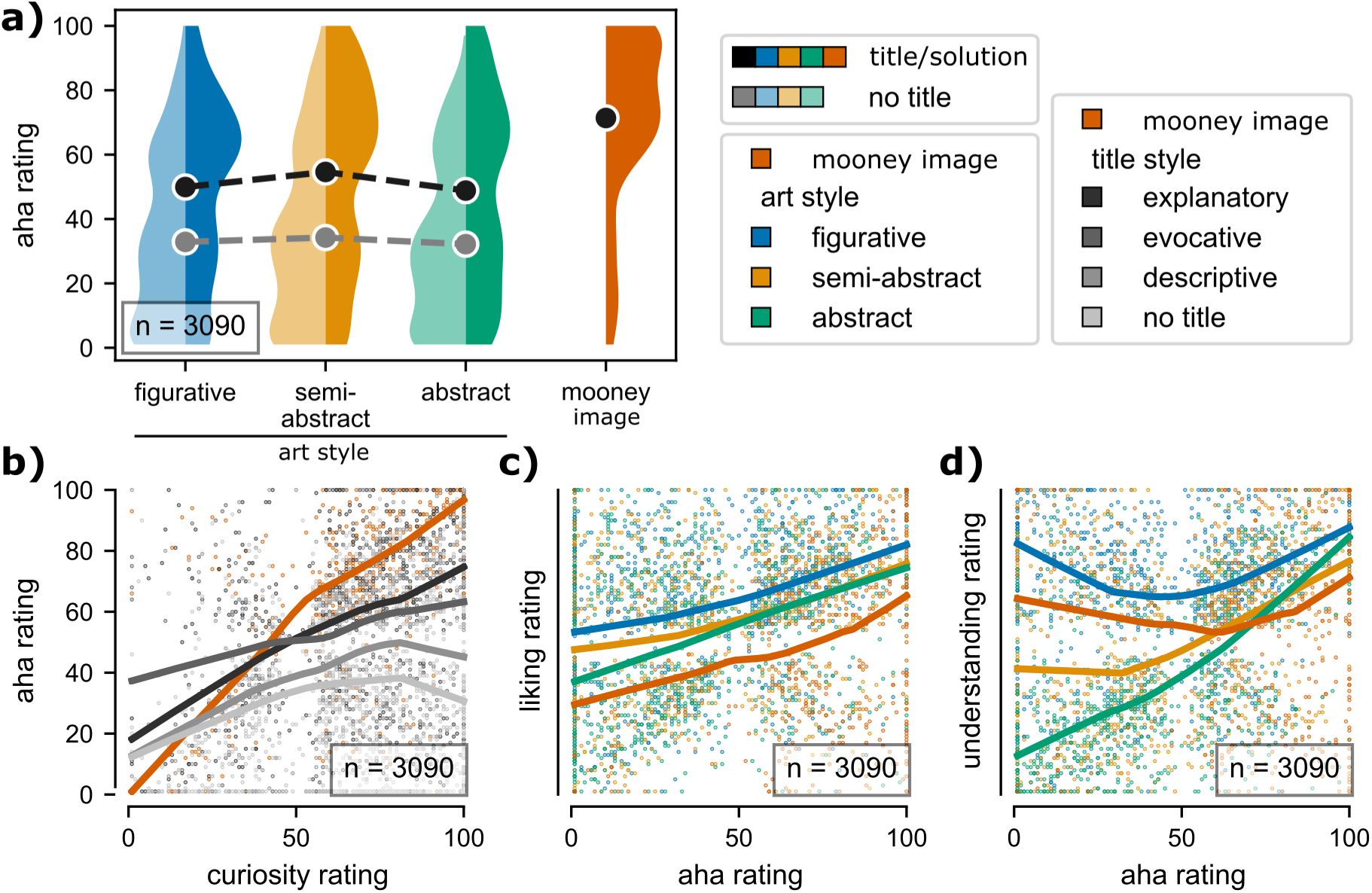
Using the title effect to evoke insight with visual art. **a) Establishing a title effect on felt *aha*:** the title manipulation strongly affected the participants’ feeling of *aha* (present in all 3 categories of artwork); the effect was even larger than on felt understanding. **b) Relation between curiosity and insight interacts with information content of solution cues:** we replicated a strong linear relation between curiosity and *aha* for Mooney images (orange LOWESS curve; previously reported by Van de Cruys et al., 2021) and saw a similar, slightly weaker relation for artwork with explanatory titles (i.e., titles leaving no uncertainty about what is depicted in the artwork; darkest LOWESS curve). However, for the titles that carry less or no clear additional information the association became weaker (e.g., descriptive titles that solely describe in words what is depicted on the image and “untitled” dummy titles that add no semantic information at all; light gray LOWESS curve). Evocative titles, which linked to a newly introduced concept not readily depicted in the visual image (dark-gray LOWESS curve), exhibited a weaker linear relationship, with higher *aha* ratings in low curiosity trials, compared to the other title conditions. While the LOWESS curves suggest partly nonlinear interactions with the stimulus category, a likelihood ratio test of LMM with linear terms against LMM with added quadratic terms did not statistically support this observation. **c) Positive relation between *aha* and aesthetic liking:** a positive linear association was present in all stimulus categories. As reported before, average aesthetic liking was highest for figurative art (blue LOWESS curve). Mooney images (orange LOWESS curve) had the lowest average aesthetic ratings. **d) Relation between *aha* and understanding nonlinearly interacts with stimulus category:** For abstract art (green LOWESS curve) there was a linear relation between ratings of *aha* and understanding. Figurative art (blue LOWESS curve), on the other hand, exhibited a markedly non-linear U-shaped relation with high understanding both for very low and very high *aha* trials. Semi-abstract art (yellow LOWESS curve) exhibited an intermediate pattern. The Mooney images (orange LOWESS curve) also showed a U-shaped relation, but on average lower understanding ratings than figurative art. A likelihood ratio test of LMM with linear terms against LMM with added quadratic terms confirms that the nonlinear interaction effects suggested by these LOWESS curves.

#### Higher liking, but lower aha for most art vs. Mooney images

We only partially confirmed our preregistered hypothesis that visual art would receive higher liking ratings than Mooney images (see also Fig. 1b). Our LMM tested for the overall effect of image category (abstract, semi-abstract or figurative art both titled and untitled, or Mooney images; eq. 4) with Mooney images as the baseline (*intercept* = 50.87, *CI* = [46.19; 55.56], *p <* 0.001). Figurative art (*slope* = 10.62, *CI* = [5.65; 15.59], *p <* 0.001) and semi-abstract art (*slope* = 5.58, *CI* = [0.61; 10.55], *p* = 0.028) received significantly higher liking ratings than Mooney images, but abstract art did not (*slope* = 1.68, *CI* = [*−*3.29; 6.65], *p* = 0.507). A likelihood ratio test for the model was significant (*χ*^2^[3] = 24.26, *p <* 0.001).

We had further preregistered the hypothesis that Mooney images would cause, on average, higher *aha* than visual art. For this benchmark we fit an LMM (eq. 5) on *aha* ratings from Mooney image trials and all titled art trials (omitting trials with art stimuli in the untitled condition). Mooney images were again set as the baseline (*intercept* = 71.27, *CI* = [66.20; 76.33], *p < .*001). As expected, we observed lower *aha* ratings for all other categories, and the difference was significant for all art style x title style combinations. Semi-abstract art with explanatory titles came the closest^1^ (*slope* = *−*11.50, *CI* = [*−*17.71; *−*5.29], *p < .*001, then followed (ordered by slope): figurative art with evocative titles (*slope* = *−*14.69, *CI* = [*−*22.45; *−*6.92], *p <* 0.001), semi-abstract art with evocative titles (*slope* = *−*15.68, *CI* = [*−*23.47; *−*7.90], *p <* 0.001), abstract art with explanatory titles (*slope* = *−*18.26, *CI* = [*−*24.46; *−*12.06], *p <* 0.001), figurative art with explanatory titles (*slope* = *−*19.24, *CI* = [*−*25.44; *−*13.03], *p <* 0.001), abstract art with evocative titles (*slope* = *−*24.39, *CI* = [*−*32.17; *−*16.61], *p <* 0.001). The overall lowest aha ratings were observed for stimulus groups with descriptive titles, regardless of the degree of visual abstraction (semi-abstract art with descriptive titles: *slope* = *−*28.95, *CI* = [*−*36.73; *−*21.18], *p <* 0.001, abstract art with descriptive titles: *slope* = *−*29.92, *CI* = [*−*37.68; *−*22.16], *p <* 0.001, and figurative art with descriptive titles: *slope* = *−*32.05, *CI* = [*−*39.82; *−*24.29], *p <* 0.001). A likelihood ratio test for the overall model was significant (*χ*^2^[9] = 89.99, *p <* 0.001). Note that Mooney images received higher *aha* ratings than art stimuli specifically preselected for receiving the highest “surprise” ratings among a very large corpus (see Methods).

We confirmed another preregistered hypothesis and replicated a robust positive linear relation between baseline curiosity (rated after seeing an image for the first time, but before seeing the title) and subsequent *aha* ratings in the Mooney image trials (*slope* = 0.63, *CI* = [0.58; 0.69], *p <* 0.001). This matches previously reported values for this relation of *r* = 0.68 (on the image level, see Van de Cruys et al., 2021). The model (eq. 6) also indicated a significant effect with smaller slope for titled art (*slope* = 0.37, *CI* = [0.32; 0.41], *p <* 0.001). For untitled art the slope was again much lower, though still significant (*slope* = 0.11, *CI* = [0.07; 0.15], *p < .*001). A likelihood ratio test was significant (*χ*^2^[5] = 773.51, *p <* 0.001).

#### Title type shaped ratings in non-trivial ways

We further explored the relationship between curiosity and *aha* using local regression (LOWESS, see Fig. 2b). Qualitatively, this suggested there might be a non-linear interaction with the type of title cue: while Mooney images and art with explanatory titles (i.e. titles leaving little uncertainty about what was depicted) exhibited a relatively linear relationship between curiosity and *aha*, this association was weaker and markedly nonlinear for the titles that should reduce uncertainty to a lower degree or not at all (e.g., descriptive titles that solely described in words what was depicted on the image and “untitled” dummy titles that added no new information at all). Here, for low to medium-high curiosity trials the LOWESS curve of *aha* ratings monotonically increased, but then flattened and even fell off again for high curiosity trials. This would be consistent with a disappointment effect for trials with high curiosity but lacking subsequent relief or closure (i.e. without actual information gain from the title). Evocative titles, which introduced and linked to a new concept not readily depicted in the visual image, again exhibited a linear relationship, with an overall lower slope than explanatory titles or Mooney images, but higher *aha* ratings in the low curiosity trials than all other title conditions. While these trends seemed very intriguing, a likelihood ratio test showed that an LMM with added quadratic terms (eq. 9) did not fit the data significantly better than a model with only linear terms (eq. 8; *χ*^2^[6] = 0, *p* = 1), thus suggesting that the observed pattern might be spurious. For full model details see Tabs. Supp. A.13 and A.14.

#### Aha is linearly related to liking and non-linearly to understanding

We saw a significant linear relation between *aha* and liking ratings, thereby confirming another preregistered hypothesis. The model (eq. 10) suggests that this effect was significant for all image categories (Mooney images: *slope* = 0.15, *CI* = [0.10; 0.20], *p <* 0.001, semi-abstract art: *slope* = 0.24, *CI* = [0.19; 0.29], *p <* 0.001, abstract art: *slope* = 0.26, *CI* = [0.21; 0.31], *p <* 0.001, figurative art: *slope* = 0.32, *CI* = [0.27; 0.37], *p <* 0.001 – combining titled and untitled trials). A follow-up local regression suggested that this relation was indeed linear and did not depend much on the picture type (see LOWESS curve in Fig. 2c). A likelihood ratio test confirmed that a model with added quadratic terms (eq. 11) did not fit the data significantly better than the model with linear terms (eq. 10; *χ*^2^[4] = 0, *p* = 1). Our data thereby shows that previous observations with Mooney images (Muth & Carbon, 2013) can generalize to other stimulus domains, here visual art.

For the relation between *aha* and understanding, local regression suggested a nonlinear interaction with the stimulus category. Indeed we had hypothesized to find understanding ratings interact with the level of stimulus abstraction (preregistered, though no exact hypothesis how the interaction would look). While the association was linear for abstract art, it appeared more quadratic (U-shaped) for semi-abstract and figurative art. Mooney images also exhibited a more U-shaped association (see LOWESS curves in Fig. 2d). Here a likelihood ratio test confirmed that the model with quadratic terms (eq. 13) did indeed fit the data significantly better than a model with linear terms (eq. 12; *χ*^2^[4] = 115.03, *p <* 0.001). Quadratic terms were significant for figurative art (*estimate* = 216.92, *CI* = [118.04; 315.44],*p <* 0.001), Mooney images (*estimate* = 141.51, *CI* = [12.73; 270.30],*p* = 0.031), and semi-abstract art (*estimate* = 96.42, *CI* = [3.87; 188.98],*p* = 0.041) but not for abstract art (*estimate* = *−*19.48, *CI* = [*−*116.45; 77.48],*p* = 0.694; for the full model details see Tabs. Supp. A.17 and A.18). The observed pattern would be plausible if low aha–high understanding trials were due to a high initial understanding of the images (already without titles or Mooney solution) which would leave little uncertainty to be reduced by the additional information (see also Fig. Supp. A.3).

### 3.2 Experiment 2 – Neuronal correlates and the temporal sequence of information processing

To study the temporal profile of the epistemic arc we repeated the experiment recording observers’ EEG, heart rate, and pupil size. The experimental design was preregistered (https://doi.org/10.17605/OSF.IO/6KW9N) but as this part of the study was largely exploratory we had not committed to any specific analysis. The physiological signals were analyzed using a time-resolved decoding approach to show whether and when relevant information was represented in the physiological data. We aimed to decode both the experimental conditions and the various behavioral ratings collected during the trials. Importantly, we ran these decoding pipelines on data from both the pretitle trials and the posttitle trials separately, to reveal potential effects of the interim presented title or Mooney solution (except in one pipeline specifically decoding whether trials belonged to the pretitle or posttitle condition). We further ran the pipelines on a difference signal (posttitle – pretitle trials); these results are reported in Supp. A.2.4. Findings are described below, and significant decoding results are shown in Figs. 3 and 4. Decoding plots for all data modalities and all conditions are compiled in Supp. A.2.5.

**Figure 3:**
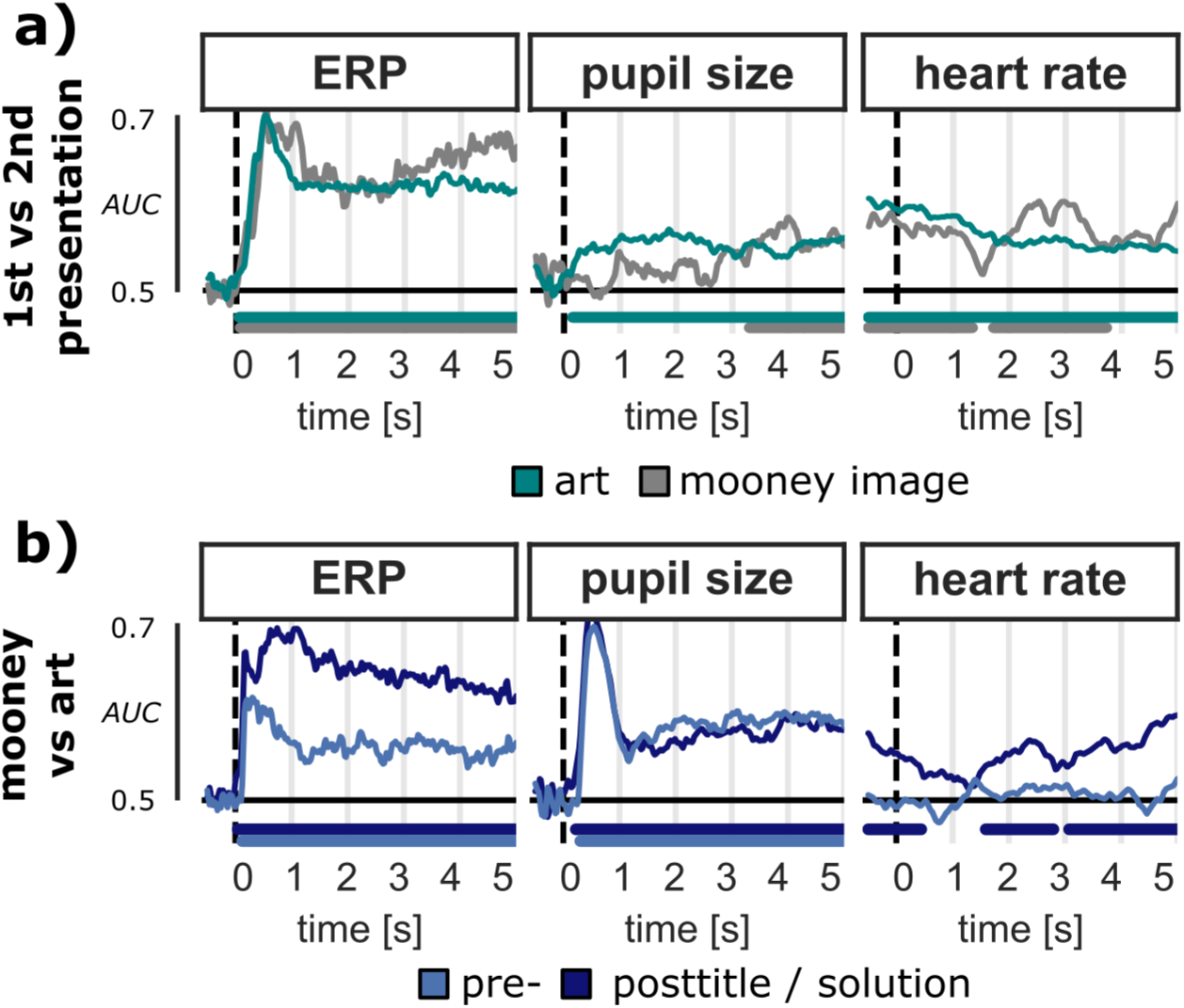
Decoding repeated presentation and image category (artwork and Mooney trials) **a) 1st or 2nd presentation of the image:** decoding was reliably possible from almost all data modalities but best from the time-domain EEG (significantly above chance throughout the full trial, with a marked peak around 0.5–1 s after image onset). In pupil trace decoding worked better with art than with Mooney images, where decoding became significant only at the end of the trial. This might be partly due to smaller trial counts in this category (48 art and 16 Mooney trials per participant). Instantaneous heart rate appeared to predict the category slightly better early in the trials. Taken together this strongly suggests that the stimuli were not processed identically during repeated presentation. **b) stimulus domain (Mooney image or artwork):** the image category could be reliably decoded from the beginning of image presentation for EEG, pupil data, and heart rate, again best from the time-domain EEG. Notably, classification worked more reliably during the second image presentation than during initial presentation (except in pupil data, where there was little difference). The frequency-domain EEG (except gamma band) also produced noisy but significant decoding results for both categories; see Supp. A.2.5 for plots of all trials and conditions. Decoding timecourses were computed for each individual participant in a stratified 10-fold cross validation; significance was determined by second level statistical testing (cluster based permutation t-test, significance level p < .05). X-axis reflects the chance level (0.5 for the used AUC metric).

**Figure 4:**
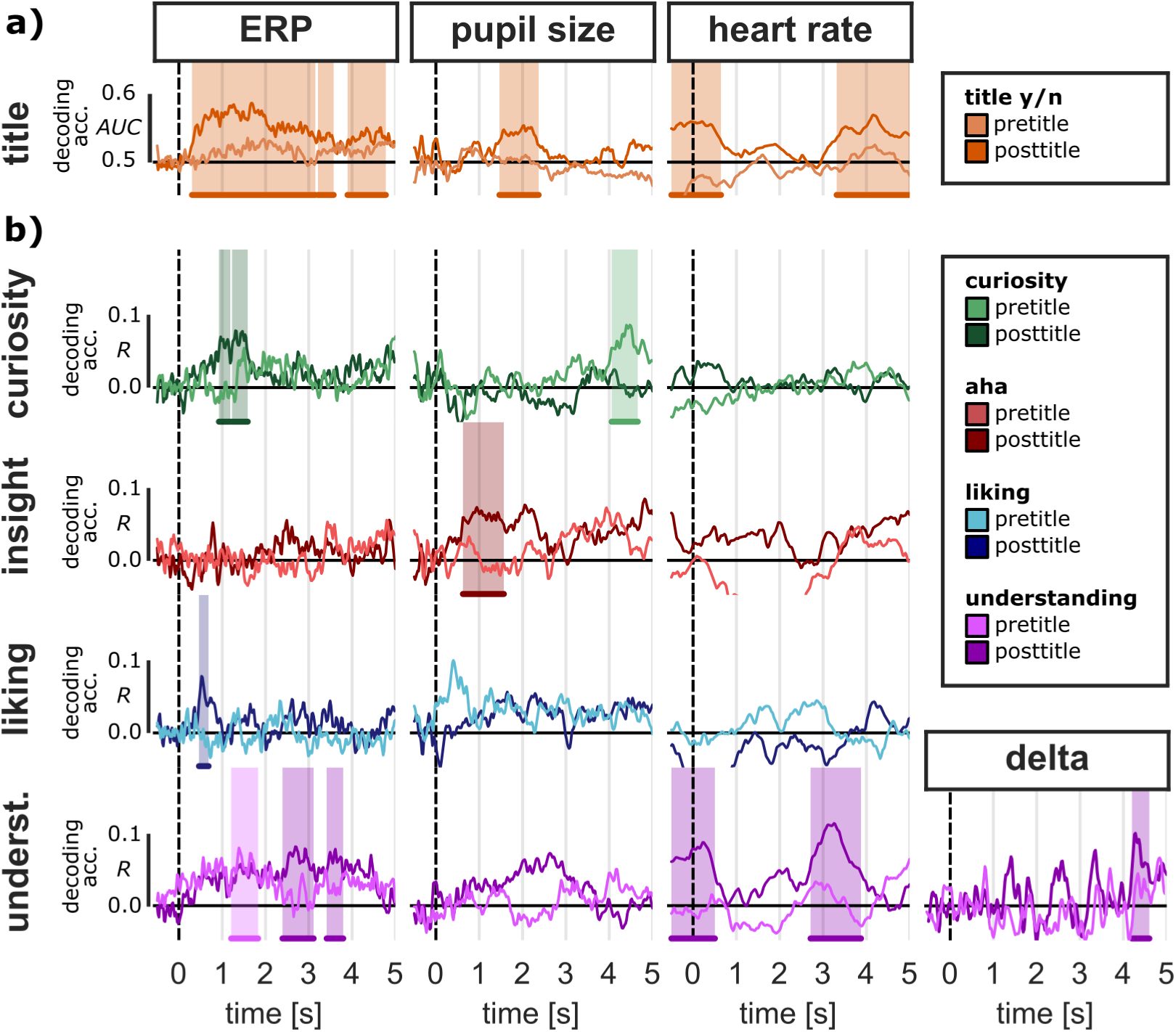
Decoding physiological correlates of meaning making (artwork trials only) **– a) Decoding the “title-effect“:** we were able to significantly decode whether a painting stimulus was paired with its real title or an uninformative dummy label (“untitled”). This worked best from time-domain EEG both during second viewing (posttitle), but also from pupil size and heart rate (heart rate during 2nd presentation appears affected from earlier processing, e.g. during title presentation, and decoding results are best before and shortly after image onset). As expected, the title condition could not be decoded from the first presentation of the images (before the title was presented). **b) Decoding trialwise ratings of curiosity for the title, *aha*, aesthetic liking, and felt understanding:** we could decode each type of rating from one or more of the recorded physiological signals. Felt understanding could be decoded best, from time-domain EEG (both in pre- and posttitle trials), from late delta band power (in posttitle trials), and from heart rate (in posttitle trials, with a timecourse similar to those for decoding the title condition, though a little earlier). Pupil dilation showed above chance decoding during post title trials too (with a timecourse similar to ERPs), but did not become significant. Curiosity for the title could be significantly decoded from the late pretitle pupil signal and the early posttitle time-domain EEG. Insight (*aha*) ratings could be significantly decoded from pupil size early in posttitle trials. Ratings of aesthetic liking showed a short peak of significant decoding in the posttitle time-domain EEG. While not becoming significant, the pupil size signal delivered generally above chance decoding with a marked peak in early pretitle trials. No other data modalities showed significant results in pre or posttitle trials; see Supp. A.2.5 for plots of all trials and conditions. Decoding timecourses were computed for each individual participant in a stratified 10-fold cross validation; significance was determined by second level statistical testing (cluster based permutation t-test, significance level p < .05). X-axis reflects the chance level (0.5 for the AUC metric in binary decoding, and 0 for the spearman R metric in continuous ratings).

#### Robust decoding of presentation, image category and title condition

The categorical experimental manipulations we applied could be detected quite robustly. Decoding the contrast between the 1st and 2nd presentation of any given image stimulus (see Fig. 3a) was reliably possible from almost all data modalities. This strongly suggests that the stimuli are not processed identically during repeated presentation; rather, cognitive processing differs in a detectable way. Whether this is caused by mere repetition suppression (cf. Kim, 2017) or other factors specific to the study such as the interim contextualization with additional information (i.e. the real or fake title, or the Mooney solution respectively) or the differing implicit or explicit tasks (e.g., generating ratings) can not be clearly determined at this point.

The presented image category (Mooney image or painting) could also be reliably decoded right from image onset from both EEG and pupil data (see Fig. 3b). Notably, except in the pupil data (see discussion) classification worked more reliably during the second image presentation than the first (i.e. higher decoding performances, and more sustained significance); this was the case for most of the investigated decoding tasks in this study.

A fuller description of the results can be found in Supp. A.2.6.

Turning to the questions more central to this study, the title condition in artworks (real title or a dummy title) could also be significantly decoded from the time-domain signals (see Fig. 4a). However, none of the EEG frequency-bands could be used to reliably decode the title condition. When decoding from ERPs during second viewing (posttitle), accuracy was the highest in a sustained but relatively early window after image onset (significant from 0.3 s onward with 2 short intermissions). Decoding from pupil size was significant in a time-window between 1.47–2.37 s in the posttitle condition. Heart rate during second viewing predicted the condition significantly in two time-windows from −0.50–0.64 s and 3.31–5.00 s, respectively. As to be expected, the title condition could not be significantly decoded during first viewing (when the title or dummy had not yet been presented) from any of the recorded data modalities.

#### Physiological signatures of subjective processing were dominated by understanding

Participant’s ratings of curiosity, liking and understanding could each be significantly decoded from ERP data at some time during the 5 s image presentation, while insight could be robustly decoded from pupil size, and understanding from heart rate (see Fig. 4b). Among these measures, felt understanding could be generally best decoded. The time-domain EEG allowed good decoding in the early pretitle trial (significant from 1.21–1.83 s after image onset). In the posttitle trial, decoding rate stayed above chance throughout the whole trial, with values similar to the pretitle trials early on (though not significant) but building up toward the end of image presentation to become significant in 2 late time-windows (2.39–3.11 s and 3.41–3.80 s after image onset). Delta band power in the late posttitle trial also significantly predicted understanding ratings (4.20-4.60 s after image onset). Heart rate in posttitle trials showed marked significant decoding for understanding in 2 time-windows (−0.50–0.50 s and 2.71–3.88 s after image onset) that were very similar to those for decoding the title condition (though slightly earlier; see above). Pupil dilation also exhibited above chance decoding, with pre- and posttitle timecourses very similar to the ERP data, though they did not reach significance.

Ratings of curiosity for the title could be significantly decoded from the late pretitle pupil signal (4.06-4.66 s after image onset) and the early posttitle time domain EEG (0.93-1.19 s and 1.23-1.59 s after image onset).

Insight (*aha*) ratings could be significantly decoded from pupil size early in posttitle trials (0.62-1.57 s after image onset).

Ratings of aesthetic liking showed a brief peak of significant decoding in the time-domain EEG signal in posttitle trials (0.46-0.68 s after image onset). The pupil signal, though not significant, generally delivered above chance decoding with a marked peak in early pretitle trials (about the first second after image onset), while in posttitle trials, this peak was absent and decoding ramped up over time.

## 4 Discussion

We investigated the relationship between curiosity, insight, felt understanding, and aesthetic liking in an attempt to unify these constructs within a coherent framework of motivated information foraging. We call this framework the “epistemic arc” of meaning making. We studied both a) the domain of behavioral ratings, to better understand the interactions of these processes and b) their (neuro)physiological correlates, to ideally gain insight into the temporal sequence of events. We studied visual art as a controlled yet naturalistic class of stimuli that can prompt meaning-making and insight, contrasted with Mooney images as an established stimulus category in the study of insight. In doing so we were able to replicate a number of earlier findings from both the behavioral literature on insight and on aesthetic valuation. Crucially, we revealed complex and partly nonlinear relations between the ratings that were shaped by the type and amount of information provided through the stimulus material. Using a time-resolved decoding approach and multi-modal physiological data (EEG, pupillometry, heart rate) we found evidence for both fast responses as well as slower processes that build up over time, though merging our results into a coherent temporal-sequence model proved difficult.

### 4.1 The relationship between ratings of curiosity, insight, felt understanding, and aesthetic valuation

The behavioral study (Exp. 1) supported most of our preregistered hypotheses: we were able to both replicate a strong effect of title manipulation on perceived understanding ratings with visual art and to establish the same (even stronger) effect of titles on *aha* ratings. We did not replicate a significant title effect on aesthetic preference ratings, though, thereby adding to an already inconclusive literature. As reported many times before (e.g. Leder et al., 2016; Vessel et al., 2018), we saw differences in aesthetic preference ratings across different stimulus categories, with generally higher ratings for figurative art than for abstract art, with semi-abstract art lying in between. This difference was even greater for perceived understanding. In *aha* ratings however, there was a trend towards slightly higher *aha* for semi-abstract art than for figurative or abstract art.

In addition to visual art we included a set of Mooney images and matching solutions. Those can cause perceptual shifts that evoke strong feelings of *aha*, and can hence act as a benchmark. Several previous studies have investigated questions similar to ours using Mooney images but not visual art (Muth & Carbon, 2013; Van de Cruys et al., 2021). As expected, *aha* ratings were higher for Mooney images than for all artwork subcategories, except some of the semi-abstract art with explanatory titles – a category somewhat similar to Mooney images as they can both lead to *perceptual* shifts after a given explanation (for example in some cubist paintings). Aesthetic preference ratings, on the other hand, were on average slightly lower for Mooney images than for art. With respect to felt understanding, Mooney images lay between figurative and abstract titled art, on par with the semi-abstract stimuli. Curiosity for a title or solution predicted subsequent *aha* (as previously reported for Mooney images; Van de Cruys et al., 2021), though our data suggests that the strength of the association depends on the information content of the title cues. While descriptive LOWESS curves suggested that this interaction might result in partly nonlinear associations, we did not find statistically significant evidence for this. *Aha* is positively associated with liking (quite linearly, as previously reported for Mooney images; Muth & Carbon, 2013) but also with felt understanding. This association between aha and understanding non-linearly interacts with the degree of abstraction of the stimuli – from a linear association in abstract art to more U-shaped relations in the other modalities (strongest in figurative art).

Other recent studies on insight have also observed nonlinear associations, thereby extending on previous findings that generally tested for linear relations only. For example, Chesebrough et al. (2023) found a U-shaped association between both perceived task difficulty and solution confidence with ratings of *aha* in a text-based analogy-solving task (both very easy and very difficult trials caused lower *aha* than the intermediate trials). This finding would be in line with our interpretation of low *aha*–high understanding trials (especially in figurative art) marking “easy” stimuli with low initial uncertainty. Orwig et al. (2025) on the other hand found that externally rated creativity of responses across 3 different text-based tasks was non-linearly associated with semantic distance of the prompt–response pairs (positive linear up to a certain threshold, after which increasing semantic distance ceased to increase aha). In our case this might be especially relevant for the interpretation of the results for evocative titles, which were explicitly selected for having a high conceptual distance to the depicted visual content and, to a lesser degree, for abstract art. While subjective ratings of *aha* are not equal to an assessment of creativity, we would expect a positive correlation. In our data we indeed see that art with purely descriptive titles (i.e. very low conceptual distance) received the lowest average *aha* ratings. Evocative titles, on the other hand, received *aha* ratings generally on par with explanatory titles, and the highest overall *aha* was for semi-abstract art with explanatory titles. This pattern seems very much in line with the findings of Orwig et al. (2025). While we could not explicitly replicate their analysis, as we did not assign a more precise stimulus-wise measure of conceptual distance between art and title, a focused reanalysis of our data could give even stronger evidence for whether this nonlinear association indeed generalizes across modalities, and where exactly the thresholds might sit for visual art. As a side note, we should keep in mind that the highest overall *aha* ratings were still given for Mooney images to which measures of semantic or conceptual distance (let alone creativity) do not easily apply. However, the perceptual shifts that can happen in trials indicated by high *aha* arguably reduce distance in the observers’ internal representation space (e.g. between a hard-to-interpret Mooney image of a cat and the participant’s internal “cat” concept). A similar case could be made for the explanatory titles, that can also trigger a state transition from high to low distance in representational space. Likewise, assigning a measure of conceptual distance for this category could also be tricky, as it depends on the prior state of the participant - distance is high if an explanation is unknown, but low if already familiar. While the cited works report nonlinear trends in the study of insight and creativity in line with our findings, we are not aware of other work that finds the shape of association within one and the same task to depend on stimulus category (i.e. our evidence for a linear association of aha and understanding in abstract art, but progressively nonlinear associations in the other image categories).

Taken together, the observed rating patterns advance our understanding of the role of insight in meaning making and aesthetic preference formation. The association of both curiosity and understanding with *aha* (especially in the high *aha* range) supports the notion that *aha* can be a valid indicator of information gain or uncertainty reduction (in line with the conclusions of Van de Cruys et al., 2021). Trial-wise curiosity ratings, in this logic, might reflect the expected *potential* for information gain – its actualization, however, depends on the additional semantic information provided, as shown by the effect of title style (i.e. the amount of newly provided semantic information) on the relationship between curiosity and *aha* after seeing the title (see fig 2b): the more uncertainty reduction was possible or expected (indicated by high curiosity ratings), the stronger the effect of the given cue and hence the association between curiosity and *aha* (with the strongest relation between curiosity and *aha* observed in Mooney images, and the weakest in untitled art).

Low felt understanding, on the other hand, might be interpreted as an indicator of remaining or residual uncertainty. The EEG decoding results support this interpretation, as understanding was best decodable towards the end of the 2nd trial (both from time-domain EEG and theta power), when all information had been brought together. High ratings of *aha* (large information gain) were consistently associated with high understanding, but depending on the stimuli (art category and title style) high understanding ratings also appeared in low *aha* trials. This was particularly the case for figurative art and Mooney images (see Fig. 2b) where low *aha* followed particularly low curiosity ratings (see also Supp. Fig. A.3). It is possible that for a good fraction of these stimuli participants were less curious about the title/solution because there was little uncertainty in the first place, and hence less potential for *aha* – a figurative painting of a lush meadow or a Mooney image that was already recognized on first sight might not make one as eager for the title or base image, while abstract and semi-abstract artworks have a relatively low likelihood of being entirely unambiguous to a viewer.

Liking ratings, on the other hand, were positively and rather linearly predicted by all other ratings – curiosity, *aha*, and understanding. These associations were largely unaffected by other factors such as art style or title style (see Fig. 2d and Supp. A.2.1). At first glance this might suggest that the role of aesthetic valuation (or at least our rating operationalization in this study) is indeed quite plain: aesthetic liking simply follows the other positively valenced judgments, while these other ratings can have a more intricate genesis. Yet, our data also contain arguments against such a simplistic causal interpretation: while we (as many groups before us) found that figurative art received on average higher aesthetic ratings than semi-abstract and abstract art, this was not the case for *aha* and curiosity – the artwork category did not matter much for *aha* ratings (with only semi-abstract art scoring higher than the others) and curiosity was even markedly lower for figurative art than for the more abstract categories. Furthermore, there was no significant effect of our title manipulation on aesthetic liking, while it substantially affected both felt understanding and *aha* (for curiosity this comparison does not make sense as the titles were presented after collecting this rating). Although all three ratings linearly predicted liking on trial-wise bases, liking does not appear to simply follow the other ratings, or the grand average patterns would be more similar. Furthermore, in the decoding timecourses liking exhibited a peak of decodability early in the trials, well before the other ratings (further discussion below). Building on our findings, future research might design intervention studies to have a closer, ideally more causal, look at these interrelations.

Above we discussed many interesting findings on the effect of information content and various other stimulus features on meaning making, but we want to emphasize that there are certainly other important factors contributing as well. Especially individual differences between participants might manifest in more complex interactions than just baseline differences. For example, a study by Wiesmann (2010) showed that different visual search strategies (as linked to art expertise or simply training) can lead to faster and more accurate object recognition in cubist paintings (i.e. more efficient extraction of information). But also positive mood was found to amplify feelings of *aha* (Chesebrough et al., 2023), and it is unclear whether this would apply equally across all conditions, or show more complex interactions.

### 4.2 Physiological correlates and the temporal sequence of information processing

In the second experiment we aimed to characterize the temporal profile of the epistemic arc by tracking fast (neuro)physiological correlates. This part of the study was more exploratory and we had only preregistered two coarse hypotheses based on previous work – namely an association between EEG alpha and gamma band activity with *aha* ratings and liking ratings. We did not find evidence to support these hypotheses. While our multivariate decoding approach was able to robustly identify the basic experimental manipulations based on time-domain EEG, frequency-domain EEG (except gamma band), heart rate and pupil data, the graded ratings proved harder to recover, and decoding worked better in the various time-domain signals than in the frequency-domain EEG.

#### Decoding from EEG

Among the ratings, felt understanding had the strongest signature in the EEG. No previous work that we are aware of has investigated felt understanding using EEG, hence our lack of any preformulated hypotheses about it. Interestingly there seemed to be an initial phase of increased decodability in both pre- and posttitle trials (roughly the first 2 seconds of image presentation, becoming significant only in the pretitle condition and hence apparently not dependent on the title information provided). In the posttitle trials however, the decoding rate did not drop after this initial phase (as in pretitle trials) but kept building up and became significant later in the trial – this would be consistent with accumulating evidence leading to a sustained understanding signal that can be captured in the data. This interpretation is further supported by the relatively similar (though not significant) decoding timecourse in the pupil trace, and the late significant peak of decoding rate in the instantaneous heart rate (see Fig. 4b). We had hypothesized understanding of a stimulus to be the desired outcome of engaging in the process of meaning making, and perhaps the most practically relevant. That felt understanding showed the strongest neurophysiological signal might be seen as support for its primacy within the epistemic arc and in neural (or at least cortical) processing.

Decoding timecourses for aesthetic liking showed a salient peak in the first second after image onset in the time-domain EEG of posttitle trials. This offers further neurophysiological evidence for the presence of a very fast aesthetic response component, consistent with previous observations that some ERP components might be linked to aesthetic processing (Oliver-Rodríguez et al., 1999; Marzi & Viggiano, 2010; Werheid et al., 2007; de Tommaso et al., 2008; Augustin et al., 2011; Wang et al., 2012; Jiang & Cai, 2013; Noguchi & Murota, 2013; Ma et al., 2015; Bölte et al., 2017; Righi et al., 2017; Carbon et al., 2018). The early peak in the pupil data of pretitle trials, though not significant, might further back this claim. Importantly, liking ratings could only be decoded from the EEG of the 2nd presentation of images, post title, but not from the initial presentation. Apparently, the early cortical response to the image differed between 1st and 2nd presentation (see also grand average response in Supp. Figs. A.4, A.8 and discussion below) and this difference was relevant in decoding aesthetic liking. This might speak against a dominant role of low-level image features on final liking ratings. It is possible that the EEG response is present in the pre-title trial too (see pupil response) but overshadowed by noise and the far greater temporal distance to the rating. However, it could also be that the liking response collected after the second viewing deviated from a hypothetical liking rating that would have been given after the initial viewing. Given our behavioral findings we might expect the title intervention and related meaning making processes to cause such divergence, or evolution of liking, and other factors, like the repeated or longer exposure (cf. Zajonc, 1968; Burgess & Sales, 1971), unknown factors, or simply noise in the ratings, might also contribute to the lack of decoding in the first presentation. Unfortunately, our data does not allow for a more thorough investigation of this effect, as we lack a measure of change in liking from before to after seeing the title – while collecting such a measure would be methodologically challenging, future research could follow up on this. We find it noteworthy that we did not see significant decoding of liking later in the trials – previous EEG (Strijbosch et al., 2021) and behavioral findings (Brielmann & Pelli, 2017), as well as our own behavioral observation in this study would lead us to expect also later, more sustained contributions to a final liking response. Further research should validate or rectify these null findings.

The early peak for liking was not immediately mirrored for *aha*; while *aha* could be significantly decoded from posttitle trials (in the pupil signal) it became significant *after* the peak for liking. Perhaps this finding is misleading, though: the pupil response is a comparatively slow signal (and might be unreliable immediately after image onset; see discussion below), so it seems possible that the actual *aha* causing the pupil response happened slightly earlier, around the same time range as the liking response. Still, these findings provide no clear support for the hypothesis that moments of aha causally lead to increased liking.

Curiosity proved a very interesting signal: while the significant cluster in the late pretitle pupil signal (immediately prior to the rating) might be unsurprising, the robust decodability from time-domain EEG in posttitle trials certainly is. Participants registered their rating several seconds prior to this timepoint, so it cannot be a case of reverse attribution. Furthermore, they had to process an entirely different stimulus in the meantime (the title cues). If there is in fact a distinct state of cortical activity that curiosity ratings accurately reflect, our findings indicate that this activity is sustained over several seconds from the initial presentation of the image, over seeing and processing the title cue, and well into the posttitle trial. This would be an important finding, as these altered states could then also affect the slower cortical processing and contextualization of the new title information with the repeated image stimulus. The peak in decoding in the posttitle trial might perhaps mark a moment of “closure” for a nested epistemic arc, one that began with the presentation of the first stimulus and ended with the internal combination of the title during the 2nd image viewing. The fact that the curiosity rating can be significantly decoded only after the peak in liking ratings (also in the time-domain EEG) is also relevant, as it makes it appear less likely that liking is a direct consequence of curiosity, despite their positive linear relation in the rating space (see discussion above).

In general, liking, curiosity and *aha* ratings proved difficult to recover from the EEG data. There are several potential reasons why this might be the case. Many of the previously proposed EEG biomarkers for insight and/or aesthetic valuation are patterns in the frequency-domain EEG. In this light, it seems surprising that in our data the EEG band power was such a poor feature to predict the ratings. This does not necessarily mean that these correlates are not present, though. Poor decoding might also be caused by poor temporal synchronicity of physiological responses across trials: with the current decoding pipeline, short bursts of activity, as proposed for *aha* (especially gamma band activity; Oh et al., 2020), would be most reliably detected if temporally aligned (i.e. immediately time-locked to the stimulus) and otherwise easily lost in noise. Previous EEG studies on insight typically addressed this issue by reverse time-locking analyses to a participant’s button press indicating moments of insight – we had decided against this procedure for our study, as 1) it might disrupt the flow of the paradigm, 2) we were interested in a graded rating, and 3) we explicitly wanted to relate this continuous rating of *aha* to other ratings like curiosity and liking. While we knew in advance that there may not be clear alignment for these internally triggered processes (insight or aesthetic liking can evolve anytime or not at all), we nevertheless expected that the alignment would be good enough to detect more sustained or partly overlapping correlates. Other analysis approaches might be better suited to find asynchronous patterns and could be tried in the future. Interesting candidate pipelines might be based on unsupervised learning (e.g. Schneider et al., 2023), temporal deconvolution (e.g. Ehinger & Dimigen, 2019), or simply decoding with flexible timelags. Another approach could be to further increase the signal to noise level in the recorded data. The most effective option would be to use intracranial recordings, as done by Lopez-Persem et al. (2020); however, this would also require transitioning to a smaller and very different sample population (i.e. patients). On the other hand, more phasic frequency responses should be detectable in our design and this seems to be the case in the alpha, beta, and even gamma band for some of the categorical decoding targets (1st vs 2nd presentation and artwork vs. Mooney; see Supp. A.2.5).

The categorical experimental factors were more easily decodable. Particularly the fact that the title effect could be decoded from time-domain signals of posttitle trials (see Fig. 4a) is more than a sanity check and very encouraging. This contrast, in which the only difference was in the text information shown prior to image onset, is reminiscent of work on contextual priming (e.g. Wirth et al., 2008; Horner & Henson, 2012) though differences in the stimulus material and priming procedures prevent a direct comparison. Different priming stimuli are thought to moderate the ease of processing gained upon repeated presentation (either further facilitating, or interfering) creating a further conceptual overlap with work on repetition suppression. Our title cues can certainly be seen as primes, and both processing fluency (Reber et al., 1998) as well as repeated exposure (though inconclusively; see e.g. Zajonc 1968; Burgess and Sales 1971, but also Park et al. 2010) are thought to play a role in aesthetic judgment. But would the titles facilitate or complicate processing of a repeated visual art stimulus? Our behavioral findings suggest that this would most likely depend on the specific combination of image stimulus and title. To further widen the picture, we again note that cortical activity related to curiosity carried over into the 2nd image presentation regardless of which type of title stimulus was presented, speaking against a monopolistic influence of the priming title cues. The interplay between repeated presentation effects, ease of processing, and additional information in processing visual art (and meaning making more generally) is indeed very interesting – we might have opened a view into these intricate processes and their interrelations, but further research is necessary to progress our understanding.

Interestingly, pre- vs post-title presentation was better decodable in art trials than in Mooney images (judging by length of significant time windows; see Fig. 3). As discussed above, one explanation could be that repetition effects are particularly strong in art. Alternatively, Mooney images might show particularly weak repetition suppression – either the prominent low level visual features of Mooney images might govern physiological responses, allowing for relatively smaller effects of repeated presentation, or the perceptual shift that can happen after understanding the solution effectively renders the second presentation of a Mooney image a different stimulus, resulting in less repetition suppression. Previous work has in fact linked the strength of repetition suppression to surprise and expectations (Summerfield et al., 2008, 2011). However, qualitatively the decoding timecourses do not differ that much and it is also possible that we simply observed false negative findings as there were fewer trials in the Mooney-only condition than in the art condition, resulting in lower statistical power (16 Mooney and 48 artworks per participant).

#### Decoding from pupillometry and heart rate

As expected, pupillometry turned out to be a very promising data modality: it could be used to reliably decode the categorical factors (1st or 2nd presentation, Mooney or artwork), showed significant time windows in the title effect, curiosity and *aha* ratings, and non-significant but above chance decoding with interesting temporal profiles for liking and understanding (see Figs. 3 and 4). While findings for aesthetic liking showed no significant effects, we saw above chance decoding throughout most of the trial and a marked peak of decodability in the first second of pretitle trials (not present in posttitle trials). If true, this would be further evidence for a first and fast component in preference formation (cf. posttitle ERP data). Importantly, pupil size was also the only recorded data modality that we could significantly predict *aha* ratings from (besides a short cluster in theta band EEG of the differential signal; see Supp. A.2.4). Perhaps the pupil, as a comparatively slow or inert signal (though not as slow as heart rate), is less affected by the lack of synchronization across insight moments. This might be consistent with the decoding timecourse in the differential signal (see Supp. A.2.4 and Supp. Fig. A.8) that exhibited a slow monotonic increase over the duration of the trial. In posttitle trials there was a significant window of decoding for *aha* early in the trials, becoming significant about half a second after image onset. It is interesting to compare this with the decoding of curiosity ratings, which showed marked significant effects late in the pretitle trials, in the last second before image offset. Taken together this pattern makes the pupil a very promising candidate for future research on the interplay of curiosity, insight and understanding in information processing (and perhaps also for work on preference formation).

Unfortunately, pupil size *early in the trial* (about the first second after image onset) might not be a reliable signal for higher-level cognitive processes. In our data, we see that no rating nor the title condition could be decoded in the early part of trials. At this time the pupil involuntarily adjusts to luminance changes on the screen (see grand average pupil traces, in Supp. Fig. A.4). Note however, that the interesting non-significant pattern in pretitle liking ratings fell exactly into this temporal profile and the significant decodability of *aha* in posttitle trials begins immediately subsequent, indicating that the pupil readjusts to a different dilation level depending on *aha*. The contrast between Mooney images and artwork, on the other hand, could be particularly well decoded in this time window; this is not surprising given that the Mooney images are high contrast black/white images with a large fraction of pure white pixels – hence the pupil should adapt stronger to those images, compared to artworks which are on average less luminous. Further supporting this interpretation, the 1st vs 2nd presentation *within Mooney images* could not be significantly decoded from the early pupil data (only later in the trial) while this worked reliably for artwork stimuli and from the other data modalities like EEG (see Fig. 3 and Supp. A.2.5). These patterns are consistent with the explanation that low level stimulus features were driving the pupil’s response in this part of the trials.

Heart rate, a signal closely linked to the autonomous nervous system via sympathetic/parasympathetic pathways, was another candidate data modality to be affected by aesthetic processing. As in previous work we observed a marked heart rate deceleration following the onset of visual stimuli (Palomba et al., 1997; Welke & Vessel, 2022). This deceleration was stronger for the first (pretitle) presentation than the second presentation, mainly because in the second presentation heart rate started on an already lower level (see Supp. A.2.3). In pretitle trials we could not decode any category or rating from heart rate, but in posttitle trials this was possible. Due to the sluggishness of the signal, it is likely that events during the preceding pretitle trial or the title revelation spill over into recordings during the posttitle trials (importantly, unlike all other data modalities, we did not baseline correct the heart rate data). This might explain why we could reliably decode events like the image category or title condition, already in the pre-stimulus period of the second image presentation – all the necessary information was there before. The decodability of the graded understanding ratings in the prestimulus period of posttitle trials is a more interesting finding. There was significant understanding signal in the ERP data early in pretitle trials, and it seems possible that a feeling of understanding could affect the heartrate; in that interpretation the first significant peak of decodability (around stimulus onset) might reflect understanding derived from the pretitle image alone (plus a long time lag) while the second peak might reflect additional understanding gained from the titles (again plus time lag). The decrease of decodability between peaks might reflect disruptions from the new stimulus onset. Yet, it also seems possible that this is an overinterpretation of the results. Importantly we did not see decodability of *aha* which might seem an equally or more plausible driver of heart rate changes than understanding alone, even though the slow heart rate signal should be more robust to a lack of event synchronization than our other measures (see discussion above).

### 4.3 Conclusion

Engaging with visual art and additional text-based information such as titles or curatorial notes is a paradigmatic case of “pleasure from understanding”, in which an information gap triggers curiosity, attempts at sense making, and potentially, pleasure derived from the resulting insight. By studying the phenomena of curiosity, insight, felt understanding and aesthetic appeal in a single paradigm, we provide a new perspective on the nature and subprocesses composing this epistemic arc. In Exp. 1 we established a paradigm to robustly evoke insight with visual art and gained new understanding of the intricate interplay between the feelings curiosity, insight, felt understanding, and aesthetic liking, revealing partly nonlinear relations shaped by the type and amount of information a specific stimulus provides. In Exp. 2 our analysis of multi-modal physiological correlates, despite not confirming all our *a priori* hypotheses, opened new angles to further investigate the topic in the future, leading to a more mechanistic understanding of meaning making and its role in the aesthetic evaluation process. Though much remains to explore, our study provides a new window into core behavioral and physiological properties of motivated information foraging.

## 5 Additional information

### Data and code availability

The experimental design and hypotheses reported here were preregistered (Exp. 1: https://doi.org/10.17605/OSF.IO/PAGVH and Exp. 2: https://doi.org/10.17605/OSF.IO/6KW9N). The dataset for statistical analysis, as well as python scripts replicating the results, and python scripts used in data collection were made available in a public online repository (https://osf.io/j42g5). The stimulus set is entirely in the public domain and included in the online repository; all artwork stimuli are further depicted in the online Supplement of this paper. An earlier stage of this article was uploaded to a preprint server (https://www.biorxiv.org/content/10.1101/2025.05.15.654230).

## Acknowledgments

This work was supported by the Max Planck Society.

## Author statement (CRediT)

DW: Conceptualization, Methodology, Investigation, Formal Analysis, Software, Data curation, Project administration, Visualization, Writing - Original Draft, Writing - Review & Editing EV: Conceptualization, Writing - Original Draft, Writing - Review & Editing, Resources, Supervision, Funding acquisition

## A Supplementary Material

### A.1 Material & Methods

#### A.1.1 Participant demographics

See Tabs. A.1 and A.2 for full sample demographics.

**Table A.1:** Categorical demographic factors.

| Variable | Exp. 1 | Exp. 2 |
| --- | --- | --- |
| n | 56 | 49*** |
| Sex |  |  |
| Female | 37 | 28 |
| Male | 19 | 19 |
| Other | – | 2 |
| Handedness* |  |  |
| Right | 49 | 41 |
| Left | 3 | 8 |
| Ambidextrous | 3 | – |
| N/A | 1 | – |
| Eye dominance* |  |  |
| Right | 7 | 28 |
| Left | 14 | 17 |
| No dominance | 4 | 4 |
| N/A | 31 | – |
| Highest degree of education** |  |  |
| Realschulabschluss | 1 | – |
| Hochschulreife | 25 | 26 |
| Hochschulabschluss | 23 | 23 |
| Promotion | 6 | – |
| N/A | 1 | – |
| Mental disorder |  |  |
| No | 51 | 38 |
| Yes | 5 | 11 |
| Neurological disorder |  |  |
| No | 54 | 49 |
| Yes | 2 | 1*** |

*Note:* \* in Exp. 1 handedness and eye dominance were collected as self-reports, while in Exp. 2 the FLANDERS questionnaire was used to assess handedness, and the PORTA test was used to assess eye dominance; \*\* German qualification levels – RS: finished school, HSR: qualified for/currently in higher education, HSA: bachelor/master degree, PR: PhD ; \*\*\* in Exp. 2, data from N=50 participants were collected, but 1 participant with neurological disorders was excluded from the sample.

**Table A.2:** Continuous demographic factors.

| Variable | n | M | SD | min | max | 25% | 50% | 75% |
| --- | --- | --- | --- | --- | --- | --- | --- | --- |
| Exp. 1 |  |  |  |  |  |  |  |  |
| Age | 55 | 40.0 | 16.97 | 20 | 82 | 25 | 35 | 51 |
| Years of education | 53 | 17.4 | 3.89 | 7 | 26 | 15 | 18 | 20 |
| BFI (range 3-15) |  |  |  |  |  |  |  |  |
| extraversion | 56 | 9.36 | 2.178 | 4 | 15 | 8 | 9 | 11 |
| open mindedness | 56 | 11.52 | 1.935 | 8 | 15 | 10 | 12 | 13 |
| agreeableness | 56 | 11.00 | 2.132 | 5 | 15 | 9 | 11 | 13 |
| conscientiousness | 56 | 10.93 | 2.213 | 5 | 14 | 9 | 10.5 | 12 |
| negative emotionality | 56 | 8.59 | 2.661 | 3 | 14 | 7 | 9 | 11 |
| AReA (range 0-4) |  |  |  |  |  |  |  |  |
| total | 56 | 2.02 | 0.692 | 0.64 | 3.36 | 1.57 | 2.07 | 2.5 |
| aesthetic appreciation | 56 | 2.60 | 0.736 | 1 | 4 | 2.25 | 2.75 | 4 |
| intense experience | 56 | 1.50 | 0.831 | 0 | 3.5 | 1 | 1.5 | 3.5 |
| creative behavior | 56 | 1.15 | 0.938 | 0 | 3 | 0.25 | 1.33 | 3 |
| BPS (range 8-56) | 56 | 19.71 | 8.329 | 8 | 41 | 13 | 17 | 25 |
| PANAS (range 10-50) |  |  |  |  |  |  |  |  |
| positive | 56 | 29.38 | 6.323 | 17 | 46 | 14 | 18 | 21 |
| negative | 56 | 12.97 | 4.311 | 10 | 31 | 6 | 6 | 7.5 |
| Exp. 2 |  |  |  |  |  |  |  |  |
| Age | 49 | 28.0 | 7.17 | 18 | 52 | 24 | 26 | 30 |
| Years of education | 48 | 17.3 | 3.90 | 8 | 27 | 15 | 17 | 20 |
| BFI (range 3-15) |  |  |  |  |  |  |  |  |
| extraversion | 49 | 9.47 | 2.256 | 4 | 13 | 8 | 9 | 11 |
| open mindedness | 49 | 11.96 | 1.707 | 7 | 15 | 11 | 12 | 13 |
| agreeableness | 49 | 10.82 | 1.997 | 7 | 15 | 9 | 11 | 12 |
| conscientiousness | 49 | 9.80 | 2.491 | 5 | 14 | 8 | 10 | 12 |
| negative emotionality | 49 | 9.29 | 2.458 | 4 | 14 | 8 | 10 | 11 |
| AReA (range 0-4) |  |  |  |  |  |  |  |  |
| total | 49 | 1.93 | 0.624 | 0.64 | 2.93 | 1.43 | 2 | 2.43 |
| aesthetic appreciation | 49 | 2.47 | 0.619 | 1 | 3.62 | 2 | 2.5 | 3 |
| intense experience | 49 | 1.30 | 0.802 | 0 | 2.75 | 0.75 | 1.25 | 1.75 |
| creative behavior | 49 | 1.26 | 1.037 | 0 | 3.33 | 0.33 | 1.33 | 2 |
| BPS (range 8-56) | 49 | 21.86 | 7.979 | 9 | 45 | 16 | 21 | 26 |
| Caffeine intake (in mg/kg)* | 48 | 1.06 | 0.923 | 0 | 3.61 | 0.34 | 0.89 | 1.50 |
| PANAS (range 10-50) |  |  |  |  |  |  |  |  |
| positive | 49 | 25.80 | 6.423 | 12 | 41 | 22 | 26 | 30 |
| negative | 49 | 12.35 | 3.345 | 10 | 27 | 10 | 11 | 13 |

Note: \* caffeine intake was only assessed in Exp. 2

#### A.1.2 Full task description (in German)

Willkommen zu unserer Studie!

Bitte lesen Sie sich die Folgenden Instruktionen gründlich durch. Es ist einiges an Text, also beißen Sie sich nicht fest - am Ende werden die wichtigsten Punkte nocheinmal kurz zusammengefasst. Sobald Sie mit einer Seite fertig sind, können Sie eine beliebige Taste drücken, um zur nächsten zu gelangen.

Viel Vergnügen!

——

In dieser Studie untersuchen wir, inwiefern die Titel von Kunstwerken zu ihrem Verständnis beitragen können. Dabei interessiert uns vor allem ein Phänomen, das wir „Aha! Erlebnis” nennen – das plötzlich auftretende Gefühl etwas verstanden zu haben (eine plötzliche Einsicht).

Stellen Sie sich z.B. vor, Sie betrachten ein abstraktes Kunstwerk, das Sie beim besten Willen nicht einordnen können, und überhaupt nicht verstehen worum es geht. Verwirrt wandert Ihr Blick zum Titel, und plötzlich macht es “Klick“: Sie verstehen auf einmal die Intention des Bildes, oder was die Künstlerin damit vielleicht ausdrücken möchte.

Oder denken Sie an eine kubistische Komposition (Alltagsgegenstände oder Personen, die in abstrakte Formen zerlegt dargestellt werden) – diese können schwierig zu erkennen sein, aber wenn Ihnen der Titel erklärt, was abgebildet sein soll, erkennen Sie plötzlich auch die Konturen oder ein Muster, das das Motiv für Sie zusammenhängend hervortreten lässt.

——

Auch in darstellender Kunst in der die abgebildeten Objekte und Personen klar zu erkennen sind, können solche Effekte auftreten: wenn zum Beispiel ein historisches, religiöses oder mythologisches Thema dargestellt wird, das Ihnen zwar bekannt ist, welches Sie im dargestellten Bild aber nicht sofort erkennen: hier kann der Titel dazu führen, dass das Thema für Sie offensichtlich wird und sich plötzlich ein erweiterter Bedeutungshorizont erschließt. Ähnlich könnte es bei surrealistischen Darstellungen sein, deren Titel auf abstrakte Konzepte oder Trauminhalte verweisen, die mit dem dargestellten vermeintlich sogar in Widerspruch stehen.

Dies sind nur ein paar greifbare Beispiele, um Ihnen zu verdeutlichen, worum es uns in dieser Studie geht. Wichtig ist: Wir interessieren uns nicht für eine bestimmte Art von „aha! Erlebnis”, sondern für jegliche Form von plötzlich empfundener Einsicht! Manche Bilder haben zum Beispiel gar keinen Titel; dennoch kann auch ein Bild für sich genommen evtl. ein Aha! Erlebnis auslösen. Es ist uns auch nicht wichtig, ob Ihr Verständnis “richtig” war, oder sich hinterher vielleicht als falsche Fährte heraus stellen würde. Genauso könnten die Bilder oder Ihre Titel z.B. auch persönliche Erinnerungen auslösen, die irgendwie in Verbindung damit stehen und Ihnen eine plötzliche „persönliche” Einsicht verschaffen, die nur Sie selbst haben können.

——

Um ihnen zu verdeutlichen, wie sich solche „Aha! Erlebnisse” anfühlen können, werden wir im Verlauf des Experimentes immer wieder “Camouflage” Bilder einstreuen: Abstrakte Muster aus schwarzen und weißen Flächen, die von Fotografien abgeleitet werden.

Oft ist nur schwer oder überhaupt nicht zu erraten, was das zugrunde liegende Bild darstellt, aber sobald Sie das originale Foto gesehen haben, werden Sie es danach im schwarz-weiß-Muster erkennen. Diese Form der Auflösung eines „visuellen Rätsels” kann erwiesenermaßen starke Aha! Erlebnisse auslösen.

Auf der nächsten Seite sehen Sie ein Beispiel.

——

[Mooney image example]

——

Nocheinmal, wir möchten Sie damit nicht darauf festlegen, im Falle der Kunstwerke ausschließlich auf „Aha! Erlebnisse” zu achten, die denen mit den Camouflage Bildern gleichen – uns interessiert jede Form von plötzliche empfundener Einsicht - sei es in die Identität des Dargestellten, die tiefere Bedeutung des Bildes, oder etwas ganz anderes; zb eine persönliche Erkenntnis.

——

Dazu werden wir ihnen gleich eine Reihe von Bildern zeigen (Gemälde und „Camouflage” Bilder). Jedes Bild wird für 5 Sekunden zu sehen sein und dann ausgeblendet. Als nächstes werden wir Ihnen dann den Titel des Werkes, bzw. im Falle der Camouflage Bilder das zugrunde liegende Foto, zeigen. Im Anschluss blenden wir erneut das ursprüngliche Bild ein. Die Titel, die wir einblenden sind die tatsächlichen Titel der Gemälde. Auch einige unbetitelte werden dabei sein.

Nach dieser Sequenz bitten wir Sie jeweils, auf einer stufenlosen Skala anzugeben, wie stark Sie ein Aha! Erlebnis empfunden haben.

——

Die Stärke der Empfindung kann durchaus variieren – von einem vagen Gefühl, vielleicht etwas verstanden zu haben, das man aber gar nicht genau ausdrücken oder beschreiben könnte, bis zu einem glasklaren und expliziten Einsicht ist alles möglich.

Außerdem bitten wir Sie, anzugeben wie sehr das Bild Sie ästhetisch angesprochen hat, und inwiefern Sie das Gefühl haben, das Bild verstanden zu haben. Das werden wir im Folgenden genauer erklären.

——

Mit der Frage „Wie ästhetisch ansprechend fanden Sie das Bild?” möchten wir einen Eindruck Ihrer spontanen persönlichen Vorliebe für das Kunstwerk gewinnen.

Auch hierbei gibt es keine richtigen oder falschen Antworten: Gemälde lassen sich zwar möglicherweise nach einer ganzen Reihe verschiedener Kriterien beurteilen, etwa anhand des Stils, der Farben, oder der technischen Perfektion, und Sie können eine Bandbreite von Reaktionen von “schön”, über “sonderbar”, bis “hässlich”, von “bewegend” bis “langweilig” usw. auslösen. Aber auch traurige oder auf den ersten Blick hässliche, grob gearbeitete Bilder können Einen unter Umständen stark ansprechen. Wir möchten ihre ganz persönliche Reaktion erfahren: Wie sehr spricht Sie das abgebildete Kunstwerk an, egal aus welchen Gründen?

——

In der dritten Frage interessiert uns, inwiefern Sie denken, den Sinn des Bildes oder die Intention der Künstlerin verstanden zu haben.

Sie könnten meinen, dass diese Frage sehr ähnlich zur Frage nach dem „aha! Erlebnis” ist, aber das muss nicht der Fall sein: wie beschrieben interessiert uns im Falle des „aha!” die Stärke einer plötzlichen Empfindung – diese muss nicht unbedingt zu einem tatsächlichen und nachhaltigen Verständnis des Bildes beitragen, und kann sich im nachhinein als falsche Fährte erweisen – dennoch war die Empfindung da. Oder das Bild war für Sie so einfach zu verstehen, dass Sie es bereits ohne den Titel verstanden haben – in diesem Fall tritt vielleicht kein besonders starkes „aha! Erlebnis” auf.

——

Noch vor diesen drei Fragen bitten wir Sie jeweils darum, anzugeben, wie neugierig Sie auf den Titel des Bildes sind (bzw auf die zugrunde liegende Fotographie im Falle der Camouflage Bilder). Diese Frage erscheint bereits nach der erstmaligen Präsentation des Bildes, kurz bevor der Titel eingeblendet wird. Alternativ hätten wir fragen können, wie sehr Sie sich wünschen, den Titel zu erfahren.

——

All diese Bewertungen werden Sie mithilfe eines Schiebereglers abgeben.

Bitte beantworten Sie die Fragen jeweils, indem Sie den Schieber auf der abgebildeten stufenlosen Skala mit den Polen + und - entsprechend ihrem spontanen Urteil platzieren. Dazu bewegen Sie die Maus auf-bzw. abwärts (das kann anfangs etwas ungewöhnlich erscheinen) und bestätigen die endgültige Position mit einem Mausklick. Der Schieber rastet dabei ein und wechselt die Farbe. Kurz darauf rücken Sie zum nächsten Schritt vor.

Auf der nächsten Seite wird ein Antwortslider wie oben beschrieben eingeblendet. Bitte beantworten Sie die Testfrage, um weiter geleitet zu werden!

——

[Example of slider to exercise using it]

——

Sie werden nun einige Trainingsdurchläufe machen, um sich mit der Aufgabe vertraut zu machen, bevor die eigentlich Datenerhebung beginnt.

Auf der Folgenden Seiten fassen wir das wichtigste zu den Fragen noch einmal zusammen.

——

Frage zu Neugier: Bitte geben Sie hier an, wie neugierig oder gespannt Sie auf den Titel des Kunstwerkes (bzw. auf die Auflösung des Camouflage Bildes) sind.

Frage zu Aha! Erlebnis: Bitte geben Sie an, wie stark ihr Aha! Erlebnis war – es handelt sich um ein plötzlich auftretendes Gefühl etwas verstanden zu haben (eine plötzliche Einsicht).

Frage zu Ästhetische Präferenz: Bitte geben Sie hier an, wie sehr das Kunstwerk (Bild und Titel) Sie ästhetisch angesprochen hat; egal aus welchen Gründen.

Frage zu Verständnis: Bitte geben Sie an, wie sehr Sie den Eindruck haben, das Bild oder die künstlerische Intention verstanden zu haben. Dies kann sich durchaus von der Stärke des Aha! Erlebnisses unterscheiden.

Es gibt keine richtigen oder falschen Antworten. Bitte versuchen Sie, Ihren spontanen Eindruck möglichst genau widerzugeben. Sie brauchen sich auch nicht viel Zeit nehmen, um Ihre Antwort im feinsten Detail einzustellen: folgen Sie am besten Ihrer spontanen Intuition.

——

Zwei Dinge bitten wir Sie dabei noch zu beachten:

Da wir Ihre Augenbewegungen aufzeichnen, ist es sehr wichtig für uns, dass Sie während der Präsentation der Bilder versuchen möglichst wenig zu Blinzeln. Während Sie die Fragen beantworten, oder in den kurzen Pausen zwischen den größeren Blöcken können Sie das jedoch gerne nachholen.

Außerdem wird vor jedem Bild für zwei Sekunden ein Fixationskreuz in der Mitte des Bildschirms eingeblendet. Bitte versuchen Sie, Ihren Blick in dieser Zeit auf dem Kreuz zu fixieren - sobald das Bild eingeblendet wird, und das Kreuz verschwindet, können Sie Ihren Blick dann frei schweifen lassen.

Sobald Sie bereit sind, mit dem Training zu beginnen, drücken Sie bitte eine Taste.

——

[Training trials]

——

Das Training ist zu Ende. Wir hoffen, Sie haben nun ein gutes Gefühl dafür, wie es abläuft.

Wenn Sie noch Fragen haben sollten, können Sie diese bei der Kalibrierung des Eye Trackers mit dem/der Versuch- sleiter/in erörtern.

——

- Kalibrierung des Eyetrackers -

Zuerst Kalibrieren wir den Eye tracker. Legen Sie Ihr Kinn dafür auf der Kinnstütze ab. Versuchen Sie eine möglichst bequeme Position einzunehmen, die Sie ohne zu verkrampfen über eine längere zeit aufrecht halten können.

Auf dem Bildschirm wird gleich ein Auswahlmenü erscheinen. Sie müssen nicht aktiv werden, die Auswahl treffen die Versuchsleiter. Während der Kalibrierung wird ein Punkt auf dem Bildschirm eingeblendet, der zu verschiedenen Positionen auf dem Bildschirm springt. Ihre Aufgabe ist es, dem Punkt mit den Augen zu fokussieren und zu verfolgen.

Drücken Sie eine Taste, um mit dem Hauptteil der Studie zu beginnen!

#### A.1.3 Stimulus subselection for EEG study

Besides for the main analyses, the rating data from our online study (Exp. 1) were used to subselect the stimulus set for the subsequent EEG study (Exp. 2). A reduction of overall stimulus count was necessary in order to have all participants see the same stimuli. Subselection was based on the magnitude of the title effect on aha for each image (extracted as the random-effect slope coefficients of an LMM; the details of the procedure are described in the methods). Fig. A.1 shows the ranked slope coefficients - we selected the top 16 images for each stimulus category (figurative art, semi-abstract art, abstract art, Mooney image). While there was a high variance in the slope coefficients across all individual stimuli (with a few even showing *lower* aha ratings when the title was presented) the selected image subsets for Exp. 2 have high average coefficients on a comparable level (Mooney: average coefficient = 38.6; abstract art: average coefficient = 25.7; semi-abstract art: average coefficient = 31.9; figurative art: average coefficient = 27.2). The final stimulus sets for Exp. 1 (partly overlapping sets across groups of participant) and Exp. 2 (fully shared across participants) are depicted below.

**Figure A.1:**
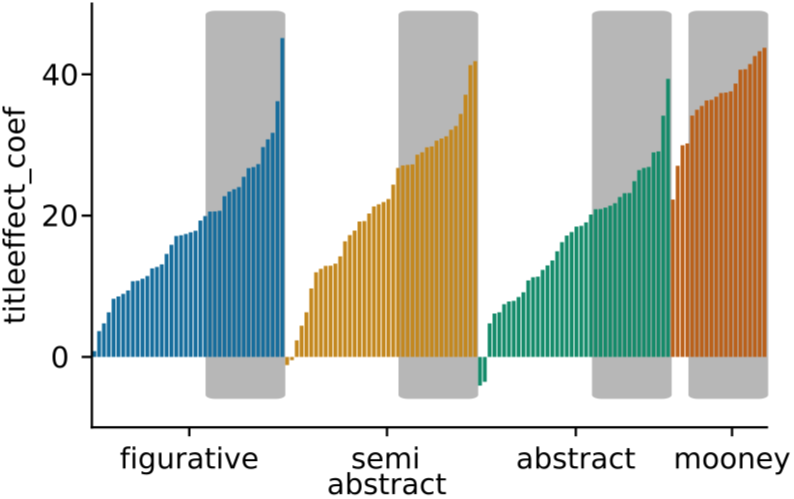
Stimulus subselection for EEG study. gray boxes indicate the selected stimuli (16 per category)

#### A.1.4 Artwork stimuli

Depictions of the final stimulus sets for Exp. 1 (partly overlapping sets across groups of participant) and Exp. 2 (fully shared across participants) are depicted below. All stimuli are in the public domain and the full stimulus set in high resolution (as used in the study) can be downloaded from our OSF online repository under https://osf.io/dafyv/ and https://osf.io/s7jtx/.

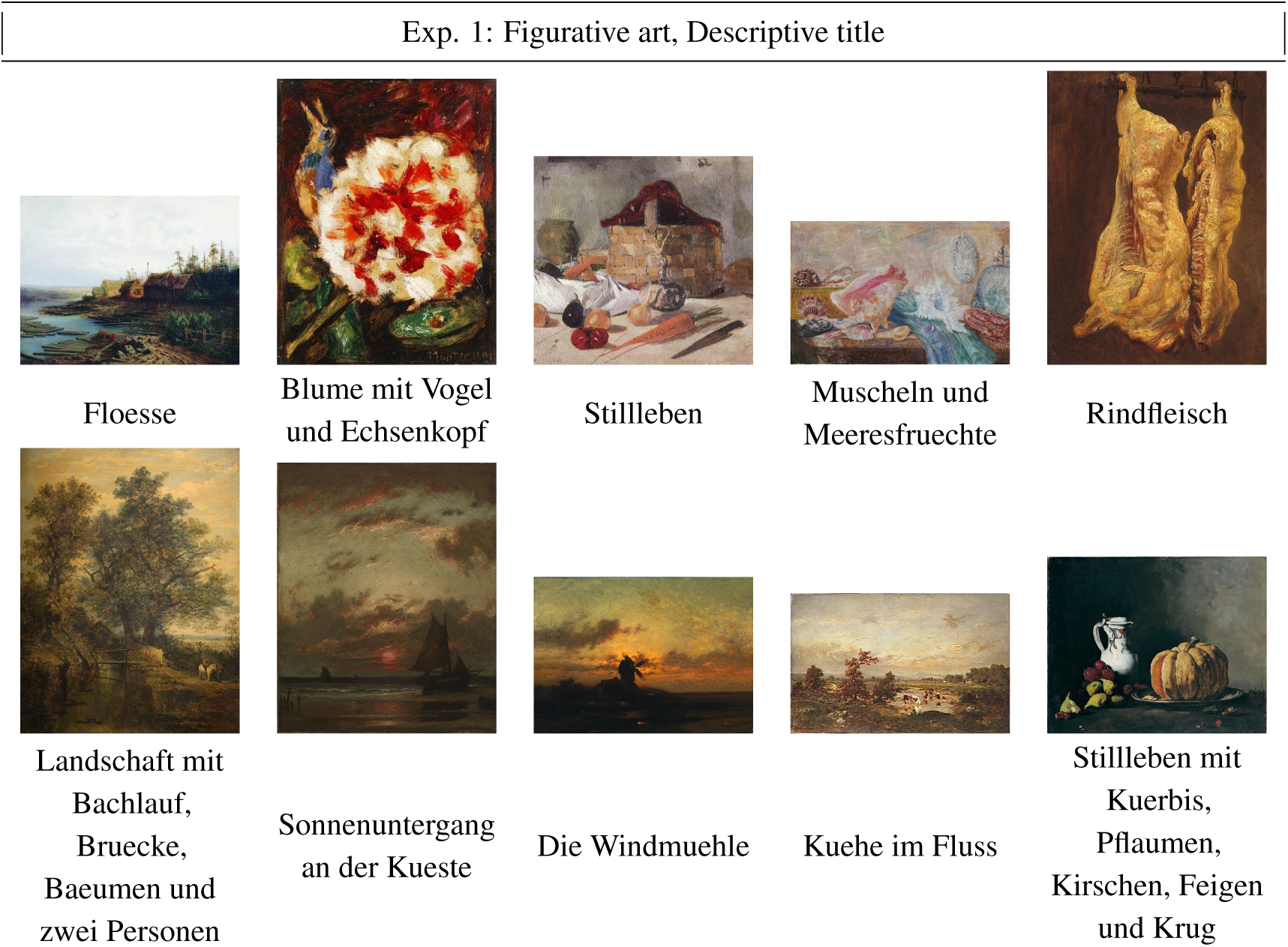
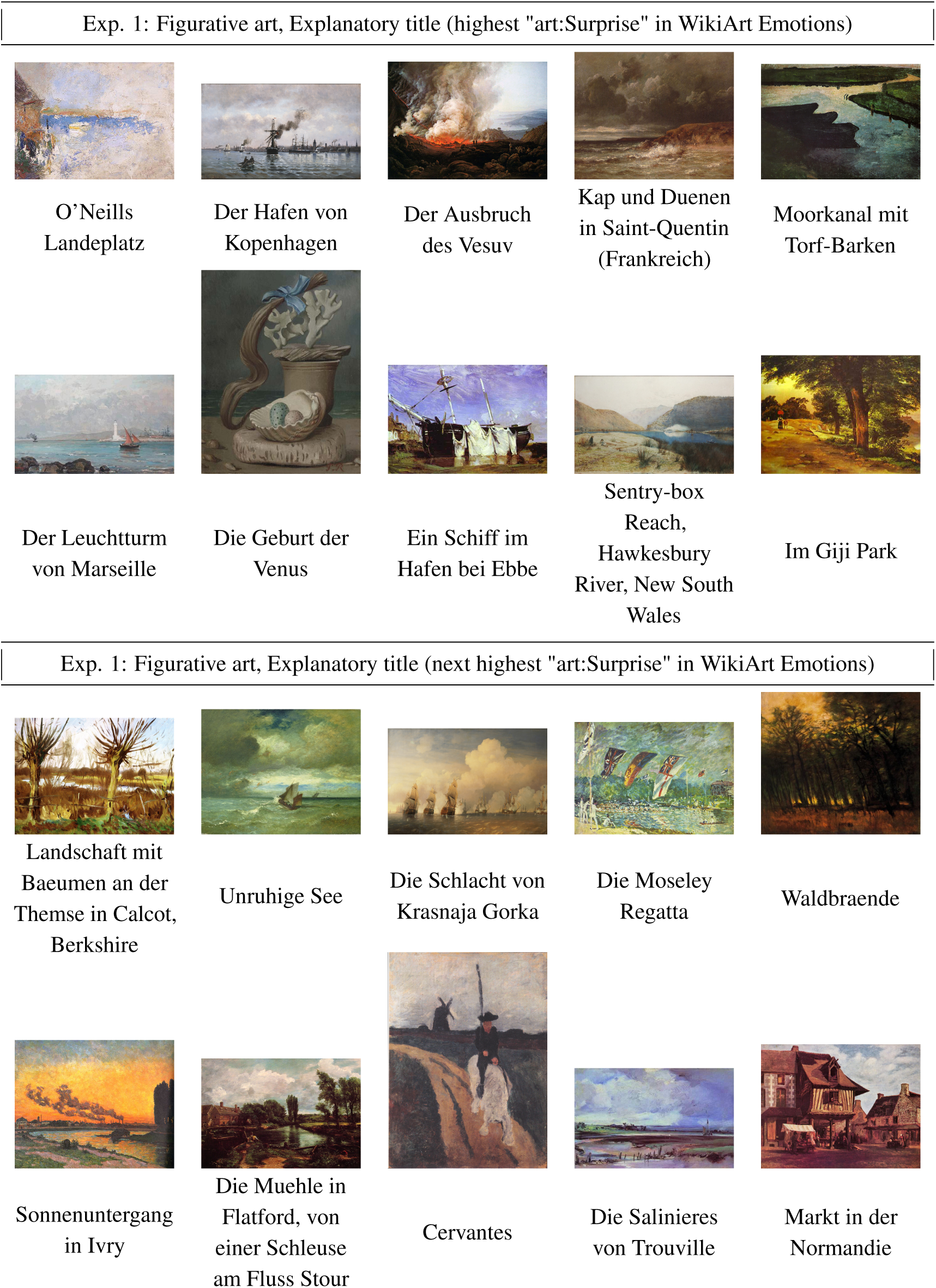
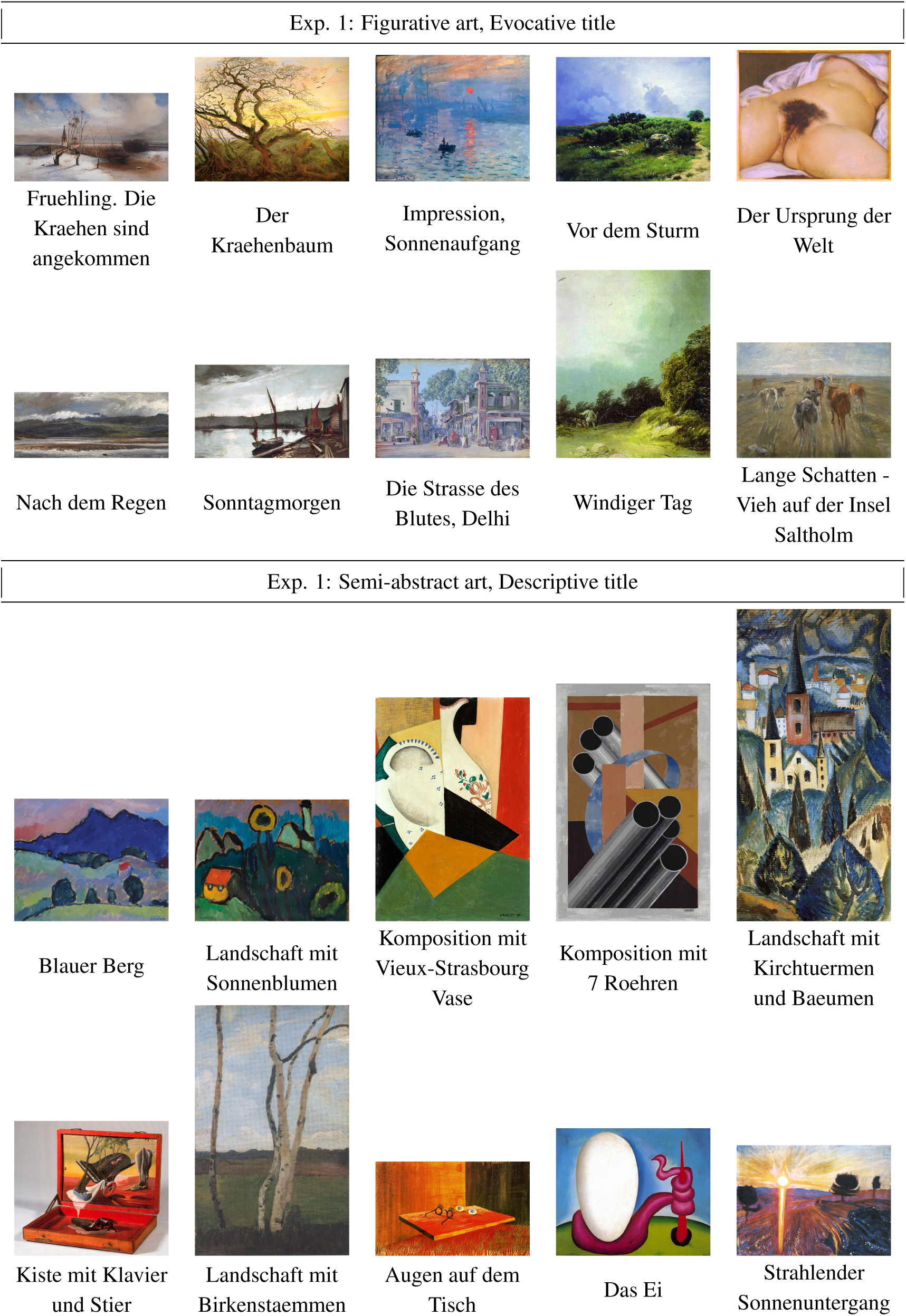
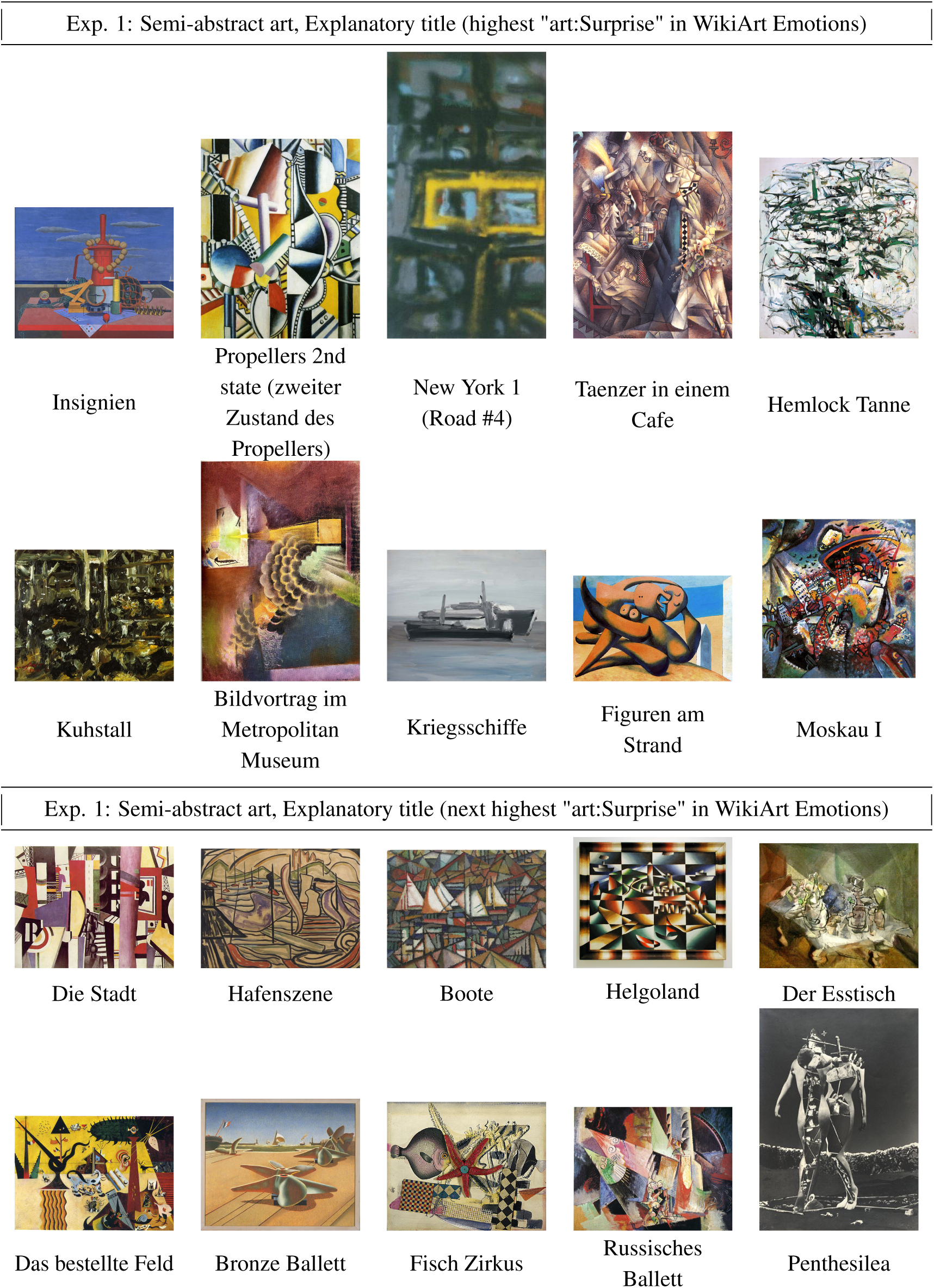
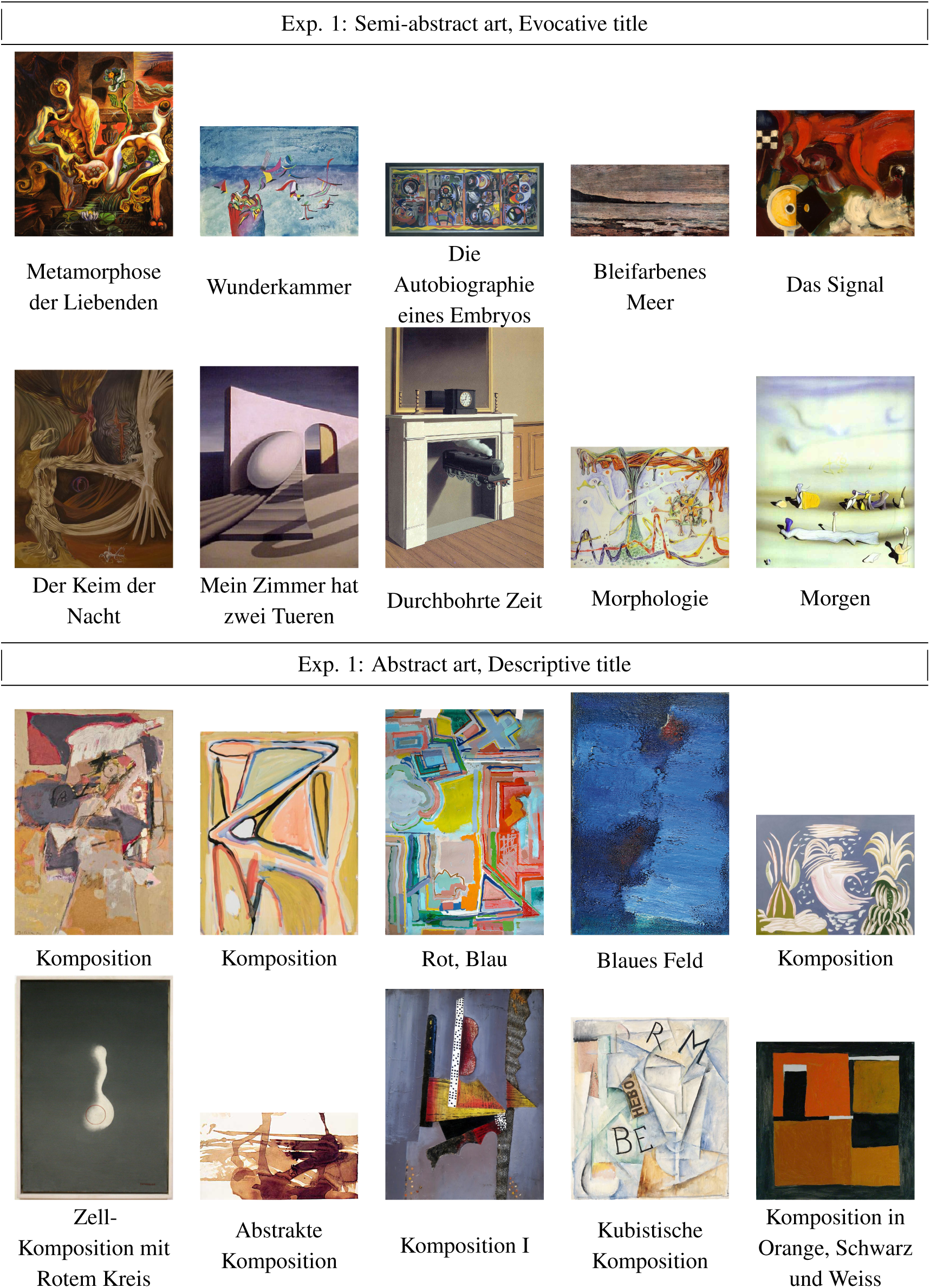
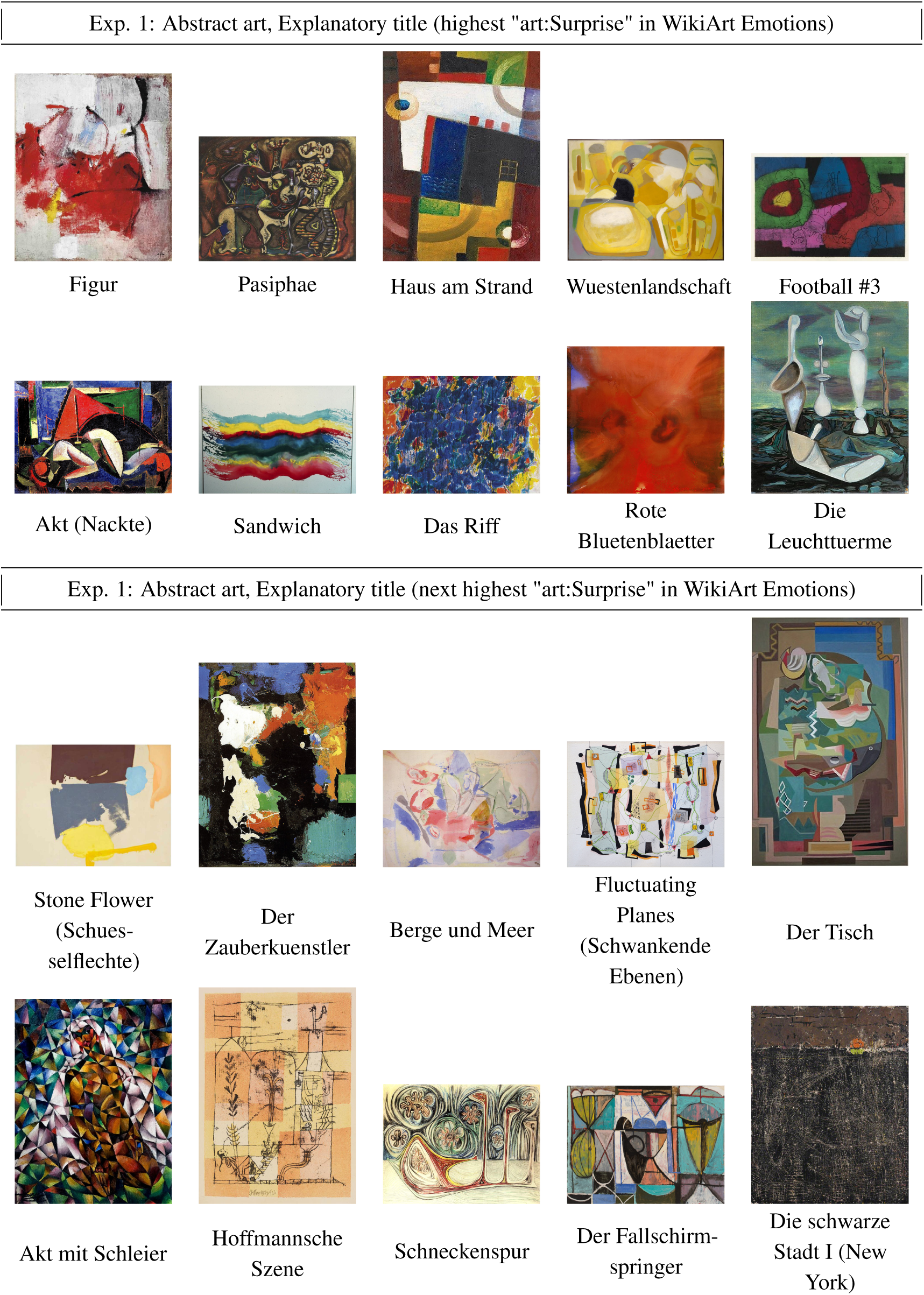
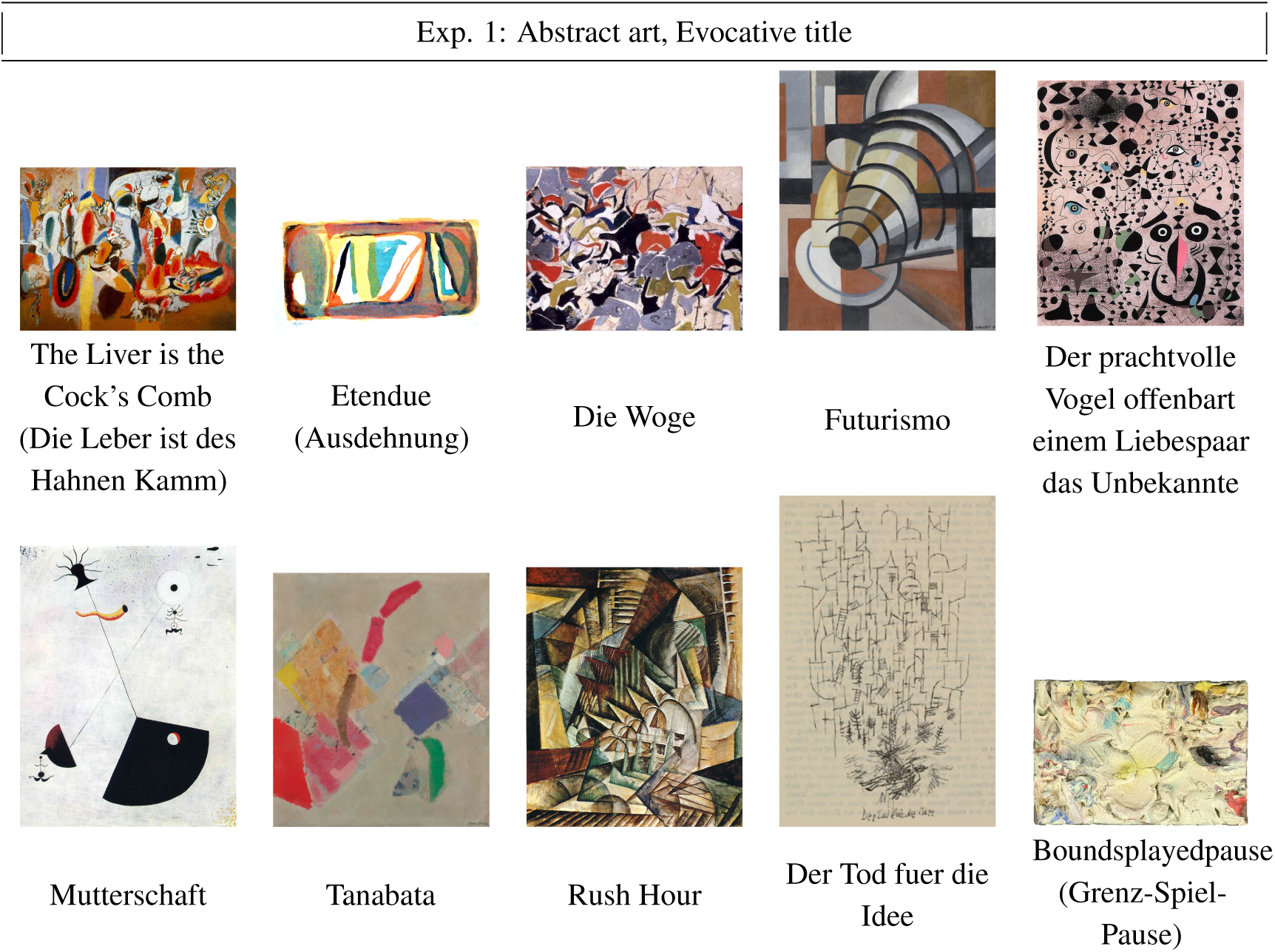

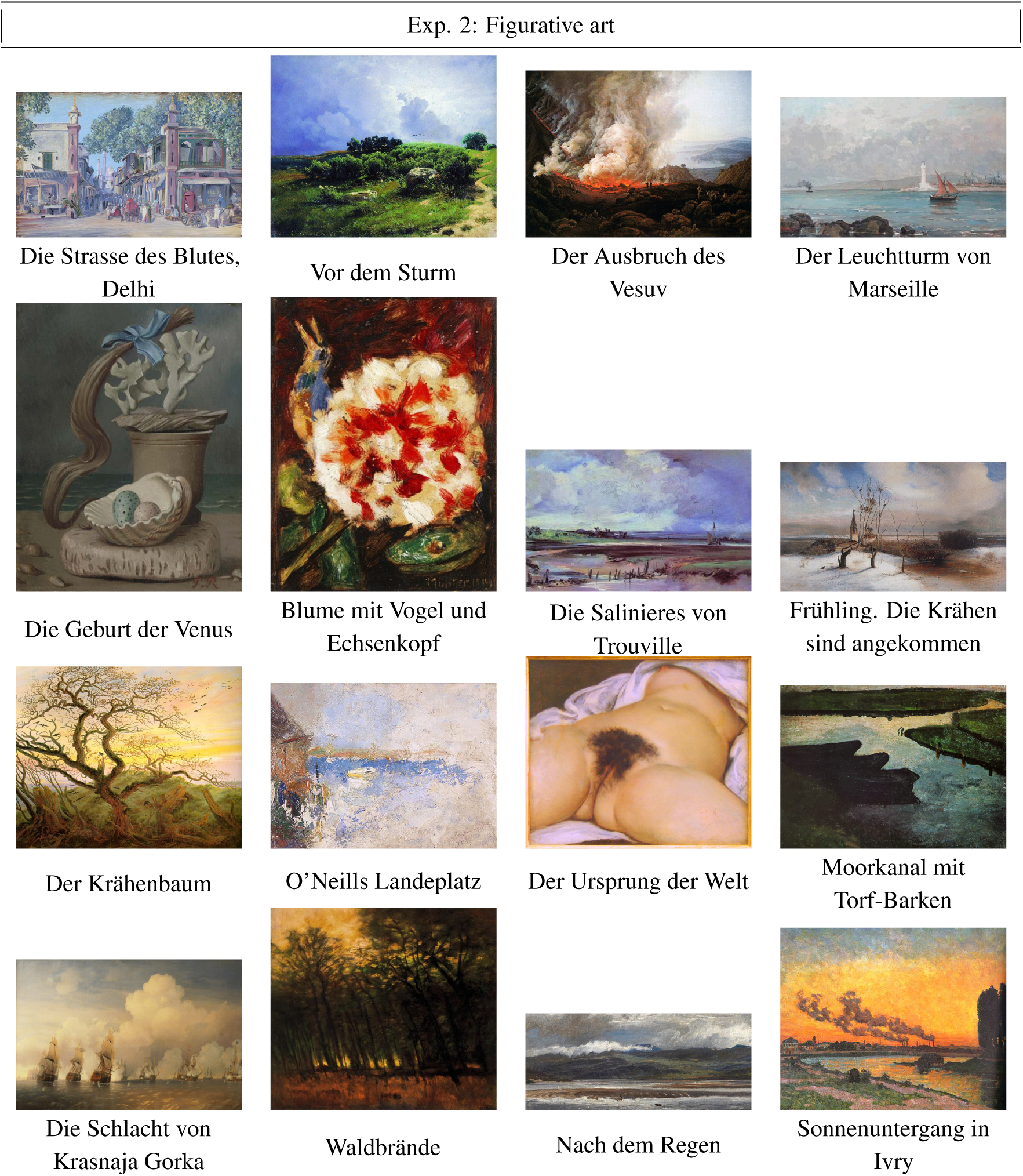
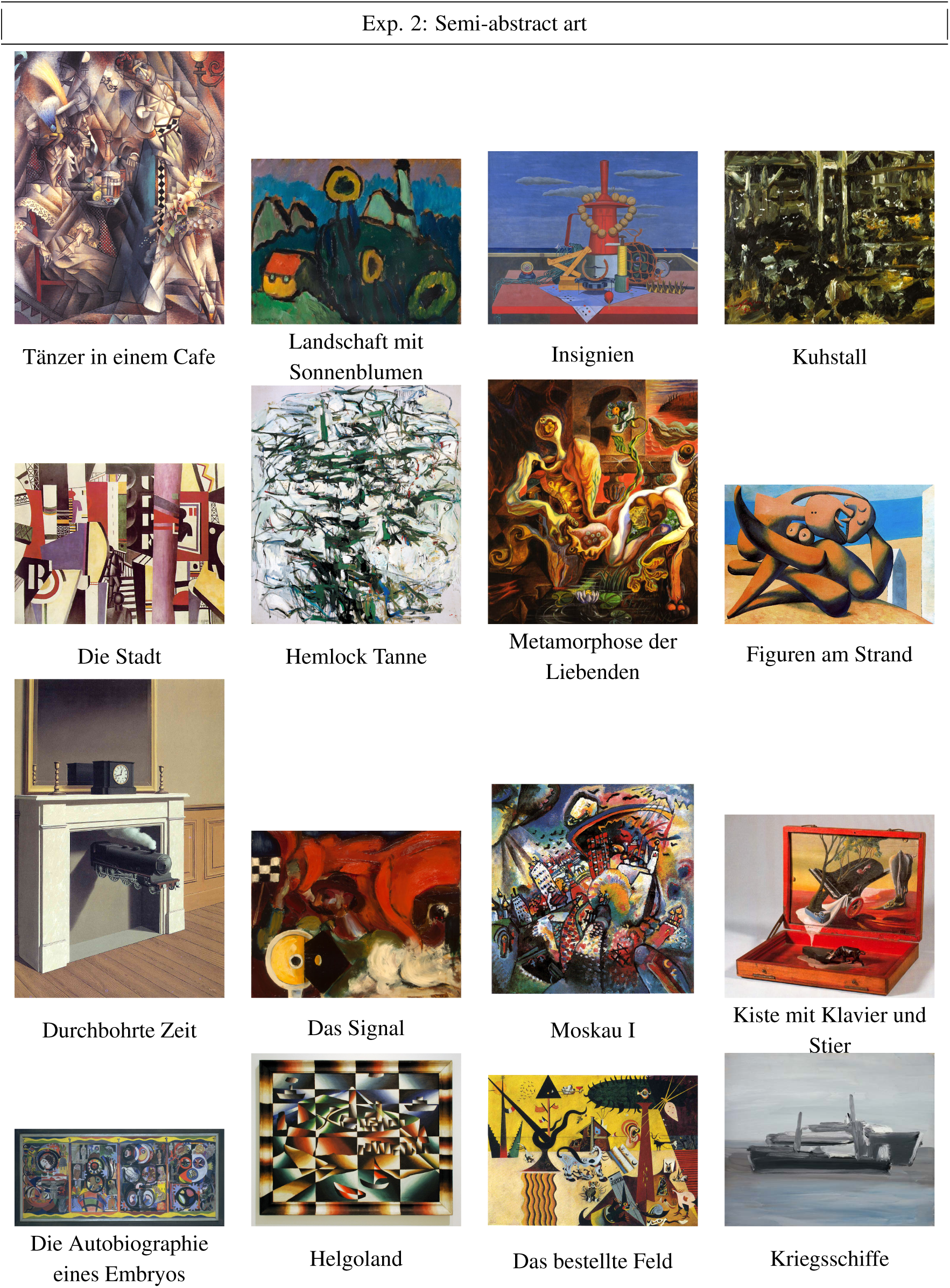
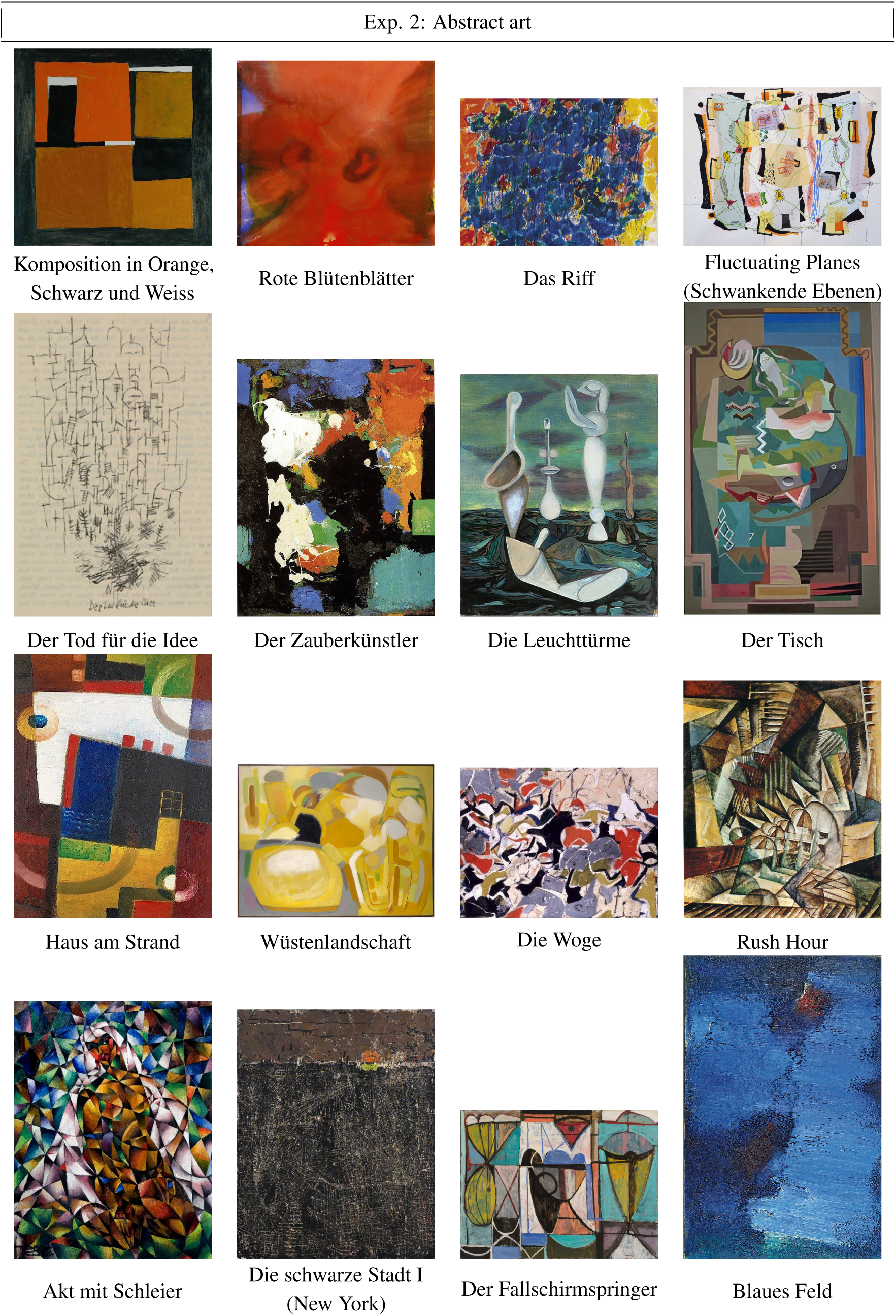

### A.2 Additional results & Discussion

#### A.2.1 Correlation of trial wise ratings

Repeated measures correlation (Fig. A.2) and paired local regression (Fig. A.3) of all collected ratings (data from Exp. 1). Repeated measures correlation (Bakdash & Marusich, 2017) implemented in the pingouin python package (Vallat, 2018, version 0.5.5).

**Figure A.2:**
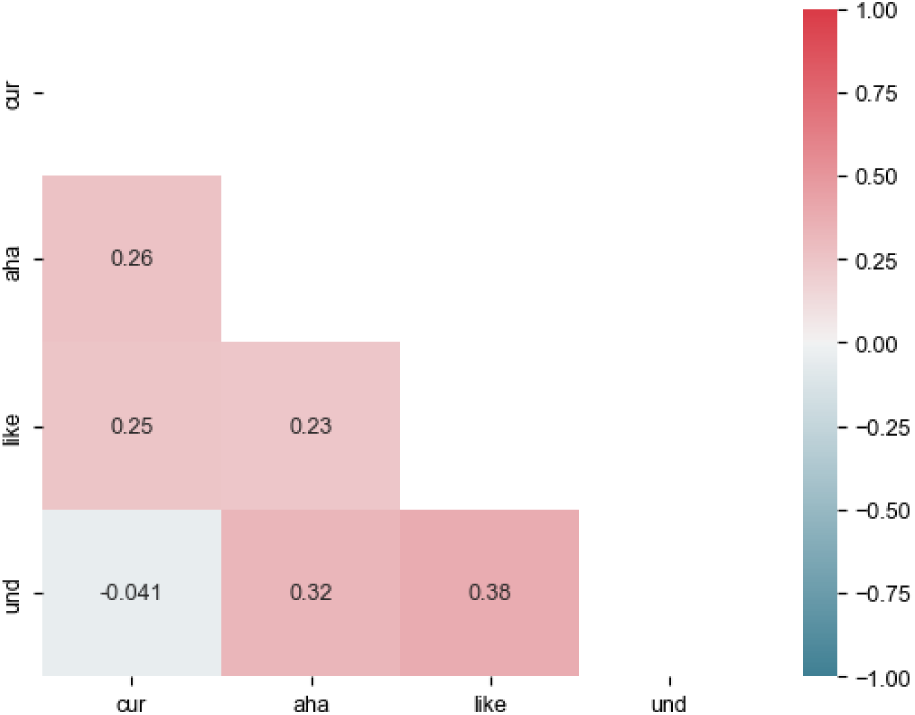
Correlation of ratings. Repeated measures correlation; values indicate R; all *p < .*05 (data from Exp. 1)

#### A.2.2 Full LMM details for Exp. 1

Tabs. Supp. A.3–A.18 report full model specification for all LMMs from the results.

**Table A.3:** *understanding* ∼ *titlecondition* + (*titlecondition*|*participant*) + (*titlecondition*|*picture*) (Eq. 1)

| Predictors | rating_understanding |  |  |
| --- | --- | --- | --- |
|  | Estimates | CI | p |
| title condition [untitled] (Intercept) | 45.87 | 41.32–50.42 | **<0.001** |
| title condition [titled] | 11.06 | 8.11–14.02 | **<0.001** |
| **Random Effects** |  |  |  |
| $\sigma^2$ | 520.53 | | |
| $\tau_{00}$ (stim_file) | 232.70 | | |
| $\tau_{00}$ (sub_id) | 169.85 | | |
| $\tau_{11}$ (stim_file.title_conditiontitle) | 34.93 | | |
| $\tau_{11}$ (sub_id.title_conditiontitle) | 65.47 | | |
| $\rho_{01}$ (stim_file) | -0.18 | | |
| $\rho_{01}$ (sub_id) | -0.51 | | |
| ICC | 0.42 |  |  |
| N (sub_id) | 56 |  |  |
| N (stim_file) | 120 |  |  |
| Observations | 2647 |  |  |
| Marginal $R^2$ / Conditional $R^2$ | 0.033 / 0.443 | | |

**Table A.4:**
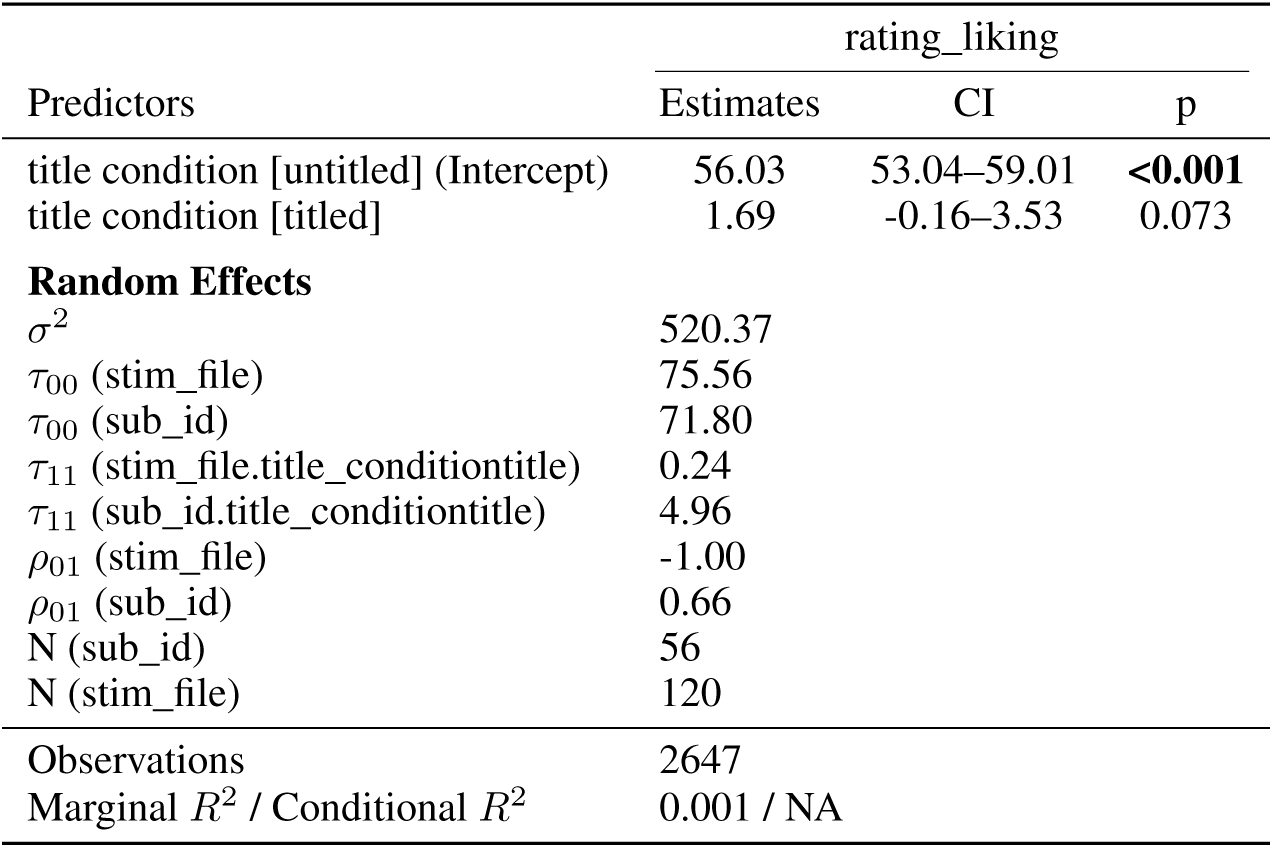
*liking* ∼ *titlecondition* + (*titlecondition*|*participant*) + (*titlecondition*|*picture*) (Eq. 2)

**Table A.5:** *aha* ∼ *titlecondition* + (*titlecondition*|*participant*) + (*titlecondition*|*picture*) (Eq. 3)

| Predictors | rating_aha |  |  |
| --- | --- | --- | --- |
|  | Estimates | CI | p |
| title condition [untitled] (Intercept) | 32.72 | 28.02–37.42 | **<0.001** |
| title condition [titled] | 17.89 | 13.80–21.97 | **<0.001** |
| **Random Effects** |  |  |  |
| $\sigma^2$ | 467.64 | | |
| $\tau_{00}$ (stim_file) | 3.05 | | |
| $\tau_{00}$ (sub_id) | 300.06 | | |
| $\tau_{11}$ (stim_file.title_conditiontitle) | 91.23 | | |
| $\tau_{11}$ (sub_id.title_conditiontitle) | 159.57 | | |
| $\rho_{01}$ (stim_file) | 1.00 | | |
| $\rho_{01}$ (sub_id) | -0.60 | | |
| ICC | 0.40 |  |  |
| N (sub_id) | 56 |  |  |
| N (stim_file) | 120 |  |  |
| Observations | 2647 |  |  |
| Marginal $R^2$ / Conditional $R^2$ | 0.093 / 0.457 | | |

**Figure A.3:**
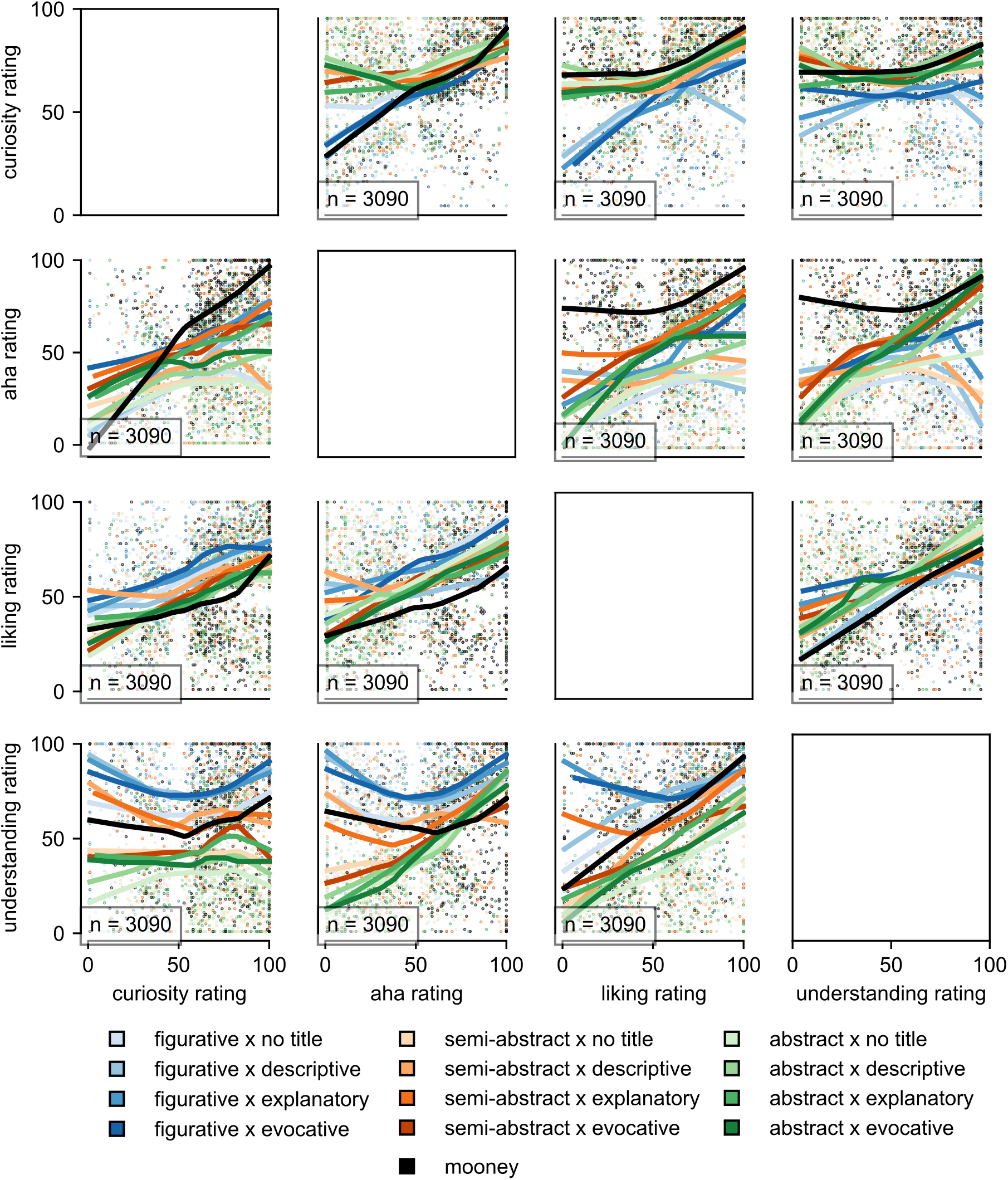
(Non)linear relation of ratings. Scatter plots with local regression (LOWESS) curves grouped by stimulus categories (data from Exp. 1)

**Table A.6:** *understanding* ∼ *titlecondition* + (*titlecondition*|*participant*) + (*1*|*picture*) (simpler alternative to Eq. 1)

| Predictors | rating_understanding |  |  |
| --- | --- | --- | --- |
|  | Estimates | CI | p |
| title condition [untitled] (Intercept) | 45.87 | 41.34–50.39 | **<0.001** |
| title condition [titled] | 11.08 | 8.31–13.85 | **<0.001** |
| **Random Effects** |  |  |  |
| $\sigma^2$ | 529.59 | | |
| $\tau_{00}$ (stim_file) | 224.70 | | |
| $\tau_{00}$ (sub_id) | 170.26 | | |
| $\tau_{11}$ (sub_id.title_conditiontitle) | 65.67 | | |
| $\rho_{01}$ (sub_id) | -0.51 | | |
| ICC | 0.41 |  |  |
| N (sub_id) | 56 |  |  |
| N (stim_file) | 120 |  |  |
| Observations | 2647 |  |  |
| Marginal $R^2$ / Conditional $R^2$ | 0.033 / 0.433 | | |

**Table A.7:** *liking* ∼ *titlecondition* + (*titlecondition*|*participant*) + (*1*|*picture*) (simpler alternative to Eq. 2)

| Predictors | rating_liking |  |  |
| --- | --- | --- | --- |
|  | Estimates | CI | p |
| title condition [untitled] (Intercept) | 56.03 | 53.07–58.99 | **<0.001** |
| title condition [titled] | 1.69 | -0.15–3.53 | 0.073 |
| **Random Effects** |  |  |  |
| $\sigma^2$ | 520.45 | | |
| $\tau_{00}$ (stim_file) | 71.34 | | |
| $\tau_{00}$ (sub_id) | 71.81 | | |
| $\tau_{11}$ (sub_id.title_conditiontitle) | 4.96 | | |
| $\rho_{01}$ (sub_id) | 0.66 | | |
| ICC | 0.23 |  |  |
| N (sub_id) | 56 |  |  |
| N (stim_file) | 120 |  |  |
| Observations | 2647 |  |  |
| Marginal $R^2$ / Conditional $R^2$ | 0.001 / 0.234 | | |

**Table A.8:** *aha* ∼ *titlecondition* + (*titlecondition*|*participant*) + (*1*|*picture*) (simpler alternative to Eq. 3)

| Predictors | rating_aha |  |  |
| --- | --- | --- | --- |
|  | Estimates | CI | p |
| title condition [untitled] (Intercept) | 32.72 | 27.90–37.54 | **<0.001** |
| title condition [titled] | 17.94 | 14.18–21.71 | **<0.001** |
| **Random Effects** |  |  |  |
| $\sigma^2$ | 491.62 | | |
| $\tau_{00}$ (stim_file) | 40.55 | | |
| $\tau_{00}$ (sub_id) | 297.67 | | |
| $\tau_{11}$ (sub_id.title_conditiontitle) | 163.13 | | |
| $\rho_{01}$ (sub_id) | -0.60 | | |
| ICC | 0.37 |  |  |
| N (sub_id) | 56 |  |  |
| N (stim_file) | 120 |  |  |
| Observations | 2647 |  |  |
| Marginal $R^2$ / Conditional $R^2$ | 0.094 / 0.428 | | |

**Table A.9:**
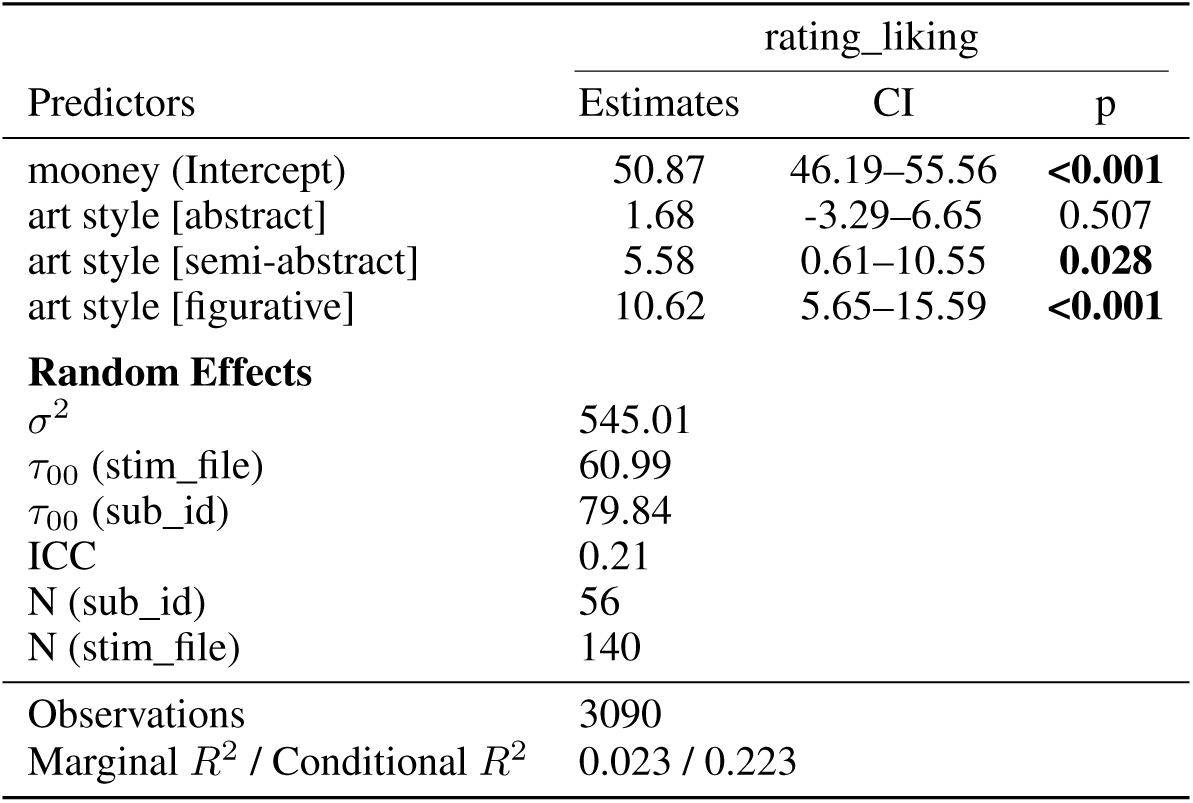
*liking* ∼ *artstyle* + (*1*|*participant*) + (*1*|*picture*) (Eq. 4)

| Predictors | rating_liking |  |  |
| --- | --- | --- | --- |
|  | Estimates | CI | p |
| mooney (Intercept) | 50.87 | 46.19–55.56 | **<0.001** |
| art style [abstract] | 1.68 | -3.29–6.65 | 0.507 |
| art style [semi-abstract] | 5.58 | 0.61–10.55 | **0.028** |
| art style [figurative] | 10.62 | 5.65–15.59 | **<0.001** |
| **Random Effects** |  |  |  |
| $\sigma^2$ | 545.01 | | |
| $\tau_{00}$ (stim_file) | 60.99 | | |
| $\tau_{00}$ (sub_id) | 79.84 | | |
| ICC | 0.21 |  |  |
| N (sub_id) | 56 |  |  |
| N (stim_file) | 140 |  |  |
| Observations | 3090 |  |  |
| Marginal $R^2$ / Conditional $R^2$ | 0.023 / 0.223 | | |

**Table A.10:** *aha* ∼ *artstyle : titlestyle* + (*1*|*participant*) + (*1*|*picture*) (Eq. 5)

| Predictors | rating_aha |  |  |
| --- | --- | --- | --- |
|  | Estimates | CI | p |
| mooney (Intercept) | 71.27 | 66.20–76.33 | <0.001 |
| art style [abstract] × title style [evocative] | -24.39 | -32.17– -16.61 | <0.001 |
| art style [semi-abstract] × title style [evocative] | -15.68 | -23.47– -7.90 | <0.001 |
| art style [figurative] × title style [evocative] | -14.69 | -22.45– -6.92 | <0.001 |
| art style [abstract] × title style [explanatory] | -18.26 | -24.46– -12.06 | <0.001 |
| art style [semi-abstract] × title style [explanatory] | -11.50 | -17.71– -5.29 | <0.001 |
| art style [figurative] × title style [explanatory] | -19.24 | -25.44– -13.03 | <0.001 |
| art style [abstract] × title style [descriptive] | -29.92 | -37.68– -22.16 | <0.001 |
| art style [semi-abstract] × title style [descriptive] | -28.95 | -36.73– -21.18 | <0.001 |
| art style [figurative] × title style [descriptive] | -32.05 | -39.82– -24.29 | <0.001 |
| **Random Effects** |  |  |  |
| $\sigma^2$ | 617.53 | | |
| $\tau_{00}$ (stim_file) | 57.86 | | |
| $\tau_{00}$ (sub_id) | 132.98 | | |
| ICC | 0.24 |  |  |
| N (sub_id) | 56 |  |  |
| N (stim_file) | 140 |  |  |
| Observations | 1764 |  |  |
| Marginal $R^2$ / Conditional $R^2$ | 0.121 / 0.329 | | |

**Table A.11:**
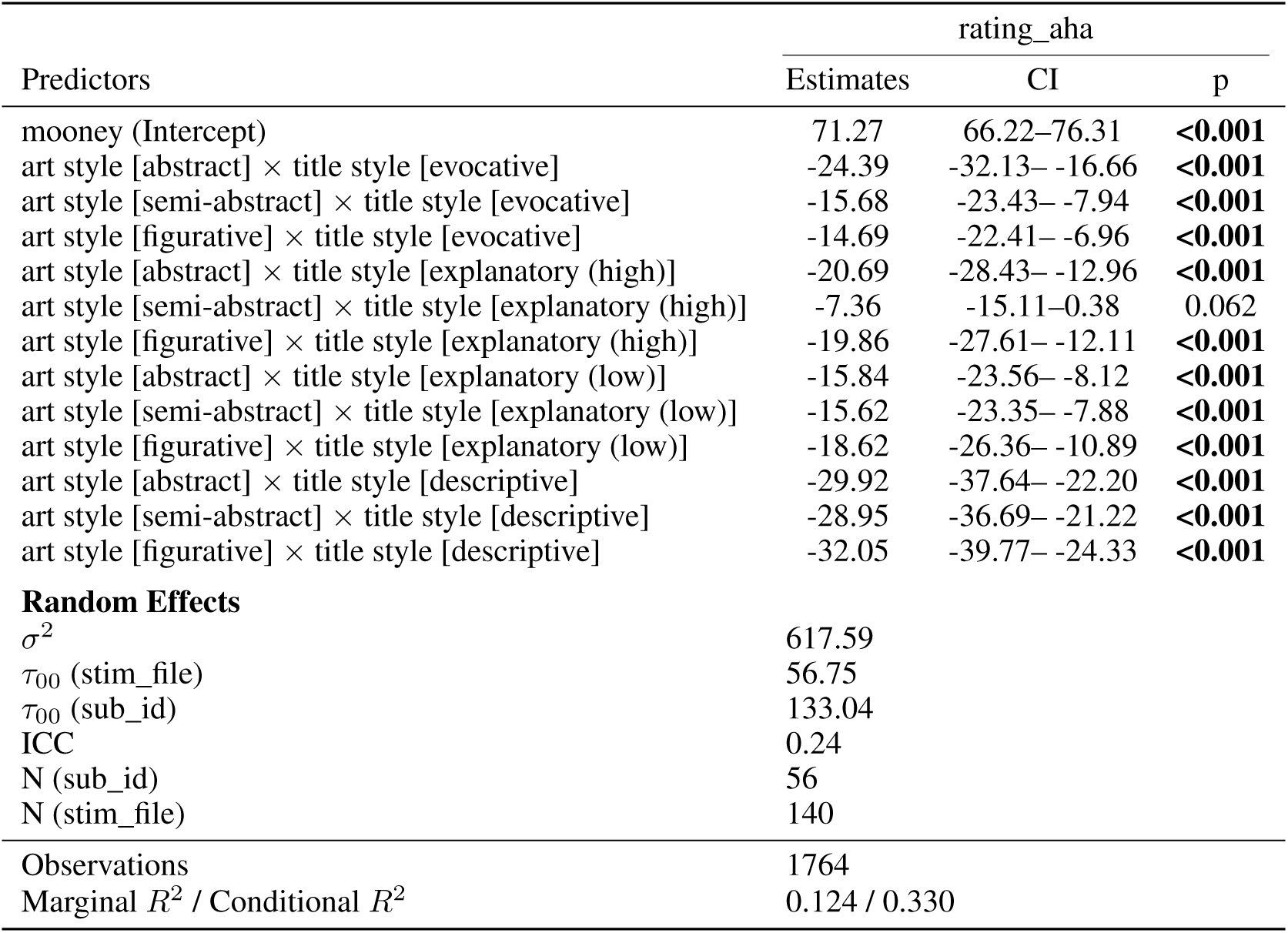
*aha* ∼ *artstyle* : *titlestyle* +(1|*participant*)+(1|*picture*) (Eq. 5) – resolving explanatory title category into subgroups (see Methods)

**Table A.12:** *aha* ∼ *curiosity : titlecondition* + (*1*|*participant*) + (*1*|*picture*) (Eq. 6)

| Predictors | rating_aha |  |  |
| --- | --- | --- | --- |
|  | Estimates | CI | p |
| (Intercept) | 26.23 | 21.98–30.48 | <0.001 |
| rating curiosity × title condition [untitled] | 0.11 | 0.07–0.15 | <0.001 |
| rating curiosity × title condition [titled] | 0.37 | 0.32–0.41 | <0.001 |
| rating curiosity × title condition [mooney] | 0.63 | 0.58–0.69 | <0.001 |
| **Random Effects** |  |  |  |
| $\sigma^2$ | 522.00 | | |
| $\tau_{00}$ (stim_file) | 34.82 | | |
| $\tau_{00}$ (sub_id) | 138.21 | | |
| ICC | 0.25 |  |  |
| N (sub_id) | 56 |  |  |
| N (stim_file) | 140 |  |  |
| Observations | 3090 |  |  |
| Marginal $R^2$ / Conditional $R^2$ | 0.257 / 0.442 | | |

**Table A.13:** *aha* ∼ *curiosity : titlestyle* + (*1*|*participant*) + (*1*|*picture*) (Eq. 8)

| Predictors | rating_aha |  |  |
| --- | --- | --- | --- |
|  | Estimates | CI | p |
| (Intercept) | 26.09 | 21.89–30.28 | <0.001 |
| rating curiosity × title style [mooney] | 0.63 | 0.58–0.69 | <0.001 |
| rating curiosity × title style [evocative] | 0.38 | 0.32–0.43 | <0.001 |
| rating curiosity × title style [explanatory (high)] | 0.43 | 0.37–0.48 | <0.001 |
| rating curiosity × title style [explanatory (low)] | 0.42 | 0.37–0.48 | <0.001 |
| rating curiosity × title style [descriptive] | 0.25 | 0.19–0.30 | <0.001 |
| rating curiosity × title style [untitled] | 0.11 | 0.07–0.15 | <0.001 |
| **Random Effects** |  |  |  |
| $\sigma^2$ | 518.24 | | |
| $\tau_{00}$ (stim_file) | 26.12 | | |
| $\tau_{00}$ (sub_id) | 136.77 | | |
| ICC | 0.24 |  |  |
| N (sub_id) | 56 |  |  |
| N (stim_file) | 140 |  |  |
| Observations | 3090 |  |  |
| Marginal $R^2$ / Conditional $R^2$ | 0.270 / 0.445 | | |

**Table A.14:**
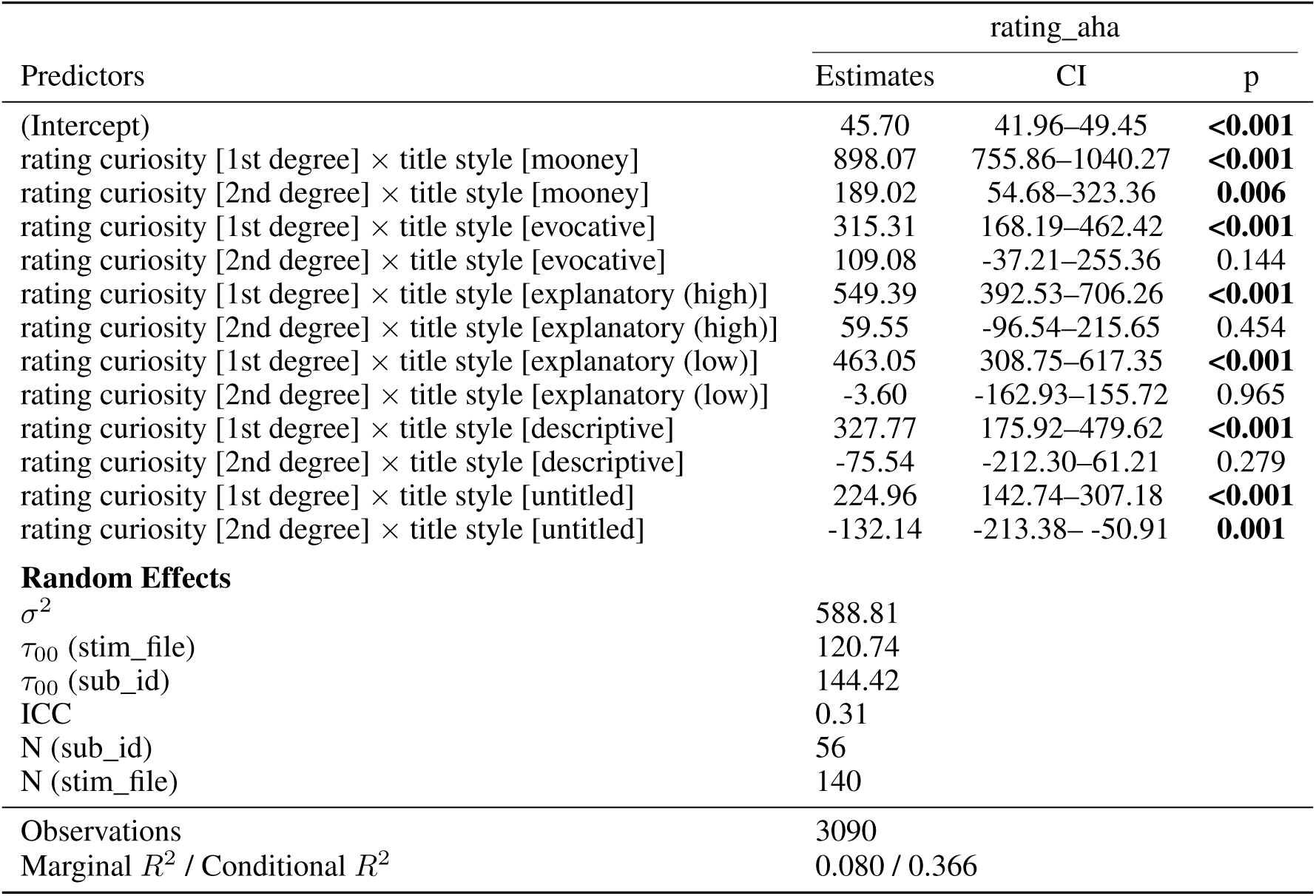
*aha* ∼ *quadratic*(*curiosity*) : *titlestyle* + (*1*|*participant*) + (*1*|*picture*) (Eq. 9)

| Predictors | rating_aha |  |  |
| --- | --- | --- | --- |
|  | Estimates | CI | p |
| (Intercept) | 45.70 | 41.96–49.45 | **<0.001** |
| rating curiosity [1st degree] × title style [mooney] | 898.07 | 755.86–1040.27 | **<0.001** |
| rating curiosity [2nd degree] × title style [mooney] | 189.02 | 54.68–323.36 | **0.006** |
| rating curiosity [1st degree] × title style [evocative] | 315.31 | 168.19–462.42 | **<0.001** |
| rating curiosity [2nd degree] × title style [evocative] | 109.08 | -37.21–255.36 | 0.144 |
| rating curiosity [1st degree] × title style [explanatory (high)] | 549.39 | 392.53–706.26 | **<0.001** |
| rating curiosity [2nd degree] × title style [explanatory (high)] | 59.55 | -96.54–215.65 | 0.454 |
| rating curiosity [1st degree] × title style [explanatory (low)] | 463.05 | 308.75–617.35 | **<0.001** |
| rating curiosity [2nd degree] × title style [explanatory (low)] | -3.60 | -162.93–155.72 | 0.965 |
| rating curiosity [1st degree] × title style [descriptive] | 327.77 | 175.92–479.62 | **<0.001** |
| rating curiosity [2nd degree] × title style [descriptive] | -75.54 | -212.30–61.21 | 0.279 |
| rating curiosity [1st degree] × title style [untitled] | 224.96 | 142.74–307.18 | **<0.001** |
| rating curiosity [2nd degree] × title style [untitled] | -132.14 | -213.38– -50.91 | **0.001** |
| **Random Effects** |  |  |  |
| $\sigma^2$ | 588.81 | | |
| $\tau_{00}$ (stim_file) | 120.74 | | |
| $\tau_{00}$ (sub_id) | 144.42 | | |
| ICC | 0.31 |  |  |
| N (sub_id) | 56 |  |  |
| N (stim_file) | 140 |  |  |
| Observations | 3090 |  |  |
| Marginal $R^2$ / Conditional $R^2$ | 0.080 / 0.366 | | |

**Table A.15:** *liking* ∼ *aha : artstyle* + (*1*|*participant*) + (*1*|*picture*) (Eq. 10)

| Predictors | rating_liking |  |  |
| --- | --- | --- | --- |
|  | Estimates | CI | p |
| (Intercept) | 44.74 | 41.77–47.71 | **<0.001** |
| rating aha × art style [mooney] | 0.15 | 0.10–0.20 | **<0.001** |
| rating aha × art style [abstract] | 0.26 | 0.21–0.31 | **<0.001** |
| rating aha × art style [semi-abstract] | 0.24 | 0.19–0.29 | **<0.001** |
| rating aha × art style [figurative] | 0.32 | 0.27–0.37 | **<0.001** |
| **Random Effects** |  |  |  |
| $\sigma^2$ | 506.10 | | |
| $\tau_{00}$ (stim_file) | 60.56 | | |
| $\tau_{00}$ (sub_id) | 65.70 | | |
| ICC | 0.20 |  |  |
| N (sub_id) | 56 |  |  |
| N (stim_file) | 140 |  |  |
| Observations | 3090 |  |  |
| Marginal $R^2$ / Conditional $R^2$ | 0.083 / 0.266 | | |

**Table A.16:** *liking* ∼ *quadratic*(*aha*) : *artstyle* + (*1*|*participant*) + (*1*|*picture*) (Eq. 11)

| Predictors | rating_liking |  |  |
| --- | --- | --- | --- |
|  | Estimates | CI | p |
| (Intercept) | 56.36 | 53.65–59.06 | **<0.001** |
| rating_aha [1st degree] × art style [mooney] | 244.62 | 87.16–402.07 | **0.002** |
| rating_aha [2nd degree] × art style [mooney] | 111.82 | -17.13–240.76 | 0.089 |
| rating_aha [1st degree] × art style [abstract] | 499.27 | 404.65–593.89 | **<0.001** |
| rating_aha [2nd degree] × art style [abstract] | -37.12 | -133.82–59.57 | 0.452 |
| rating_aha [1st degree] × art style [semi-abstract] | 372.49 | 281.78–463.20 | **<0.001** |
| rating_aha [2nd degree] × art style [semi-abstract] | 45.35 | -47.09–137.79 | 0.336 |
| rating_aha [1st degree] × art style [figurative] | 418.36 | 319.50–517.22 | **<0.001** |
| rating_aha [2nd degree] × art style [figurative] | 7.88 | -90.43–106.20 | 0.875 |
| **Random Effects** |  |  |  |
| $\sigma^2$ | 504.56 | | |
| $\tau_{00}$ (stim_file) | 77.79 | | |
| $\tau_{00}$ (sub_id) | 64.42 | | |
| ICC | 0.22 |  |  |
| N (sub_id) | 56 |  |  |
| N (stim_file) | 140 |  |  |
| Observations | 3090 |  |  |
| Marginal $R^2$ / Conditional $R^2$ | 0.079 / 0.282 | | |

**Table A.17:** *understanding* ∼ *aha : artstyle* + (*1*|*participant*) + (*1*|*picture*) (Eq. 12)

| Predictors | rating_understanding |  |  |
| --- | --- | --- | --- |
|  | Estimates | CI | p |
| (Intercept) | 37.94 | 33.33–42.55 | **<0.001** |
| rating_aha × art style [mooney] | 0.16 | 0.09–0.23 | **<0.001** |
| rating_aha × art style [abstract] | 0.48 | 0.42–0.53 | **<0.001** |
| rating_aha × art style [semi-abstract] | 0.38 | 0.33–0.44 | **<0.001** |
| rating_aha × art style [figurative] | 0.24 | 0.18–0.29 | **<0.001** |
| **Random Effects** |  |  |  |
| $\sigma^2$ | 502.79 | | |
| $\tau_{00}$ (stim_file) | 309.00 | | |
| $\tau_{00}$ (sub_id) | 144.60 | | |
| ICC | 0.47 |  |  |
| N (sub_id) | 56 |  |  |
| N (stim_file) | 140 |  |  |
| Observations | 3090 |  |  |
| Marginal $R^2$ / Conditional $R^2$ | 0.114 / 0.534 | | |

**Table A.18:** *understanding* ∼ *quadratic*(*aha*) : *artstyle* + (*1*|*participant*) + (*1*|*picture*) (Eq. 13)

| Predictors | rating_understanding |  |  |
| --- | --- | --- | --- |
|  | Estimates | CI | p |
| (Intercept) | 53.96 | 50.13–57.79 | **<0.001** |
| rating_aha [1st degree] × art style [mooney] | 60.79 | -101.97–223.56 | 0.464 |
| rating_aha [2nd degree] × art style [mooney] | 141.51 | 12.73–270.30 | **0.031** |
| rating_aha [1st degree] × art style [abstract] | 935.52 | 840.52–1030.52 | **<0.001** |
| rating_aha [2nd degree] × art style [abstract] | -19.48 | -116.45–77.48 | 0.694 |
| rating_aha [1st degree] × art style [semi-abstract] | 677.45 | 586.62–768.28 | **<0.001** |
| rating_aha [2nd degree] × art style [semi-abstract] | 96.42 | 3.87–188.98 | **0.041** |
| rating_aha [1st degree] × art style [figurative] | 304.85 | 205.74–403.96 | **<0.001** |
| rating_aha [2nd degree] × art style [figurative] | 216.92 | 118.40–315.44 | **<0.001** |
| **Random Effects** |  |  |  |
| $\sigma^2$ | 496.47 | | |
| $\tau_{00}$ (stim_file) | 179.21 | | |
| $\tau_{00}$ (sub_id) | 131.01 | | |
| ICC | 0.38 |  |  |
| N (sub_id) | 56 |  |  |
| N (stim_file) | 140 |  |  |
| Observations | 3090 |  |  |
| Marginal $R^2$ / Conditional $R^2$ | 0.135 / 0.468 | | |

#### A.2.3 Grand average physiological responses

Below we provide plots of the grand average data traces (only artwork trials, split by 1st or second presentation of each stimulus). While these plots do not map directly to the decoded data features (for these results see Fig. 3a) they might serve to give a general impression of the signal. In the case of ERP we see a prominent frontal/occipital gradient throughout pre and pottitle trials, that degrades over time. In the differential signal (subtraction done on the individual trial level) we see that the most prominent differences are in central vertex channels.

**Figure A.4:**
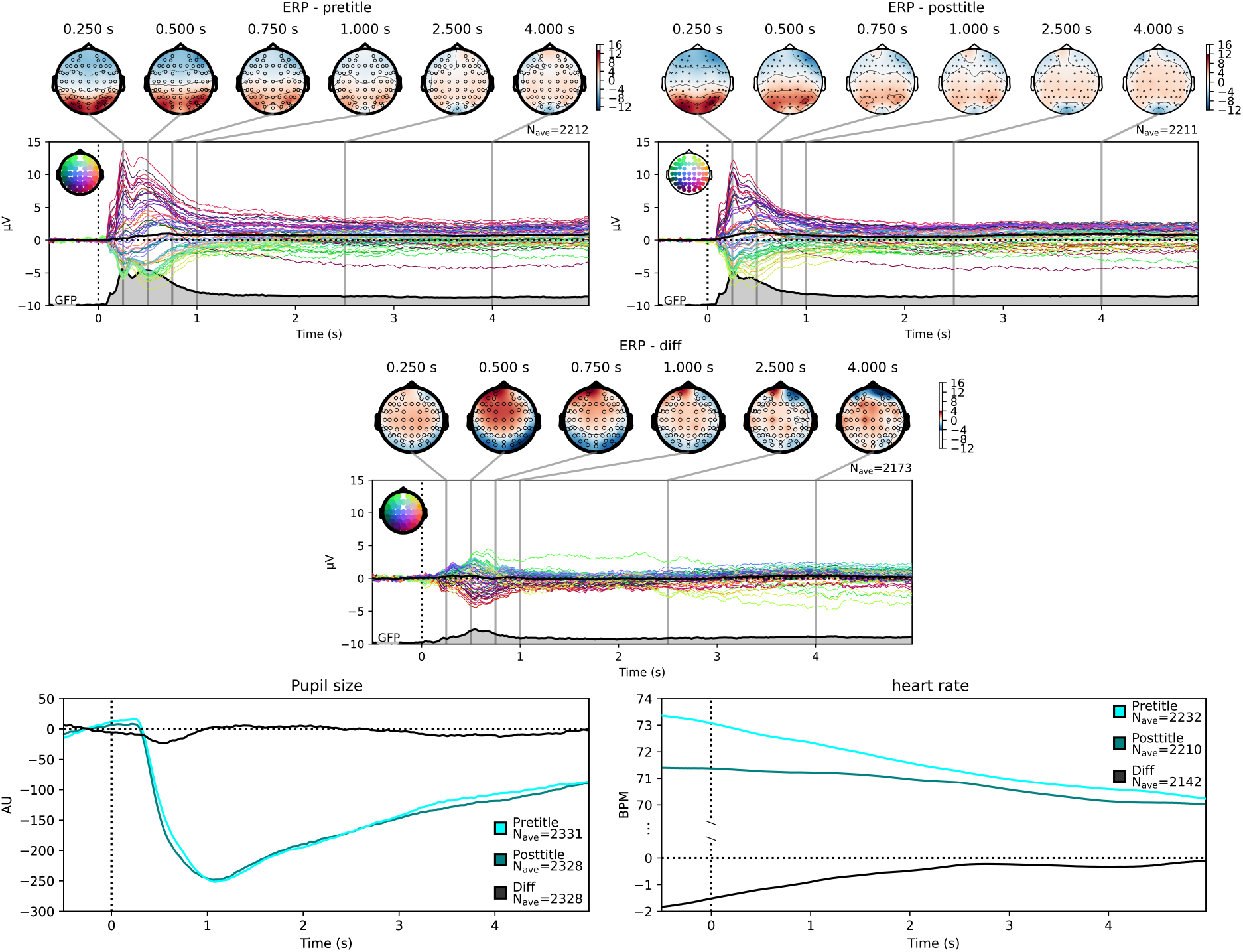
Grand average physiological responses in time domain signals – 1st vs 2nd presentation. depicted are grand average responses across all artwork trials, split by trial condition (pretitle – 1st presentation before title, posttitle – 2nd presentation after title, diff – trace during 2nd - trace during 1st presentation). Colored lines in EEG depict individual channels, black line the grand average, and at the bottom is the global field power (GFP). Above the time course are topography plots at 6 representative timepoints (selection not data driven). Pupil size and heart rate have only 2, respectively 1 data channel, hence the cleaner plot.

A similar comparison for the ratings is not straight forward, as these are continuous graded responses.

#### A.2.4 Decoding using a differential physiological signal

We repeated the decoding pipeline on the differential physiological responses (posttitle–pretitle traces). See Fig. Supp. A.8 for all decoding time-courses.

From this differential signal we could significantly decode the presented image category (Mooney image or painting) throughout the trial (starting 160 ms after image onset) in the time-domain EEG, and there were two short significant clusters in EEG alpha power (0.80-1.07 s and 4.24-4.44 s).

Title condition: time domain EEG - differential: significant 0.42–2.63 s after image onset

Insight: late differential pupil signal (2nd – 1st presentation; becoming significant from 4.18 s after image onset until the end of trial) There was further a short significant cluster in the differential theta-band power (2.83-2.99 s after image onset).

#### A.2.5 Full decoding results

**Figure A.5:**
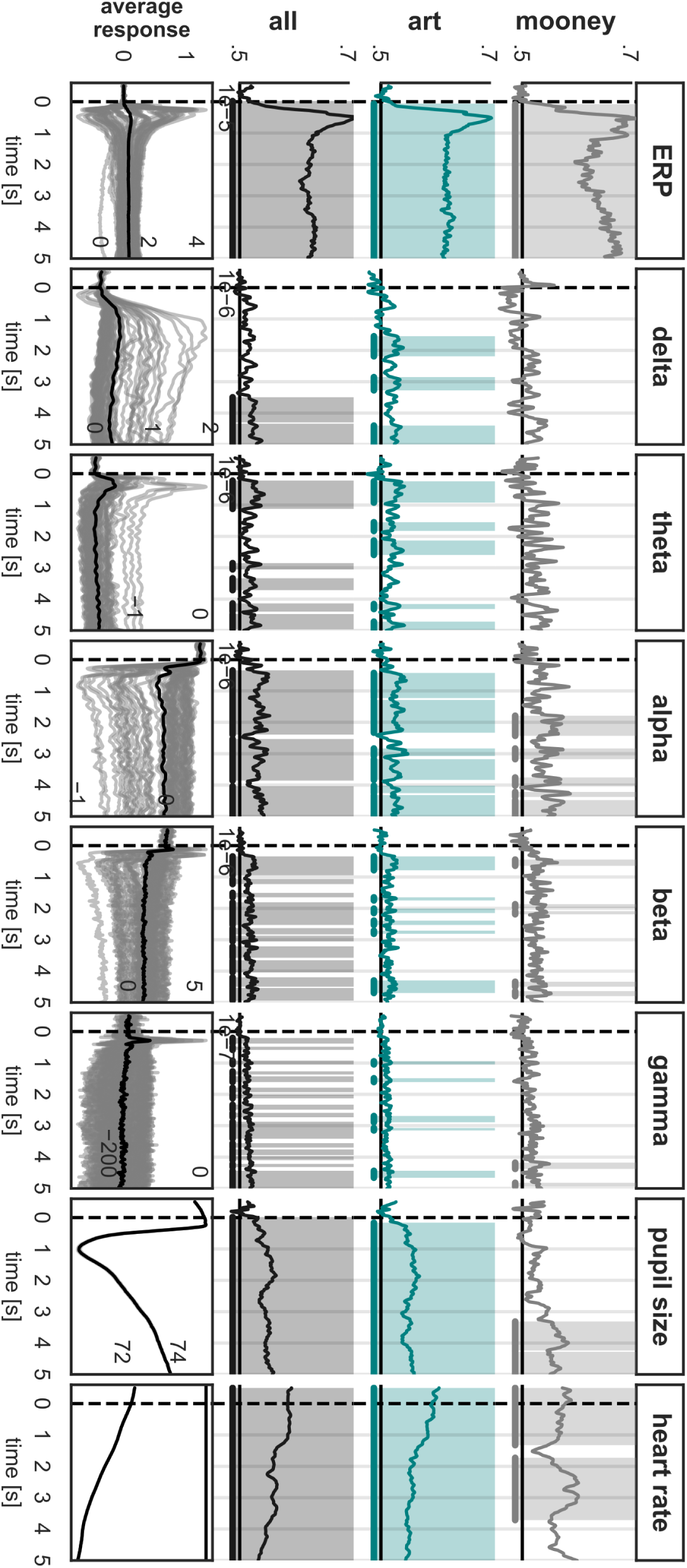
Full decoding results for 1st vs 2nd presentation of images decoding rate over time, significant timewindows are marked by color underlay (significance level *p < .*05 based on second level cluster permutation *T* -test).

**Figure A.6:**
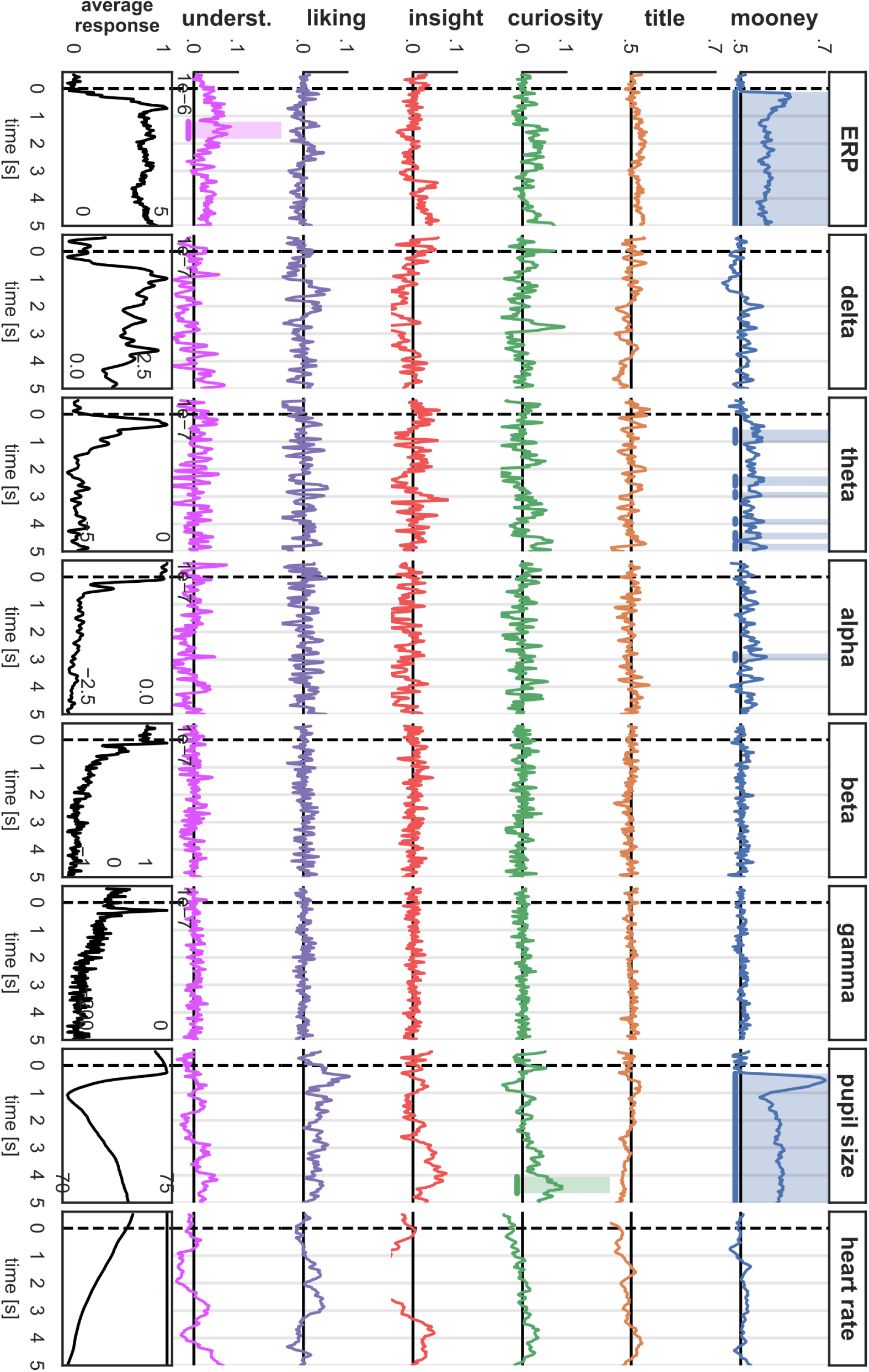
Full decoding results based on pretitle signal (1st presentation of images) decoding rate over time, significant timewindows are marked by color underlay (significance level *p < .*05 based on second level cluster permutation *T* -test).

**Figure A.7:**
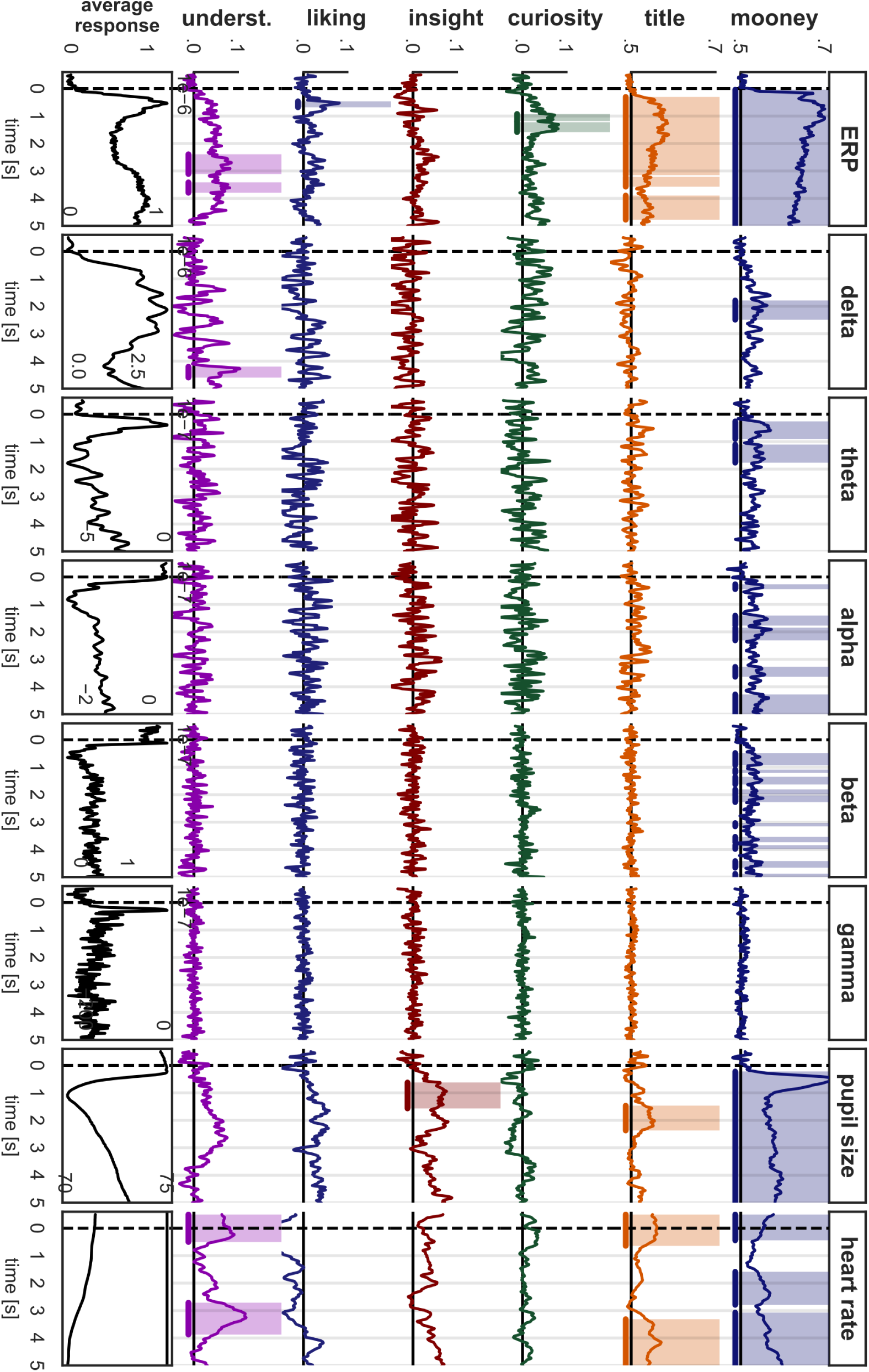
Full decoding results based on posttitle signal (2nd presentation of images) decoding rate over time, significant timewindows are marked by color underlay (significance level *p < .*05 based on second level cluster permutation *T* -test).

**Figure A.8:**
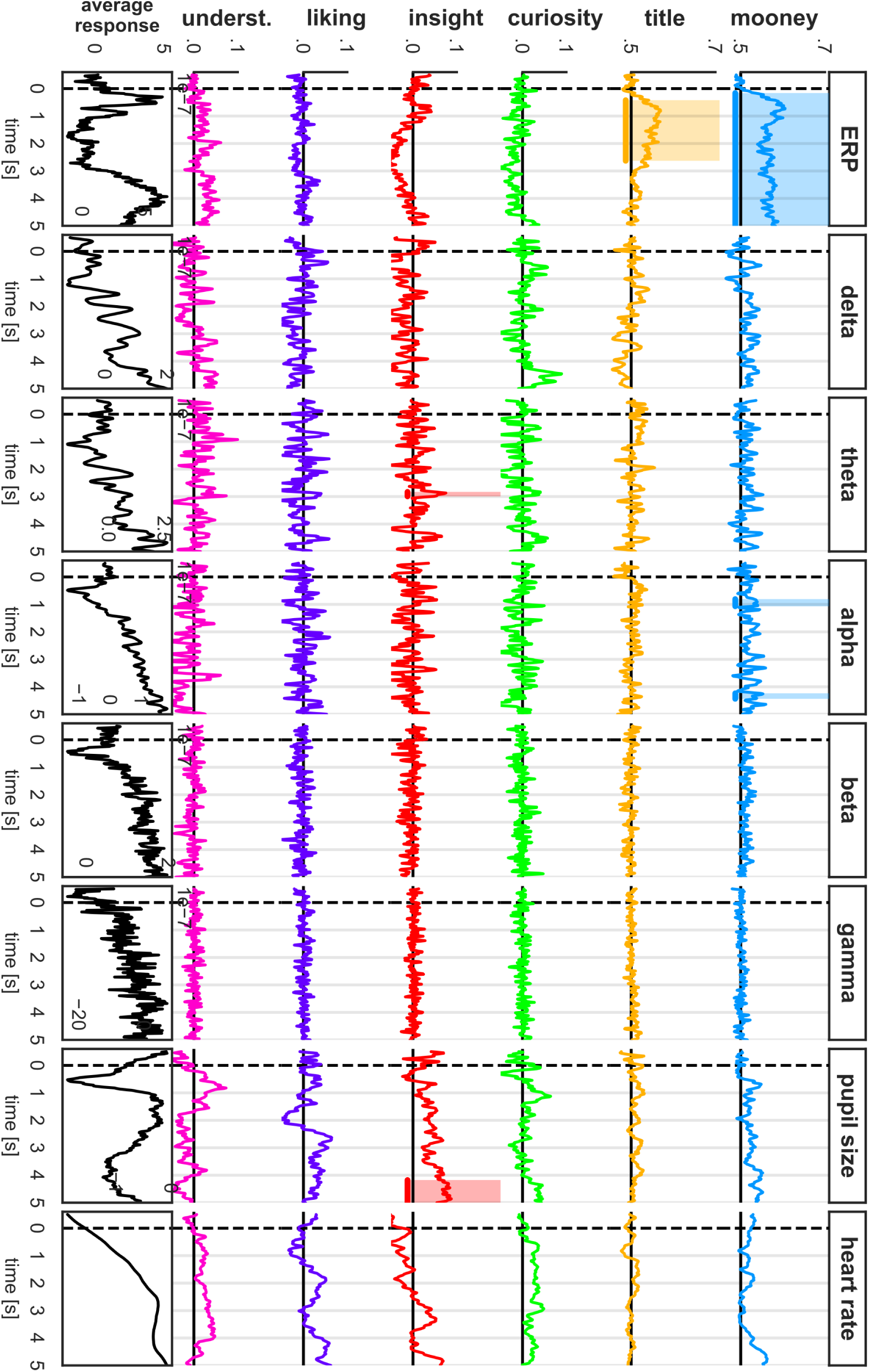
Full decoding results based on differential signal (2nd presentation – 1st presentation) decoding rate over time, significant timewindows are marked by color underlay (significance level *p < .*05 based on second level cluster permutation *T* -test).

#### A.2.6 Categorical decoding results

The categorical experimental manipulations we applied could be decoded quite robustly from the recorded data. Decoding worked very well for the contrast between the 1st and 2nd presentation of any given image stimulus (see Fig. 3a). Decoding was reliably possible from almost all data modalities. It was strongest from the time-domain EEG (significantly above chance throughout the full trial with a marked peak around 0.5–1 s after image onset, very high accuracy throughout). Time-frequency EEG in all frequency bands allowed above chance decoding during most of the trial duration, though not significant throughout. Alpha band EEG gave overall the best results, while decoding worked better for art stimuli than for Mooney images. There were no clear peaks, but in theta EEG decoding was better at the beginning of trials. The pupil trace allowed for significant above chance decoding throughout the full trial in art trials and all trials combined. When looking at Mooney trials alone, decoding became significant only at the end of the trial, starting around 3.3 s after image onset. The heart rate signal allowed significant decoding throughout the full trial – including the period before image onset – in art trials and all trials combined. When looking at Mooney trials alone, decoding was significant in the early (−0.5–1.33 s) and medium late trial (1.73–3.72 s). Looking at the grand average responses this does not come as a surprise (cf. Supp. Fig. A.4). Taken together this strongly suggests that the stimuli are not processed identically during repeated presentation; rather, cognitive processing differs in a reliably detectable way. Whether this is caused by mere repetition suppression (cf. Kim, 2017) or other factors specific to the study such as the interim contextualization with additional information (i.e. the real or fake title, or the Mooney solution respectively) or the differing implicit or explicit tasks (e.g., generating ratings) can not be clearly determined at this point.

The presented image category (Mooney image or painting) could also be reliably decoded right from the beginning from both EEG and pupil data (see Fig. 3b). In the time domain EEG data it could be clearly decoded during the first image presentation (pretitle: starting 120 ms after image onset), the second image presentation (posttitle: starting 40 ms after image onset). Decoding accuracy was well above chance throughout the full trial, but with a clear and marked peak at the beginning, about 1 s after image onset. Notably, classification worked more reliably during the second image presentation (i.e. higher decoding performances, and more sustained significance); this was the case for most of the investigated decoding tasks in this study. In the frequency domain EEG, findings were mixed. Theta and alpha band power were overall the best predictors; both delivered above chance decoding throughout the trials with several significant patches. Theta delivered good decoding in pretitle trials (significant: 0.56-1.05 s, 2.27-2.61 s, 2.83-3.03 s, 3.82-4.00 s, 4.32-4.56 s, and 4.72-4.94 s) and throughout the first 2 seconds of posttitle trials (significant: 0.26-0.91 s, and 1.11-1.77 s). Alpha was worse in pretitle trials (only significant: 2.79-3.03 s) but quite reliable throughout posttitle trials (significant: 0.26-0.44 s, 1.41-1.77 s, 1.85-2.31 s, 3.27-3.64 s, and 4.28-5.00 s). Beta band power was also a quite reliable predictor in posttitle trials (significant: 0.48-0.93 s, 1.09-1.21 s, 1.35-1.63 s, 1.81-1.99 s, 2.03-2.27 s, 3.03-3.15 s, 3.53-3.74 s, 3.84-3.98 s, 4.42-4.66 s, 4.88-5.00 s), but not in pretitle trials. Delta showed a significant blob in posttitle trials (1.79-2.49 s) but again not in pretitle trials. No significant results were found in the gamma band. In the pupil data the image category could be decoded during pretitle (starting at 300 ms) and posttitle presentation (starting 220 ms after image onset). As with ERPs, there was a clear peak at the beginning, around 1 s after image onset, then decreasing but staying well above chance for the rest of the trial. Interestingly, there was no big difference between decoding traces in pretitle and posttitle trials. Heart rate only predicted image category in the second presentation (significant from −0.5–0.44 s and from 1.59 s onward with a short non-significant intermission) but not in the pretitle condition. Besides the early peak in the decoding rate from pupil data (see discussion), single trial ERPs yielded the best results.

## Footnotes

1 The semi-abstract art with explanatory titles category seems to have even higher potential: we also fit a model that more strictly matches the study design (i.e. accounting for our oversampling of stimuli with explanatory titles, see Methods, by splitting these into two seperate title groups “explanatory (high)” and “explanatory (low)” ordered by “surprise” rating in the Wikiarts Emotion dataset). Here the semi abstract x explanatory (high) group had only marginally lower *aha* ratings than Mooney images (*slope* = *−*7.36, *CI* = [*−*15.11; *−*0.38], *p* = .062; see Tab. Supp. A.11

## Notes

https://osf.io/j42g5

https://osf.io/pagvh

https://osf.io/6kw9n

### Competing Interest Statement

The authors have declared no competing interest.

### Summary of Updates

revisions of the result reporting, especially for Exp. 1

